# Mitochondrial genome of non-photosynthetic mycoheterotrophic plant *Hypopitys monotropa*,its structure, gene expression and RNA editing

**DOI:** 10.1101/681718

**Authors:** Viktoria Y. Shtratnikova, Mikhail I. Schelkunov, Aleksey A. Penin, Maria D. Logacheva

## Abstract

Heterotrophic plants – the plants that lost the ability to photosynthesis – are characterized by a number of changes at all levels of organization. Heterotrophic plants divide into two large categories – parasitic and mycoheterotrophic. The question of to what extent these changes are similar in these two categories is still open. Plastid genomes of non-photosynthetic plants are well characterized and they demonstrate similar patterns of reduction in both groups. In contrast, little is known about mitochondrial genomes of mycoheterotrophic plants. We report the structure of the mitochondrial genome of *Hypopitys monotropa*, a mycoheterotrophic member of Ericaceae, and the expression of mitochondrial genes. In contrast to its highly reduced plastid genome, the mitochondrial genome of *H. monotropa* is larger than that of its photosynthetic relative *Vaccinium macrocarpon*, its complete size is ~810 Kbp. We found an unusually long repeat-rich structure of the genome that suggests the existence of linear fragments. Despite this unique feature, the gene content of the *H. monotropa* mitogenome is typical of flowering plants. No acceleration of substitution rates is observed in mitochondrial genes, in contrast to previous observations on parasitic non-photosynthetic plants. Transcriptome sequencing revealed trans-splicing of several genes and RNA editing in 33 genes of 38. Notably, we did not find any traces of horizontal gene transfer from fungi, in contrast to plant parasites which extensively integrate genetic material from their hosts.

## Introduction

*Hypopitys monotropa* (Ericacae, Monotropoideae) is a non-photosynthetic plant gaining carbon from fungi that are ectomycorrhizal with tree roots (Bjorkman, 1960). In contrast to most other mycoheterotrophic (MHT) plants, which are very rare and/or very narrowly distributed, Monotropoideae, including *H. monotropa*, are quite widespread, being associated with old-growth conifer forests. Thus, *H. monotropa* is used as a model system in studies of plant-mycorrhizal associations and developmental biology of MHT plants (e.g. Olson, 1993, 1990). Recent advances in DNA sequencing allow expanding the studies of mycoheterotrophs into genomics. By now, the most attention was focused on plastid genomes of MHT plants. They are highly reduced in size and gene content, includingcomplete absence or pseudogenization of genes of photosynthesis electron transport chain (for review see Graham et al., 2017). Thus, the MHT lifestyle strongly affects plastids, but what about the mitochondrial genome?

In contrast to animals, where mitochondrial genomes are usually conserved in size and gene content across large taxonomic groups, in plants they are highly variable and may be very dissimilar even in closely related species. The size of the angiosperm mitogenome is ranging from 66 Kb in the hemiparasitic mistletoe *Viscum scurruloideum* (Skippington et al., 2015), 222 Kb in the autotrophic *Brassica napus* (Handa, 2003) to more than 11 Mb in *Silene noctiflora*. Despite such huge variation in size, the large part of mitochondrial genes – the ones that encode the components of the oxidative phosphorylation chain complexes and proteins involved in the biogenesis of these complexes – are stable in content and have very low sequence divergence. More variation exists in the group of genes involved in translation – i.e. ribosomal proteins and transfer RNAs (Adams et al., 2002; Gualberto et al., 2014), presumably due to the transfer to the nuclear genome that occurred in several plant lineages or to the non-essentiality of these genes. Besides this, many plant mitochondrial genomes carry open reading frames (ORF) that potentially encode functional proteins (Qiu et al., 2014); such ORFs are highly lineage-specific. Non-photosynthetic plants divide into two large groups—those that are parasitic on other plants and mycoheterotropic. By now only few complete mitochondrial genomes of non-photosynthetic plants are characterized, and most of them are parasitic. A comparative analysis of mitogenomes of several parasitic, hemiparasitic and autotrophic Orobanchaceae (Fan et al., 2016) showed that the gene content does not depend on trophic specialization in family range. Mitogenomes of two non-related lineages of parasitic plants: Rafflesiaceae and Cynomoriaceae also do not show reduction in gene content; besides that, they are the example of massive HGT from other plants (including, but not only, their hosts). This is not, however, a trait unique for parasitic plants—e.g. *Amborella trichopoda*, an autotrophic plant from basal angiosperms, has in its mitochondrial genome a large fraction acquired from green algae, mosses, and other angiosperms (Rice et al., 2013). In contrast, in the hemiparasitic plant *Viscum scurruloideum* the mitogenome is drastically reduced in size and gene content, it lacks all nine *nad* genes, and *matR* (Skippington et al., 2015). Mitogenomes of other *Viscum* species are not reduced in length but have reduced gene content, similar to *V. scurruloideum* ((Petersen et al., 2015); but see (Skippington et al., 2017)). The sampling is obviously insufficient even for parasitic plants; regarding mycoheterotrophs the data are almost completely lacking. There are only two mitochondrial genomes of a MHT plants characterized by date - that of orchid *Gastrodia elata* (Yuan et al., 2018) and *Epirixanthes elongata* (Polygalaceae) (Petersen et al., 2019). The former one is characterized as a part of genome sequencing project and is not assembled completely being represented by 19 contigs. *G. elata* mitogenome is large (~ 1.3 Mb) while in *E. elongata* it is much smaller (~0,4 Mb). However, both above-mentioned studies lack comparative analysis with autotrophic relatives that would allow to highlight the changes associated with mycoheterotrophy.

Taking this into account, we set the following objectives: 1) to characterize the structure and gene content of the mitochondrial genome of *H. monotropa* 2) to estimate if horizontal gene transfer (HGT) from fungi took place 3) to study the mitochondrial gene expression and RNA editing.

The photosynthetic plant with characterized mitochondrial genome phylogenetically closest to *H. monotropa* is cranberry, *Vaccinium macrocarpon* (Fajardo et al., 2014). In this study we use *V. macrocarpon* as a basis for comparative analysis aimed at finding the patterns (if there are any) of mitochondrial genome changes associated with mycoheterotrophy.

## Materials & Methods

### 2.1. Sample collection and sequencing

Sample collection, DNA and RNA libraries preparation for most datasets (DNA shotgun and mate-pair libraries and inflorescence transcriptome library) and sequencing were described in in the papers of Logacheva et al. (2016) and Schelkunov et al. (2018). In addition to the data generated in previous studies we used four new datasets which represent transcriptomes of anthers and ovules of *H. monotropa*. The samples were collected in the same location as previous ones but in 2018. RNA was extracted using RNEasy kit (Qiagen) with the addition of Plant RNA isolation aid reagent (Thermo fisher) to the lysis buffer. Removal of ribosomal RNA was performed using Ribo-Zero plant leaf kit (Illumina) and further sample preparation using NEBNext RNA library preparation kit (New England Biolabs). The libraries were sequenced on HiSeq 4000 (Illumina) in 150-nt paired-end mode. The reads are deposited in NCBI Sequence Read Archive under the BioProject PRJNA573526.

### 2.2. Assembly

Read trimming and assembly were made as described in the paper of Logacheva et al. (2016). The contig coverages were determined by mapping all reads in CLC Genomics Workbench v. 7.5.1 (https://www.qiagenbioinformatics.com/products/clc-genomics-workbench/), requiring at least 80% of a read’s length to align with at least 98% sequence similarity. To find contigs corresponding to the mitochondrial genome, we performed BLASTN and TBLASTX alignment of *Vaccinium macrocarpon* genome (GenBank accession NC_023338) against all contigs in the assembly. BLASTN and TBLASTX were those of BLAST 2.3.0+ suite (Camacho et al., 2009), the alignment was performed with the maximum allowed *e*-value of 10^−5^. Low-complexity sequence filters were switched off in order not to miss genes with extremely high or low GC-content. A contig that corresponded to the plastid genome of *Hypopitys monotropa* also aligned to the mitochondrial genome of *Vaccinium macrocarpon* because of the presence of inserts from the plastid genome into the mitochondrial genome. The complete sequence of the *H. monotropa* plastome is known from our previous study (Logacheva et al., 2016) and was excluded from further analyses. Contigs that had coverage less than 10 or ones that produce matches only to non-mitochondrial sequences when aligned to NCBI NT database by BLASTN (*e*-value 10^−3^), were considered nuclear.

After the procedures described above, 7 contigs remained, with lengths 255, 5651, 20627, 102868, 106298, 234240, 309413 bp and coverages 37, 89, 93, 110, 96, 107, 108 respectively. The lower coverage of the smallest contig is presumably an artifact caused by its length. To understand their order in a mitochondrial genome, we mapped mate-pair reads from the library with the larger insert size (8705±2537 bp) to the contigs and investigated mate-pair links. The mapping was performed by CLC Genomics Workbench, 100% of a read’s length was required to align to a contig with a sequence similarity of 100%, to minimize the amount of incorrectly mapped reads. Mate-pair links were visualized by Circos 0.67-5 (Krzywinski et al., 2009) and investigated manually. In one contig mate-pair links spanned from its end to the start and thus the contig corresponds to a circular chromosome. Other six contigs connect into a linear sequence, with a structure described in the Results section. Gaps between the contigs were closed by GapFiller (Boetzer and Pirovano, 2012) that was run with parameters -m 30 -o 2 -r 0.8 -n 50 -d 100000 -t 100 -g 0 -T 4 -i 100000. The source code of GapFiller was rewritten to allow it to use Bowtie2 as the read mapper. Additionally, we tried extending the edges of the linear chromosome by GapFiller to check that there are no assembly forks (places with two alternative extensions). The absence of such forks indicates that the linearity of this sequence is not a consequence of an assembly problem. Instead, during the extension the coverage gradually decreases to zero, implying a linear chromosome with lengths varying between different copies of the chromosome. To check whether the two chromosomes have been assembled correctly, we mapped reads from all three sequencing libraries by CLC Genomics Workbench v. 7.5.1, requiring at least 90% of a read’s length to align with at least 99% sequence similarity and checked that the coverage is approximately uniform along the whole length of the chromosomes. Just as during the analysis by GapFiller, we found that the coverage gradually decreases on the edges of the linear chromosome, thus indicating that different copies of the chromosome in a plant have different lengths of the terminal regions. For definiteness, we elongated the terminal regions such that the coverage drops to 0, thus the length of the linear chromosome approximately corresponds to the maximal possible length among all copies of the chromosome in the sequenced plant. Also, we mapped reads from the mate-pair library with the larger insert size, requiring 100% of a read length to align with a sequence similarity of 100%, and investigated a distribution of mean insert sizes over each genomic position and a number of mate-pairs than span over each position. These values were approximately uniform along both chromosomes (Supplementary Figure S1), thus suggesting that the assembly was correct.

### 2.3. Mitogenome annotation

Initial annotation was performed using Mitofy (Alverson et al., 2010), with further manual correction. To verify exon boundaries, we mapped RNA-seq reads by CLC Genomics Workbench v. 7.5.1 to the sequences of the chromosomes, requiring at least 80% of a read’s length to map with at least 90% sequence similarity. The relaxed setting for a percent of read length to be mapped allows mapping reads spanning the splice junctions and the relaxed setting for sequence similarity allows mapping of reads to regions with a high density of RNA editing sites. The alignment was visualized in CLC Genomics Workbench 7.5.1.

To check if there are any unnoticed protein-coding genes, we visually inspected the alignment to search for unannotated regions in the chromosomes with high coverage by RNA-seq reads. This search did not add new genes.

The search for genes encoding selenocysteine tRNAs (tRNA-Sec) was done for *Hypopitys monotropa* and *Vaccinium macrocarpon* by an online-version of Secmarker 0.4 (Santesmasses et al., 2017) with the default parameters. The SECIS element prediction was performed by an online-version of SECISEARCH3 (Mariotti et al., 2013), current as of 5 May 2017, with the default parameters.

### 2.4. RNA editing analysis

To obtain information on RNA editing sites, we artificially spliced gene sequences and mapped RNA-seq reads to them with the parameters stated above for RNA-seq reads mapping. The variant calling was performed by CLC Genomics Workbench 7.5.1. We considered a site in a CDS as edited if it was covered by at least 20 reads and at least 10% of the reads differ in this position from the genomic sequence.

To estimate the fraction of RNA editing sites that change amino acid sequence of corresponding proteins, we use a measure that was already used in several previous studies (Chen, 2013; Xu and Zhang, 2014; Zhu et al., 2014), but hasn’t yet got its own name. Here we call it dEN/dES. It is calculated similarly to dN/dS, but while dN/dS counts the ratio of nonsynonymous and synonymous substitutions per nonsynonymous and synonymous sites, dEN/dES (“E” here stands for “editing”) counts the ratio of nonsynonymous and synonymous RNA editing events per nonsynonymous and synonymous sites. If editing sites are distributed in a CDS irrespectively to whether they do or do not change amino acids, dEN/dES will be close to one.

To compare the RNA-editing sites in the mitochondrial mRNAs of *H. monotropa* with those in the mitochondrial mRNAs of other species, we used data on the RNA-editing in *Arabidopsis thaliana* (Giege and Brennicke, 1999) and *Oryza sativa* (Notsu et al., 2002). These data are incorporated in the annotations of *A. thaliana* and *O. sativa* in GenBank, with accession numbers NC_037304.1 and BA000029.3 respectively.

### 2.5. Search for sequences transferred to the mitogenome

In order to find the sequences transferred from the plastome to the mitogenome (known as “MIPTs”, which stands for MItochondrial Plastid Transfers), we aligned plastid genes of *Camellia sinensis* to the sequences of the *H. monotropa* mitogenome. *Camellia sinensis* was chosen because its plastome contains a complete set of typical plastid genes and is phylogenetically closest to *H. monotropa* (as of May 2017) among all sequenced plants with such complete set. We searched for transferred genes and not intergenic regions, because we were interested in transfers that may be functional in the first place. Proteins of *Camellia* were aligned to the sequences of the mitochondrial chromosomes by TBLASTN with the maximum allowed *e*-value of 10^−3^. tRNA and rRNA coding genes were similarly aligned to the sequences of the chromosomes by BLASTN. The matching regions in the chromosomes were then aligned by BLASTX to NCBI NR (for regions that matched to *Camellia* proteins) and by BLASTN to NCBI NT (for regions that matched to *Camellia* RNA coding genes). If, for a region in the mitogenome, the best matches in the database were to sequences belonging to plastomes, that region was considered a MIPT. To calculate the number of frameshifting indels and nonsense mutations in the transferred regions they were taken together with their 200 bp-long flanking sequences on both ends and aligned to homologous genes from *C. sinensis*. The alignment was done by BLASTN with the default parameters. The resultant alignments were inspected by eye. To search for possible horizontal gene transfers from fungi, the mitogenome sequences were split into windows 500 bps each, with a step size of 50 bps. These windows were aligned by BLASTX to NCBI NR and also by BLASTN to NCBI NT. The maximum *e*-value allowed for the matches was 10^−5^.The regions that yielded significant matches to fungi were extracted and aligned back to NCBI NR, in order to find if they are uniquely shared by *H. monotropa* and fungi.

The analysis of the origin of the *cox1* intron in *H. monotropa*was done as follows. First, we aligned its sequence to the NCBI NT database by BLASTN online with the default parameters, taking the best 100 matching sequences. To these sequences we added the sequences of all *cox1* introns from Ericales that were not among the best 100 matches. Then, we aligned these sequences by the MAFFT server at https://mafft.cbrc.jp/alignment/server/(Katoh et al., 2019) with the default parameters. The unrooted phylogenetic tree of the *cox1* introns was built by RAxML 8.2.12 (Stamatakis, 2014) with 20 starting maximum parsimony trees and the GTR+Gamma model. The required number of bootstrap pseudoreplicates was automatically determined by the extended majority-rule consensus tree criterion (the “autoMRE” option).

### 2.6 Phylogenetic analysis

Genes common for mitochondrial genomes of 25 seed plants (*atp1, atp4, atp8, atp9, ccmC, cob, cox1, cox2, cox3, matR, nad1, nad2, nad3, nad4, nad4L, nad5, nad6, nad7, nad8, nad9*)were used for the phylogenetic analysis. Their sequences were concatenated and aligned by MAFFT(Katoh et al., 2017). The phylogenetic analysis was performed using RaXML (raxmlGUI v.1.3.1) with nucleotide sequences under GTR+gamma substitution model with 1000 bootstrap replicates. In order to infer possible horizontal transfers all protein-coding genes found in *H. monotropa* mitogenome were aligned and analyzed in the same way as the concatenated gene set.

## Results

### 3.1. Genome structure and gene content

Genome assembly resulted in two sequences. GC content of both sequences is ~ 45%. One of them is circular with a length of 106 Kb (Figure 1). It does not have any long repeats; the mapping of mate-pair reads shows the absence of pairs with abnormal insert sizes suggesting that this sequence is not a subject of recombination. The second fragment, the longer one (Figure 1), is ~704 Kbp long. It was assembled as linear and has a more complex structure. In particular, it contains several long repeats. As shown in Figure 2, three pairs of long direct repeats are observed: the beginning of the chromosome and the 80–90 Kb region, the end of the chromosome and the 455–470 Kb region, the 415-420 Kb region and the 495-500 Kb region. Long inverted repeats are also found: between the 260-262 Kb region and the 657-659 Kb region. A peculiar feature is a gradual decrease of coverage on both of its ends (Supplementary Figure S1). This suggests that copies of the mitogenome with different lengths of this repeat coexist in plant cells. Repeats at the ends of the genome are known in linear mitochondrial genomes of several fungi, animals and protists (see, e.g., the works of Janouškovec et al. (2013) and Kayal et al. (2012)) play an important role in their replication (Nosek et al., 1998). Similar mechanism could mediate the replication of the linear mitochondrial chromosome in *H. monotropa*.

**Figure 1.**
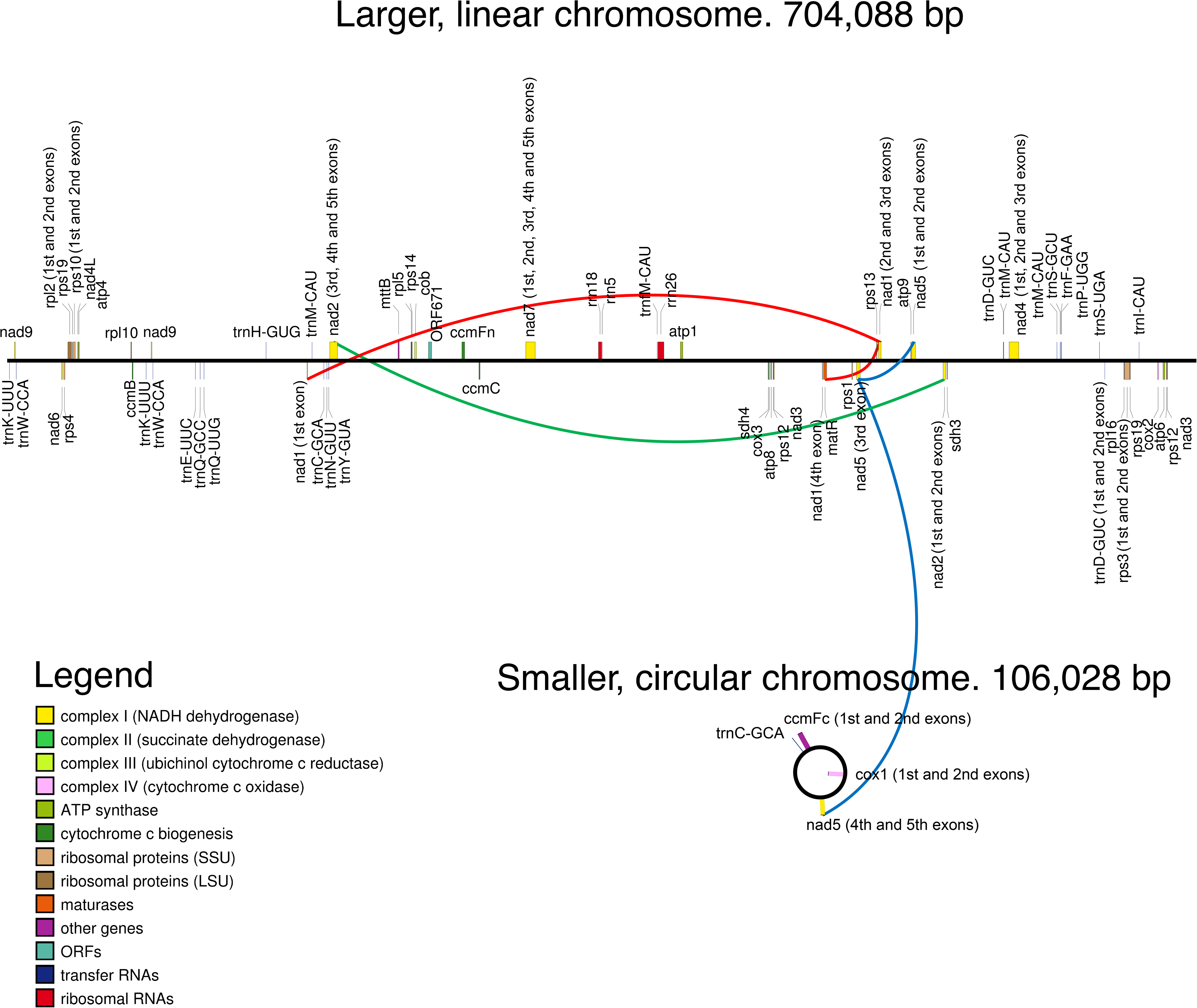
Maps of the mitochondrial chromosomes of *Hypopitys monotropa*. Trans-spliced introns are indicated by three colored lines: red in *nad1*, green in *nad2*, and blue in *nad5*.

**Figure 2.**
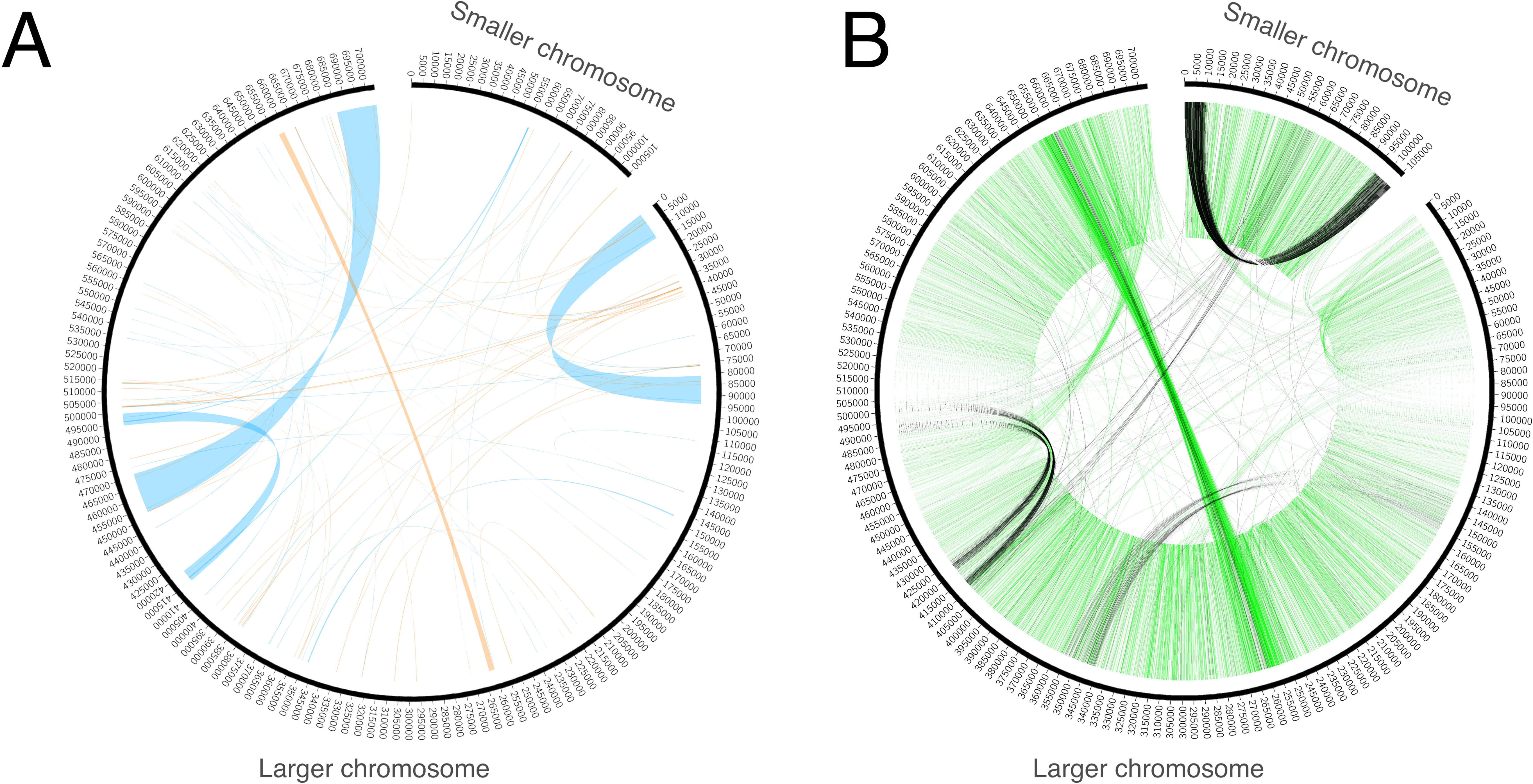
Repeats and mate-pair links in the mitochondrial chromosomes of *Hypopitys monotropa*. (A) Repeats within and between the chromosomes. Direct repeats are connected by blue lines, inverted repeats are connected by orange ones. (B) “Improper” mate-pair links indicate possible chromosome rearrangements. Read pairs with reads oriented in different directions (→ ← or ← →) are colored black and read pairs with reads oriented in the same direction (→ → or ← ←) are colored green. Only one of the two mate-pair libraries, that with the longer insert sizes (8279 bp on average, standard deviation 2583 bp), was used to build this diagram. A pair is considered improper if its reads are mapped not in the orientation→ ←, or are mapped on different chromosomes, or are mapped in the orientation → ← but are separated by more than 20000 bp. Closely situated reads in the orientations → →and ← ← that make a green “torus” across the genome (1.8% of all mapped mate-pair reads) likely represent the artifact of library preparation.

There are some mate-pair links between the two chromosomes, which suggests that they may recombine. However, the coverage of the smaller chromosome by mate-pair inserts (Supplementary Figure S1) has no sharp drops; therefore it is unlikely that these chromosomes join frequently.

In order to check for the presence of sequences transferred from fungi we performed BLAST search against all fungal sequences available in NCBI. Though BLASTX search identified several regions that had similarity to the hypothetical protein from *Rhizophagus irregularis* (Glomeromycota) genome, the same regions also have high similarity to other plants (Supplementary Table S1). This indicates that they are not the result of recent HGT mediated by mycoheterotrophy but rather either the result of ancient integration of fungal sequence into the genome of common ancestor of *Hypopitys* and other flowering plants or parallel integration of some mobile genetic elements (e.g. mitoviruses) in the fungal and plant genomes. At least one of these regions have high similarity to RNA-dependent RNA polymerase gene from a known plant mitovirus supporting the latter explanation.

Regarding gene content, the circular chromosome contains only two full-length protein coding genes—*ccmFc* and *cox1* and three exons of *nad5* gene, while two other exons are located on linear fragment. Summary data for the annotation is presented in Table 1. All genes are supported by RNA-seq reads (Supplementary Figure S2, Supplementary Table S2) and most regions with high coverage of RNA-seq reads correspond to annotated genes and their flanking regions (Supplementary Figure S3).

**Table 1.** Summary data on the structure and annotation of *H. monotropa* mitogenome.

| ***Chromosome*** | ***Accession number (NCBI)*** | ***Length*** | ***Protein-coding genes*** | ***rRNA-coding genes*** | ***tRNA-coding genes*** | ***Pseudo genes*** | ***Genes with introns*** | ***Genes with cis-splicing*** | ***Genes with trans-splicing*** |
| --- | --- | --- | --- | --- | --- | --- | --- | --- | --- |
| *Larger (linear)* | *MK990822* | *704088* | *35.5* | *3* | *17-18* | *2* | *8.5\** | *6* | *2.5* |
| *Smaller (circular)* | *MK990823* | *106028* | *2.5* | *0* | *1* | *0* | *2.5\** | *2* | *0.5* |
| *Total* |  |  | *38* | *3* | *18-19* | *1* | *11* | *8* | *3* |

*\* fractional number of genes indicates that one gene has exons in both circular and linear fragments.*

In general, in the gene content of the *H. monotropa* mitogenome is typical of flowering plants. Surprisingly it is even larger than in *V. macrocarpon*, a close photosynthetic relative: *atp6* and *rps14* are pseudogenized and *sdh3* and *rps3* are absent in *V. macrocarpon*. There is only one pseudogene in *H. monotropa,rps14*. Pseudogenization or loss of *rps14* is very common for flowering plants (e.g. (Figueroa et al., 1999)). In addition to the standard gene set we found a new ORF (ORF671) that hypothetically encodes a 671-aa protein. It has a weak similarity to a gene, which encodes a hypothetical protein in mitochondrial genomes of many flowering plants (closest match is the hypothetical mitochondrial protein Salmi_Mp020 from *Salvia miltiorrhiza*). There are 4 genes encoding ribosomal proteins of the large subunit and 8 genes of the small subunit. Regarding the RNA component of the ribosome, all three ribosomal RNAs (26S, 18S, 5S) typical of plant mitochondrial genomes are present in *H. monotropa*. The set of tRNAs consists of 17 tRNAs typical of mitogenomes of autotrophic plant species. trnD-GUC in *H. monotropa* is split into two exons. There are no tRNAs of plastome origin except for tRNA-Gly-GCC. A selenocysteine tRNA (tRNA-Sec) gene and a sequence required for selenocysteine insertion during translation (SECIS element) were reported in the mitogenome of *V. macrocarpon* (Fajardo et al., 2014). However, SecMarker, a novel specialized tool for the search of tRNA-Sec (Santesmasses et al., 2017) which has higher sensitivity and specificity than tools used by Fajardo and co-workers doesn’t confirm the presence of a tRNA-Sec gene in the *V. macrocarpon* mitogenome. Thus, we suppose that the predictions of a tRNA-Sec gene and a SECIS element in the *V. macrocarpon* mitogenome are false positives. In *H. monotropa* we also did not find tRNA-Sec genes or SECIS elements. There is a region with moderate (78.8%) similarity to the presumable tRNA-Sec of *V. macrocarpon*. It is located upstream of *ccmC*, as well as one of the two presumable tRNA-Sec of *V. macrocarpon*. tRNA genes are known to have highly conserved sequences; in particular, the similarity of other *V. macrocarpon* and *H. monotropa* tRNA genes is 98-100%. This also suggests that the presumable tRNA-Sec of *V. macrocarpon* is not a functional gene but a pseudogene of some tRNA-coding gene.

Though being highly conserved in coding sequence, mitochondrial genes sometimes differ in intron content. For example, *cox2* may consist of three exons (*D. carota*, *V. vinifera*), two exons (*C. paramensis*, *A. thaliana*), or a single exon (*V. macrocarpon*). In many angiosperm lineages, the *cox1* gene contains a group I intron, presumably acquired by multiple horizontal transfer events (Cho et al., 1998; Sanchez-Puerta et al., 2011, 2008), though there is an opposite hypothesis that postulates a single HGT and multiple losses (Cusimano et al., 2008). The *cox1* intron is highly overrepresented in the parasitic plants that have been examined to date (Barkman et al., 2007), though the hypothesis that parasitism may serve as a mediator of horizontal intron transfer is not supported by phylogenetic analysis (Fan et al., 2016). *V. macrocarpon* lacks an intron in *cox1*, while in *H. monotropa* it is present. Phylogenetic analysis indicates that the intron in *H. monotropa* has the same origin as in *Pyrola secunda* (Supplementary Figure S4) and is thus presumably vertically inherited from the common ancestor of *Pyrola* and *Hypopitys*. In other genes *H. monotropa* has the same intron content as *V. macrocarpon*. Three genes—*nad1*, *nad2*, *nad5* have trans-spliced transcripts. Notably, *nad5* exons are located in different chromosomes– exons 1, 2 and 3 in the linear one and exons 4 and 5 in the circular, suggesting the case of interchromosome trans-splicing. It was reported in other plants that have multichromosomal mitogenomes (Lloyd Evans et al., 2019; Sloan et al., 2012).

Detailed data about genes, their intron-exon structure and expressionis presented in Supplementary Table S2.

### 3.2. Sequences transferred to the mitogenome

In the mitochondrial genome we found fragments with high similarity to plastid genes, both of those that are present in the *H. monotropa* plastome (*matK*, *rps2*, *rps4*) and those that were lost (*rpoB*, *C1*, *C2*, *psbC*, *ndhJ*, *B*, *ycf2*) (Supplementary Table S3). These fragments are derived from horizontal transfers from plastids (MIPTs), presumably intracellular. Such events are frequent in plant mitochondria (for example, see the works of Alverson et al., (2011) and Grewe et al., (2014)). A rarer case of an interspecies horizontal transfer from mitochondrial genomes of other plants that already harbor inserts from plastid genome is also a source of MIPT, which are called in this case foreign MIPT (Gandini and Sanchez-Puerta, 2017). MIPTs in non-photosynthetic plants are of particular interest. If mitochondrial copies of plastid genes that were lost or pseudogenized in the plastome retain intact ORFs, potentially the reverse switch from heterotrophy to autotrophy is possible. Mitochondrial genome is in this case a “refugium” of plastid genes. The *Castilleja paramensis* mitogenome contains 55 full-length or nearly full-length plastid genes and only about half of them are obvious pseudogenes (Fan et al., 2016). By now the only example of functionality of mitochondrial genes transferred from plastids are tRNA genes of plastid origin recruited in the protein synthesis in mitochondria (Joyce and Gray, 1989). In *H. monotropa* all sequences that originated from plastid protein-coding genes do not represent intact ORFs, keeping in most cases less than 50% of the initial length and/or carrying multiple frameshift mutations (Supplementary Table S3).

### 3.3. Mitochondrial gene expression and RNA editing

In order to gain insights into mitochondrial gene expression and RNA editing and to refine the annotation we sequenced and assembled the transcriptome of *H. monotropa* (Logacheva et al., 2016; Schelkunov et al., 2018). We found expression of all annotated protein-coding genes (here and further expression level is defined as the number of mapped read pairs divided by gene length). Minimal expression level (0.1) is observed for the hypothetical protein *ORF671*. Cytochrome *c* maturation factors are also expressed weakly (3 of 4 genes have expression <1). Genes of Complex I have medium expression (1—6). The highest expression is observed for *sdh4*, *cox3*, *atp1* and *atp9* (Supplementary Table S2).

RNA editing is a phenomenon widely observed in mitochondrial transcripts. The number and position of editing sites differs a lot from species to species and from gene to gene. In *H. monotropa* we identified 545 RNA-editing site present in at least one RNA-seq sample (Supplementary Table S4) under the editing level threshold 10% and coverage threshold 20x. Most of these editing events are observed in all samples (Figure 3) Most of them are C-U. However we found at least one A-G editing which is present in all samples at the level ~ 20 % (see discussion below). Transcripts of 33 genes out or 38 are edited; maximal RNA-editing density (measured as the number of sites per 100 bp) is observed for *ccmB*. Genes of Complex I demonstrate different RNA-editing densities, from 0.2 of *nad5* to 5.6 of *nad4L*. In *nad4L* RNA editing restores start codon; presumably the same is true for *V. macrocarpon* where *nad4L* is (mis)annotated as pseudogene. RNA-editing density in genes of Complex II, III is weak, is not over 1.1. At last, in *atp1*, *rpl16*, *rps12*, *13*, *ORF671* we did not find any editing events. Several genes with high expression have low level of RNA-editing (*atp1*, *rps12*), and *vice versa* (cytochrome *c* maturation factors) but generally there is no such tendency. Median dEN/dES across all genes is 1,23. There are no edited stop codons (which is expected because most of the editing events are C-U and stop codons lack C; however the observation of non C-U editing potentially enables this), but there are several stop codons that are introduced by editing (Supplementary Table S5). To compare the RNA editing in *H. monotropa* with other species, we used data on *A. thaliana* and *O. sativa*, the model plants where RNA editing is thoroughly characterized (Giege and Brennicke, 1999), (Notsu et al., 2002). Overall, there are 842 positions edited in at least one species in the protein-coding genes common for these three species (Supplementary Table S6). Only 151 (17.9%) of these positions are edited in all the three species. Notably, 37.9% of the positions are that are edited in at least one species have T in the species that do not have editing in this position, while only 6.1% have A or G. This is congruent with the results of (Edera et al., 2018) who found that the main pattern of the loss of RNA editing is the replacement of editing sites with thymidines. 337 (38,1%) positions have unedited cytosines.

**Figure 3.**
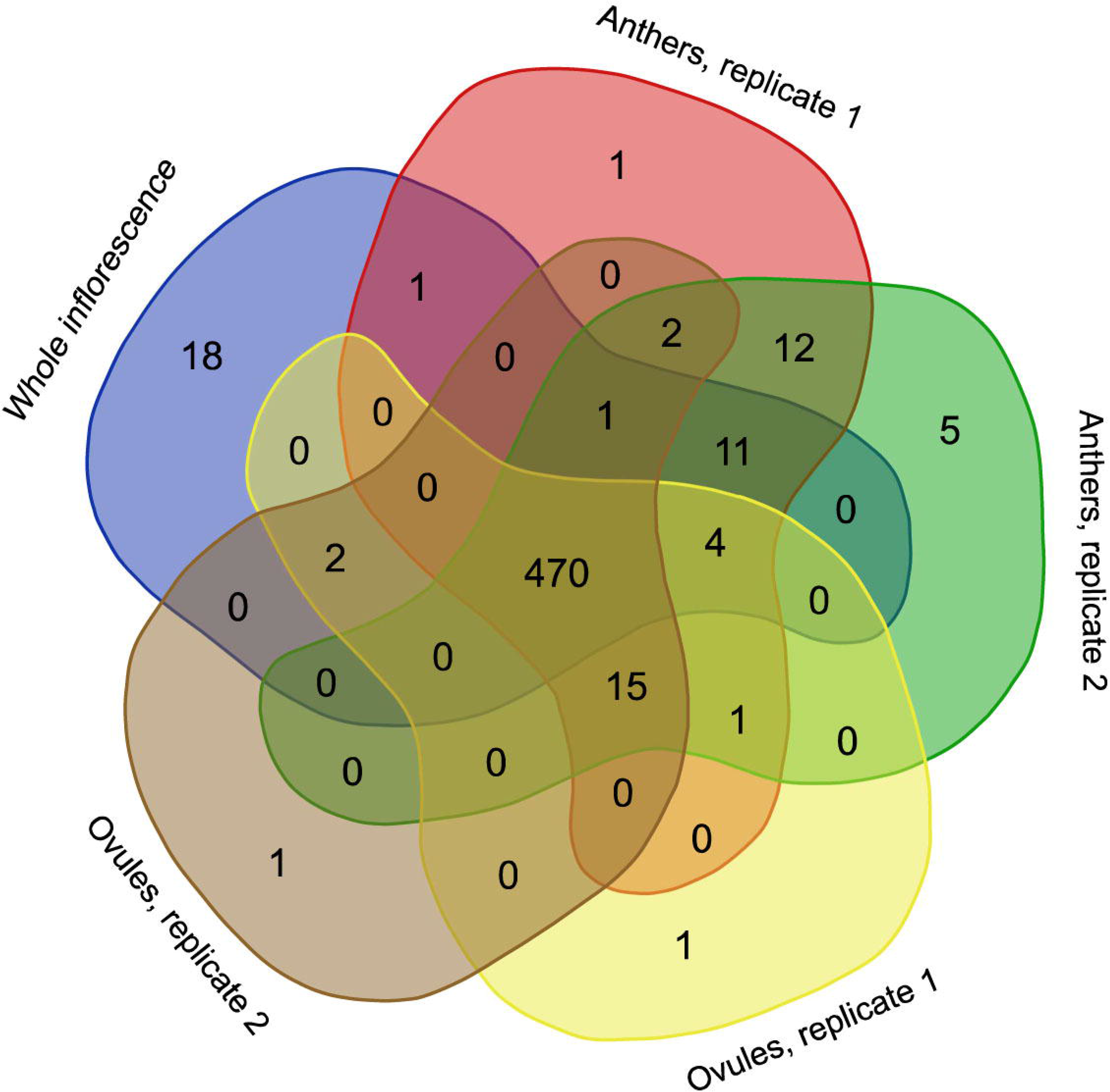
Venn diagram representing the occurrence of RNA editing events in different RNA-seq samples.

### 3.4. Phylogenetic analysis

As mentioned above, plant mitochondrial genomes are prone to HGT. It is often detected by incongruence of phylogenetic trees based on different genome regions (Bergthorsson et al., 2004; Cusimano and Renner, 2019). We performed phylogenetic analysis of the individual mitochondrial genes and found that the topologies are similar with regard to the placement of *H. monotropa* – it is always placed together with *V. macrocarpon* (excluding the cases of unresolved nodes) (see Supplementary figure S5). This evidences that no genes were acquired via HGT. The combined tree of all mitochondrial genes shared across 25 seed plant species shows topology similar to that of based on nuclear and plastid genes, with monocots representing monophyletic group and eudicots divided into two large groups – asterids and rosids. *H. monotropa* is sister to *V. macrocarpon*, and both are within the asterids, as expected (Figure 4). Notably, *H. monotropa* genes do not demonstrate any increase in the substitution rates.The same is true for the another MHT plant, *Petrosavia stellaris* (data from (Logacheva et al., 2014)). Earlier, parasitic plants were reported to have elevated substitution rates in all three genomes (Bromham et al., 2013); however, recent study encompassing broader sampling of parasitic plants shows that this is not a universal phenomenon (Zervas et al., 2019).

**Figure 4.**
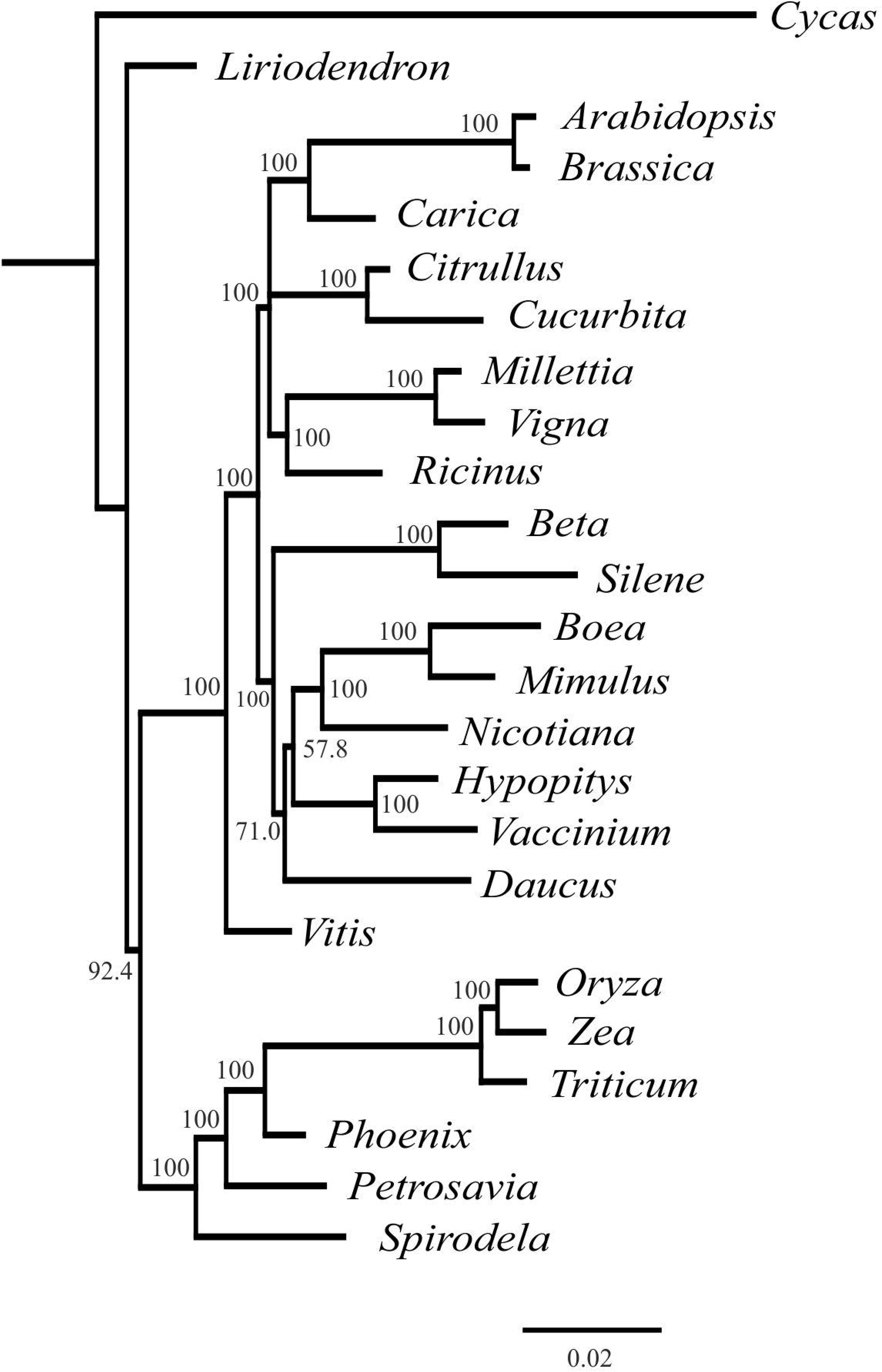
Phylogenetic tree based on the maximum likelihood analysis of nucleotide sequences of the 20 genes set. Values above nodes indicate bootstrap support. Branch lengths are proportional to the number of substitutions.

## Discussion

### 4.1 Mitogenome structure

All available evidence suggests that *H. monotropa* mitogenome has a linear fragment. While linear plasmids are found in mitochondrial genomes of several plants (e.g., in the work of Handa at el., (2002)), the linear fragment of the *H. monotropa* mitogenome lacks characteristic features of these plasmids: a terminal inverted repeat, the small size (not over 11 Kbp) and genes of RNA and DNA polymerases (reviewed in (Handa, 2008)). A linear structure is usual for mitochondrial genomes of fungi and protists (Janouškovec et al., 2013; Nosek et al., 1995). As well as other mycoheteroprophic plants, *H. monotropa* lives in intimate symbiosis with fungi (Min et al., 2012). One would hypothesize that linear fragment could be the result of either a contamination or HGT from fungi. However this is unlikely, for the following reasons: 1) DNA was isolated from inflorescences while mycorrhiza exists only in roots; 2) all potential fungal hosts of *H. monotropa* with a known mitogenome possess a circular chromosome and no linear plasmids; 3) there are no fragments with similarity to known fungal genomes; 4) all genes annotated on the linear fragments are typical plant mitochondrial genes. The fact that single circular molecule (“master circle”) is an oversimplified representation of the plant mitochondrial genomes and that they rather exist in vivo as a mix of circular, linear and branched forms is not novel (see, for example, the paper of Sloan en al. (2013)). However, a circular structure can usually be observed at the level of sequence assembly due to the presence of multiple repeats. This is not the case for *H. monotropa* where internal repeats are also present but their location, as well as the distribution of mate-pair links does not allow to reconstitute the master circle (see Figure 2).This suggests that the diversity of organization of plant mitochondrial genomes could be even greater than reported recently (Kozik et al., 2019).

A high level of convergence is observed in the gene set of plastid genomes of non-photosynthetic plants, whatever parasitic or mycoheterotrophic. They are characterized by a certain degree of reduction, which usually correlates with the time of transition to heterotrophy (for example, see the paper of Samigullin et al., (2016)) and follows the general gene loss model (Barrett et al., 2014). In contrast, mitogenomes of heterotrophic plants are very diverse in terms of structure, size and gene content. In *H. monotropa* the total size of the mitogenome is 810 Kbp, almost twice as large as that of *V. macrocarpon*. However this expansion is unlikely to be associated with the heterotrophy. Large mitogenomes are known for heterotrophic plants, in particular for the MHT orchid *Gastrodia elata* (~ 1.3 Mbp) (Yuan et al., 2018), and the parasitic *Cynomorium* (1 Mbp) (Bellot et al., 2016) and *Lophophytum mirabile* (Balanophoraceae) (~820 Kbp) (in the latter case the size is shaped by the fragments horizontally transferred from its host – see below). On the other extreme are highly miniaturized mitogenomes of *V. scurruloideum* (Skippington et al., 2015).

### 4.2. Horisontal gene transfer

HGT is a phenomenon very common in parasitic plants (Yang et al., 2016). HGT from host plants into the mitochondrial genome was shown for Rafflesiaceae (Xi et al., 2013), Orobanchaceae (Yang et al., 2016), *Cynomorium* (Cynomoriaceae) (Bellot et al., 2016), *Lophophytum mirabile* (Balanophoraceae). In the latter case horizontally transferred homologs replaced almost all native mitochondrial genes (Sanchez-Puerta et al., 2019, 2017). In contrast, there are no traces of HGT in *H. monotropa* mitogenome. Nuclear and mitochondrial genomes of a MHT orchid *Gastrodia elata* were recently characterized (Yuan et al., 2018), as well as the mitochondrial genome of an MHT dicot *Epirixanthes elongata* (Petersen et al., 2019) and there were also no observations of HGT. Despite that MHT plants are usually regarded alongside the parasitic plants that fed on other plants, the interactions between plant and its host are very different in these two cases. Parasitic plants develop specialized structures that integrate into the vascular system of a host plant and channel the flow of nutrients from the host to itself. Such connections are similar in many aspects to graft junctions and can be the route not only for nutrients but also for high molecular weight compounds including proteins and nucleic acids. The bidirectional transfer of nucleic acids through haustoria in the parasitic plant *Cuscuta pentagona* was shown (Kim et al., 2014). The transfer of RNA from host is hypothesized to mediate HGT into parasite genome. In contrast, such transfer is not known for mycorrhizal symbiosis. This emphasizes that despite similar–heterotrophic–strategy тут нужны дефисы?, plant parasites and MHT plants are fundamentally different in terms of the interaction with their hosts and, potentially, in many other features that stem from this. Thus the knowledge on MHT plant biology should be obtained using MHT models, not by the simple transfer of knowledge from the plant parasites. Many MHT plants are rare endangered plants with very small distribution ranges; in contrast, *H. monotropa* is widespread and is thus prospective as a model MHT plant.

### 4.3. RNA editing

RNA editing is the important characteristic of mitochondrial gene expression. It varies in plants in very broad ranges – from complete absence in a bryophyte *Marchantia polymorpha* to several hundreds and even thousands (for review see Ichinose and Sugita, (2016)). RNA editing in non-photosynthetic plants is of especial interest because many proteins are involved in RNA editing in both plastids and mitochondria (for example, see Bentolila et al., (2012)). Thus, the reduction of plastid genome and coordinated loss of nuclear genes involved in editing processes in plastids can influence mitochondrial RNA editing too. By now the data on RNA-editing in mitochondria for non-photosynthetic plants are scarce. C-to-U RNA editing in seven genes (*atp1*, *atp4*, *atp6*, *cox2*, *nad1*, *rps4*, and *rps12*) was found in *R. cantleyi* (Xi et al., 2013). In *V. scurruloideum* C-to-U editing was predicted computationally for the nine protein-coding genes (Skippington et al., 2015). In *H. monotropa* we observed editing in the majority of genes; almost all editing events are of C-to-U type, usual for plant mitochondria. Vast majority of the editing events are found in all samples (Figure 3, Supplementary Table S4). We observe a single non C-U editing site, the A-G editing in the *rps19* gene transcript. This editing is represented in all samples (inflorescence, ovules and anthers) at the level of 17-23% of reads (Supplementary Table S4). Considering that there is no coverage peak at this position, it is unlikely that this is a result of read mismapping. We hypothesize that the A-G conversion that we observe represents indeed not A-to-G editing, but adenosine-to-inosine (inosine reads as guanosine in sequencing data). This type of editing is usual for animals and fungi (Cattenoz et al., 2013; Liu et al., 2016), but was not found in plants. In many cases RNA editing has clear functional role (e.g. restoration of typical start codon in plastid genes *rpl2* and *psbL* (Kudla et al., 1992). We found one such case in *H. monotropa* (*nad4L*). However, the dEN/dES value in mitochondrial transcripts is 1.23 (does not significantly differ from 1, p-value of 0.060) which indicates that in general edited sites are unlikely to have functional significance. More detailed examination of RNA editing in MHT plants, including its dynamics in different organs and developmental stages is required to highlight potentially functional events.

## Conclusions

Non-photosynthetic plants represent ~1% of plant diversity and are an excellent model for the study of convergent evolution. Until recently the genomics of non-photosynthetic plants was focused on plant parasites; a usual assumption is that mycoheterotrophs - the plants that parasitize on fungi have basically the same patterns of genome evolution as plant parasites. In order to test this hypothesis and to expand our knowledge on MHT plants we characterized mitochondrial genome of *H. monotropa*. Also, using RNA-seq we performed a genome-wide analysis of gene expression and RNA editing. We showed that the mitogenome structure in *H. monotropa* is highly unusual: it includes small circular fragment and a large linear fragment that has on its ends multiple repeats that, presumably, function as telomeres. Further studies that include characterization of mitogenomes of other Ericaceae and *in vivo* analysis of *H. monotropa* mitochondria are required to investigate the details of the evolution, replication and functioning of such unusual mitogenome. The gene set is similar to that of autotrophic plants. All protein-coding genes are expressed and for the most of them (33 out of 38) we found editing of the transcripts. The intergenic regions of the mitogenome carry multiple sequences of plastid origin, including those of photosynthesis-related genes that are absent in *H. monotropa* plastome. We did not found either any traces of HGT from fungal hosts in *H. monotropa* mitogenome or the increase of nucleotide substitution rates. These new data highlight the contrast between mycoheterotrophic and parasitic plants and emphasize the need of the new model objects representing mycoheterotrophic plants.

## Acknowledgements

The authors thank Alexey Kondrashov (University of Michigan, Ann Arbor) for providing plant material and Artem Kasianov for assistance with data analysis..

---

**Supplementary Table S1.** Results of the BLAST search of regions that have significant similarity to fungal genes.

**Supplementary Table S2.** Characteristics of mitochondrial genes of *H. monotropa*

**Supplementary Table S3.** Plastid genes in mitogenome of *H. monotropa*

**Supplementary Table S4.** RNA editing of *H. monotropa* mitochondrial genes.

**Supplementary Table S5.**

Genes with stop codons that are introduced by editing.

**Supplementary Table S6.**

RNA editing of *H. monotropa* mitochondrial genes compared with *Arabidopsis thaliana* and *Oryza sativa*.

**Supplementary Figure S1.** Read mapping characteristics along the mitochondrial chromosomes of *Hypopitys monotropa*.

(A) Average insert size between mate-pair reads spanning over different genomic positions. The nearly uniform insert size distribution suggests that there are no misassemblies involving large deletions or insertions. The fluctuations at the ends of the larger chromosome result from small numbers of reads mapping to the very ends of the chromosome, which is linear.

(B) Number of mate-pair fragments covering each of the chromosomes’ positions. The absence of positions with zero coverage suggests that there are no misassemblies involving genome fragments’ rearrangements. The drops on the ends of the larger chromosome follow from its linearity.

(C) Coverage of the chromosomes by reads of all three sequencing libraries: the paired-end library and both mate-pair libraries. The coverage, though fluctuating, never reaches zero, thus suggesting the absence of misassemblies. Coverage at the ends of the smaller (circular) chromosome abruptly drops approximately sixfold, due to difficulty of mapping reads part of which map to the end and part to the beginning of the contig. In the larger (linear) chromosome, the gradual drops of coverage near the edges and near the positions 90000 bp and 450000 bp are due to the varying repeat copy number (see discussion in the main text).

**Supplementary Figure S2**

Coverage of *H. monotropa* mitochondrial CDS by RNA-seq reads

**Supplementary Figure S3**

Coverage of *H. monotropa* mitochondrial chromosomes by RNA-seq reads

**Supplementary Figure S4**

Phylogenetic tree of cox1 intron. Branches with bootstrap support below 70 are collapsed.

**Supplementary Figure S5**

Phylogenetic trees inferred from ML analysis of single mitochondrial genes. Branches with bootstrap support below 50 are collapsed.

**References**

