## Supplementary figures and images for "Mitochondrial genome of non-photosynthetic mycoheterotrophic plant *Hypopitys monotropa*,its structure, gene expression and RNA editing"

### Supplementary Figure S1

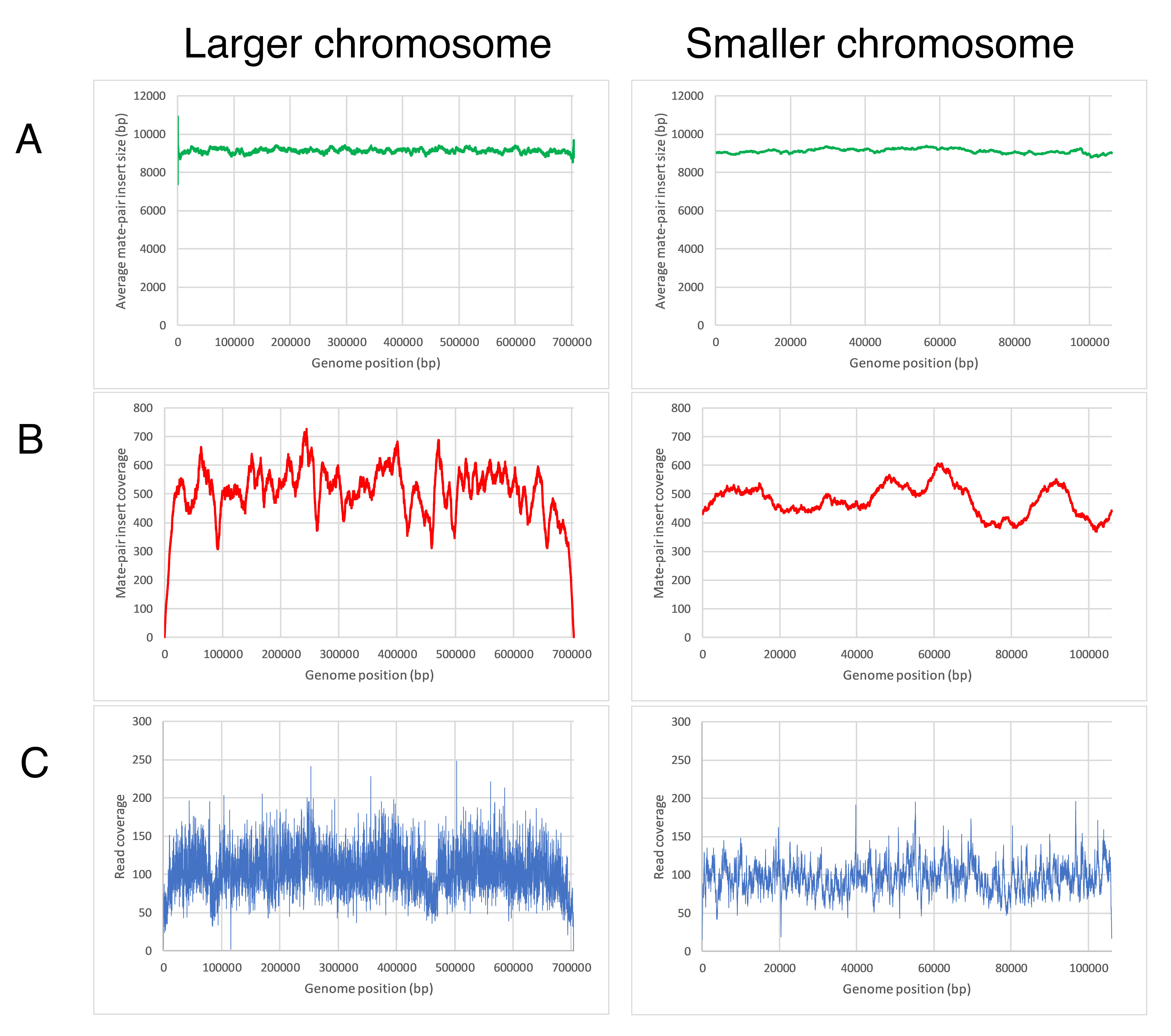

### Supplementary Figure S3

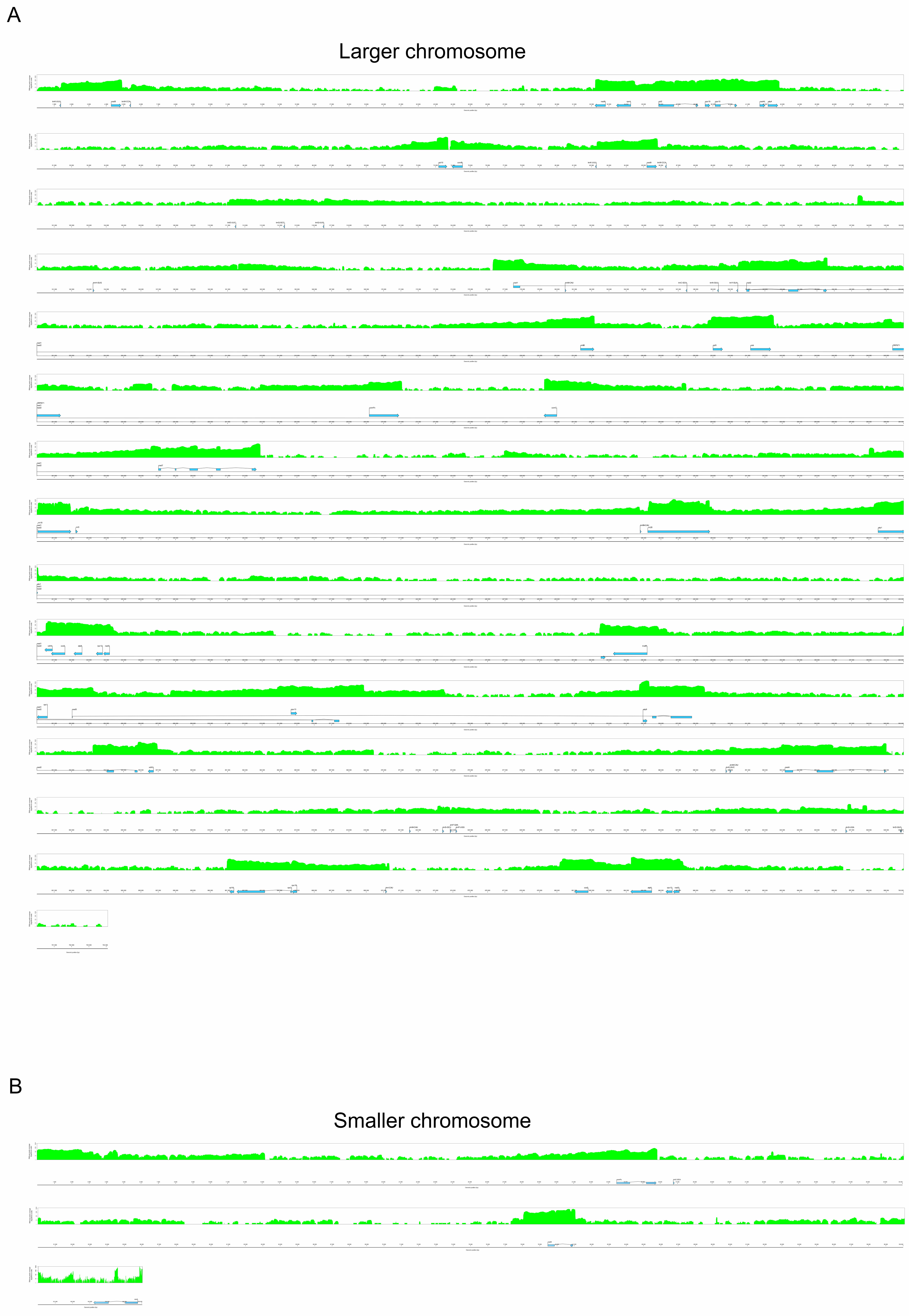

### Supplementary Figure S4

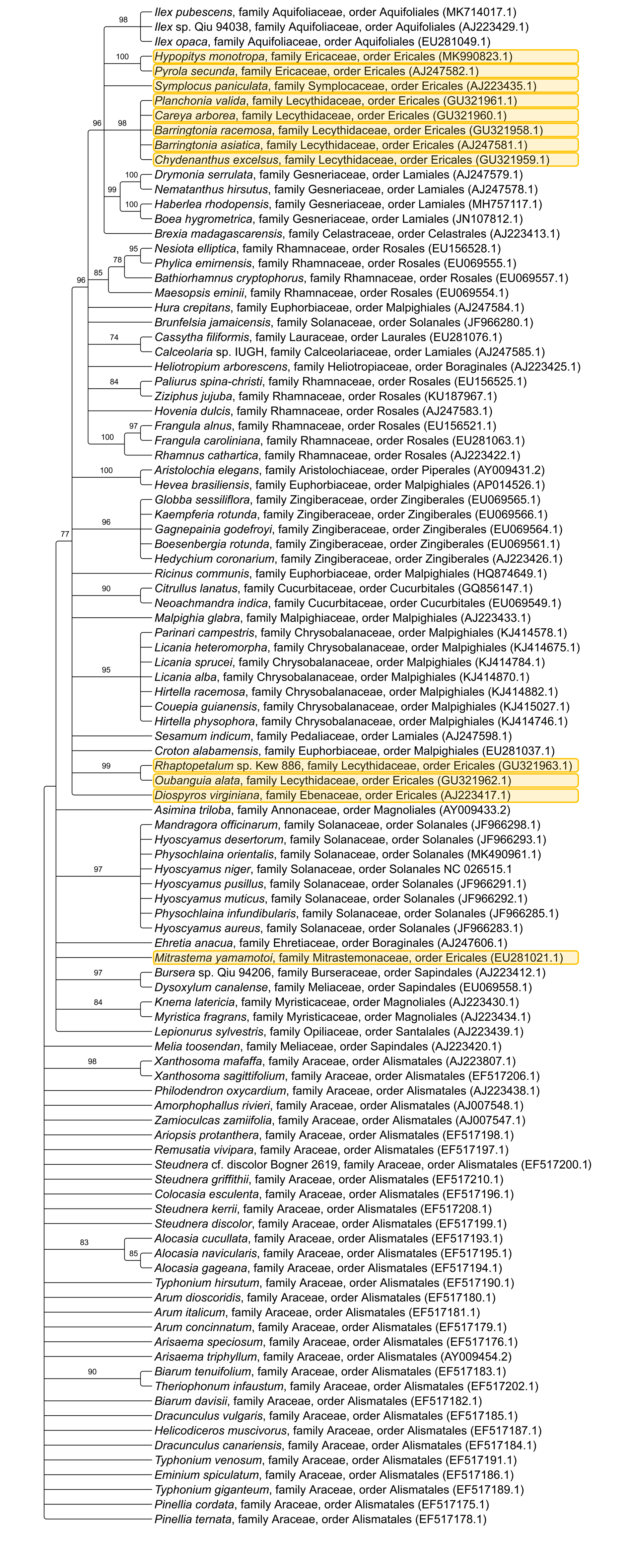
