## Supplementary Figure S2 for "Mitochondrial genome of non-photosynthetic mycoheterotrophic plant *Hypopitys monotropa*,its structure, gene expression and RNA editing"

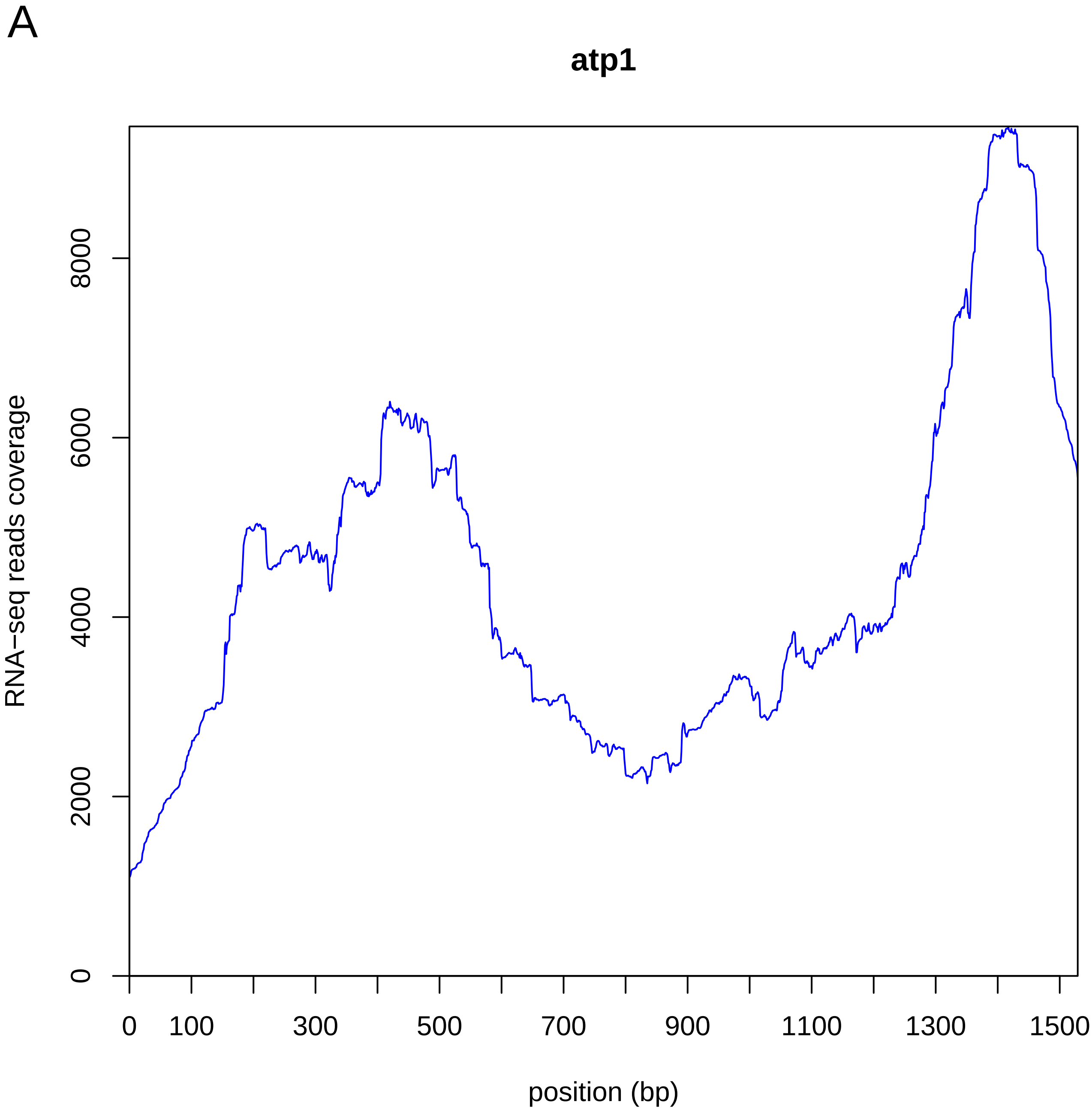
A
atp1
RNA−seq reads coverage
0
2000
4000
6000
8000
0
100
300
500
700
900
1100
1300
1500
position (bp)

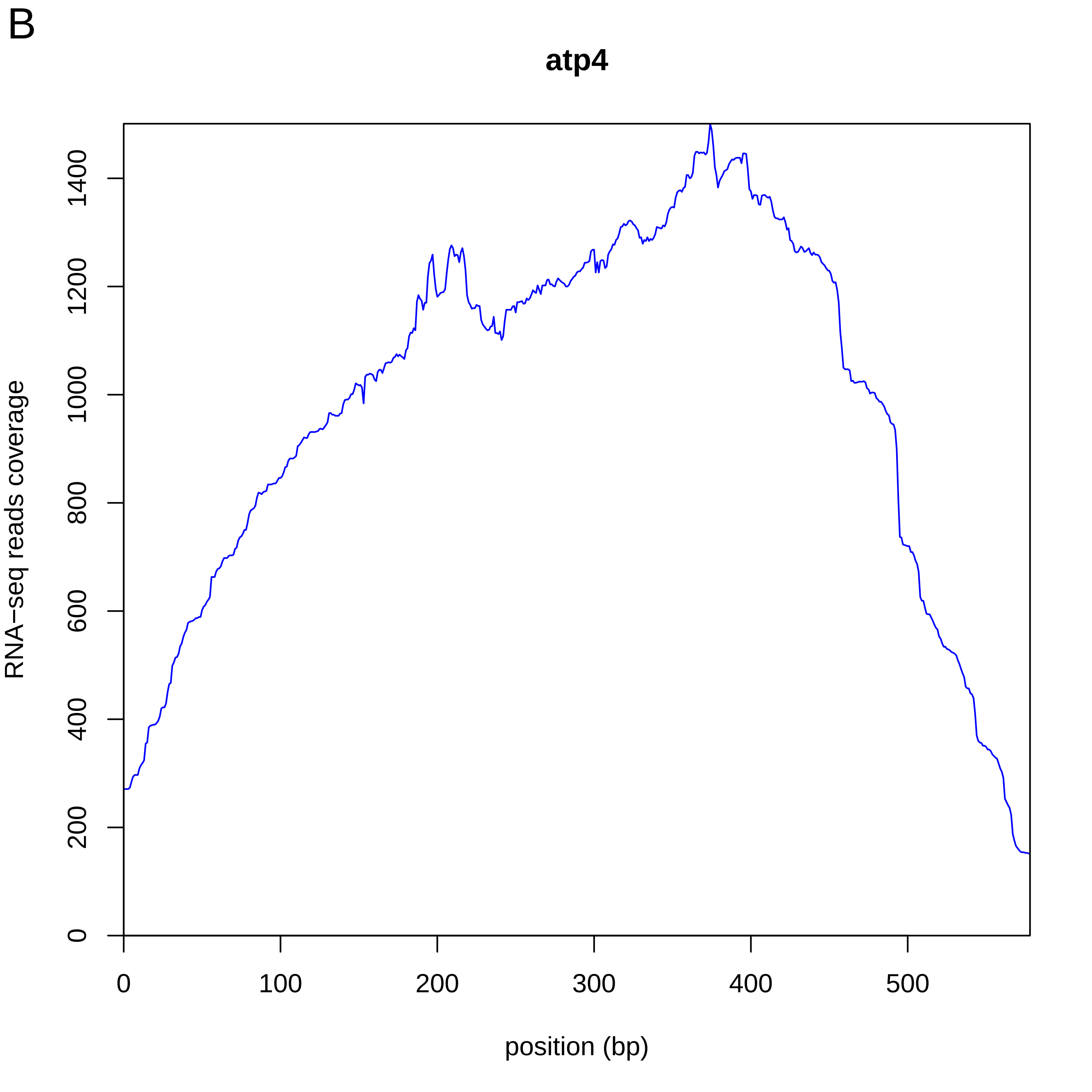
B
atp4
RNA−seq reads coverage
position (bp)
0
200
400
600
800
1000
1200
1400
0
100
200
300
400
500

C

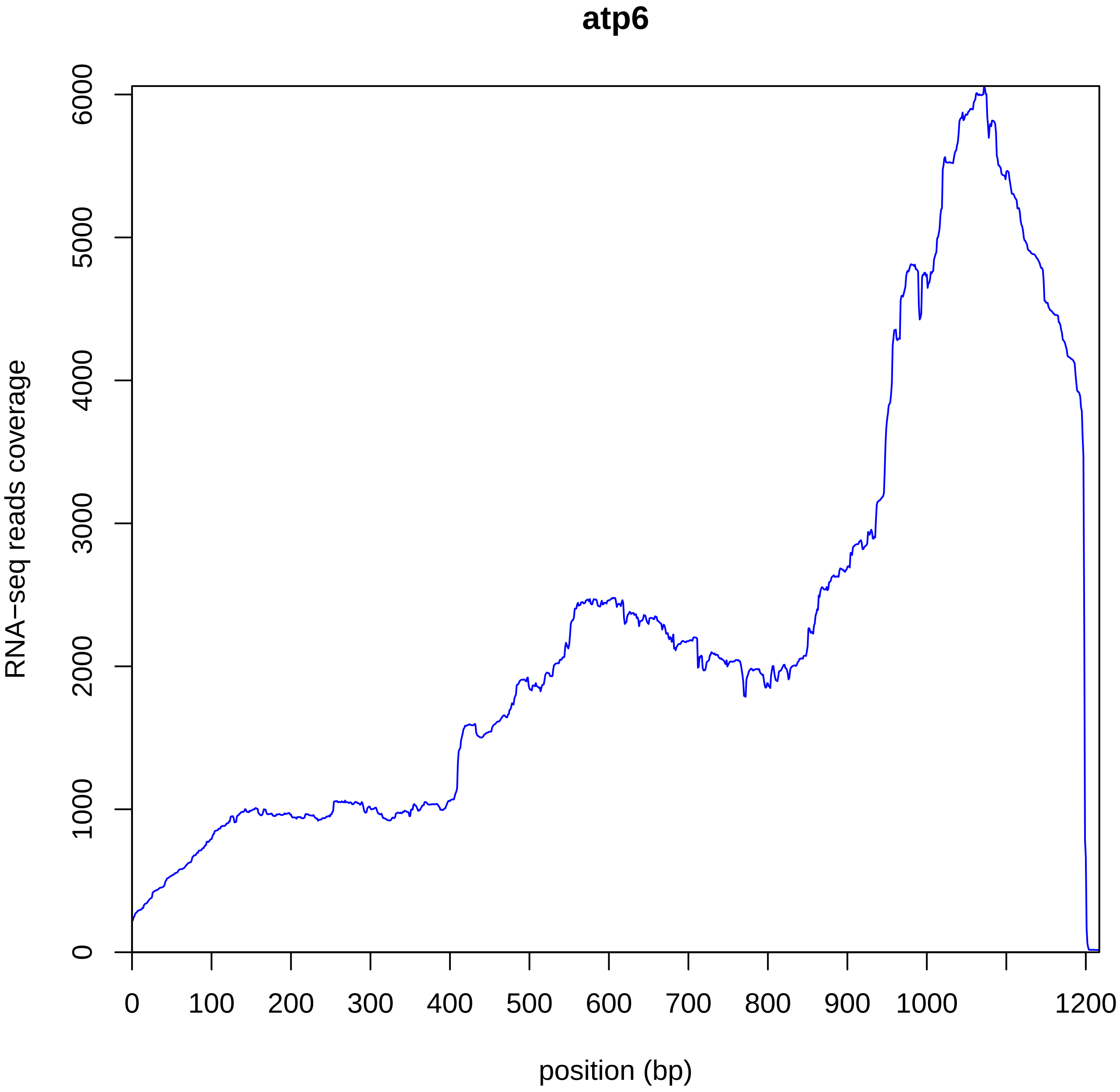
atp6
RNA−seq reads coverage
0
1000
2000
3000
4000
5000
6000
0
100
200
300
400
500
600
700
800
900
1000
1200
position (bp)

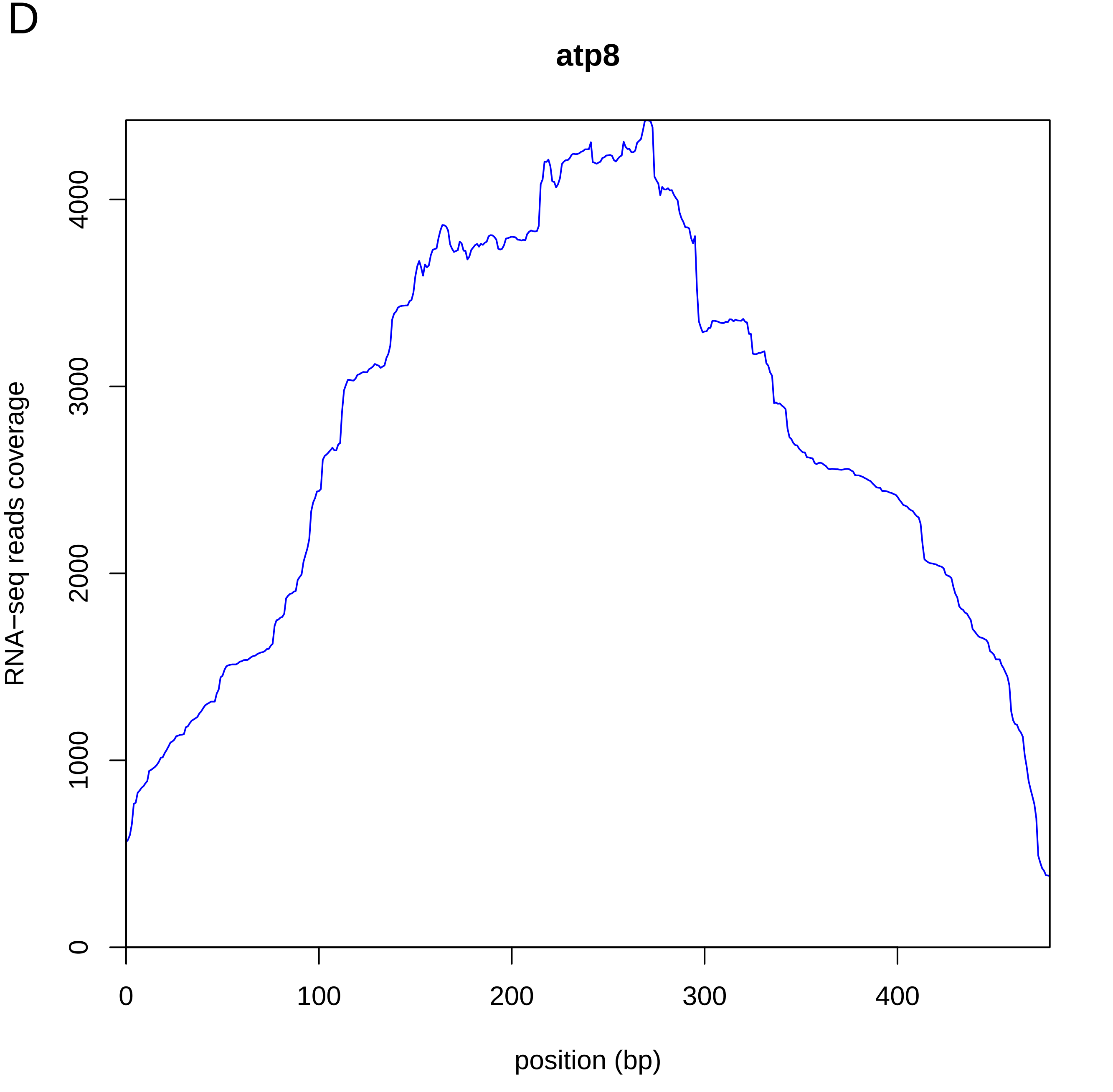

D
atp8
RNA−seq reads coverage
0
1000
2000
3000
4000
0
100
200
300
400
position (bp)

E

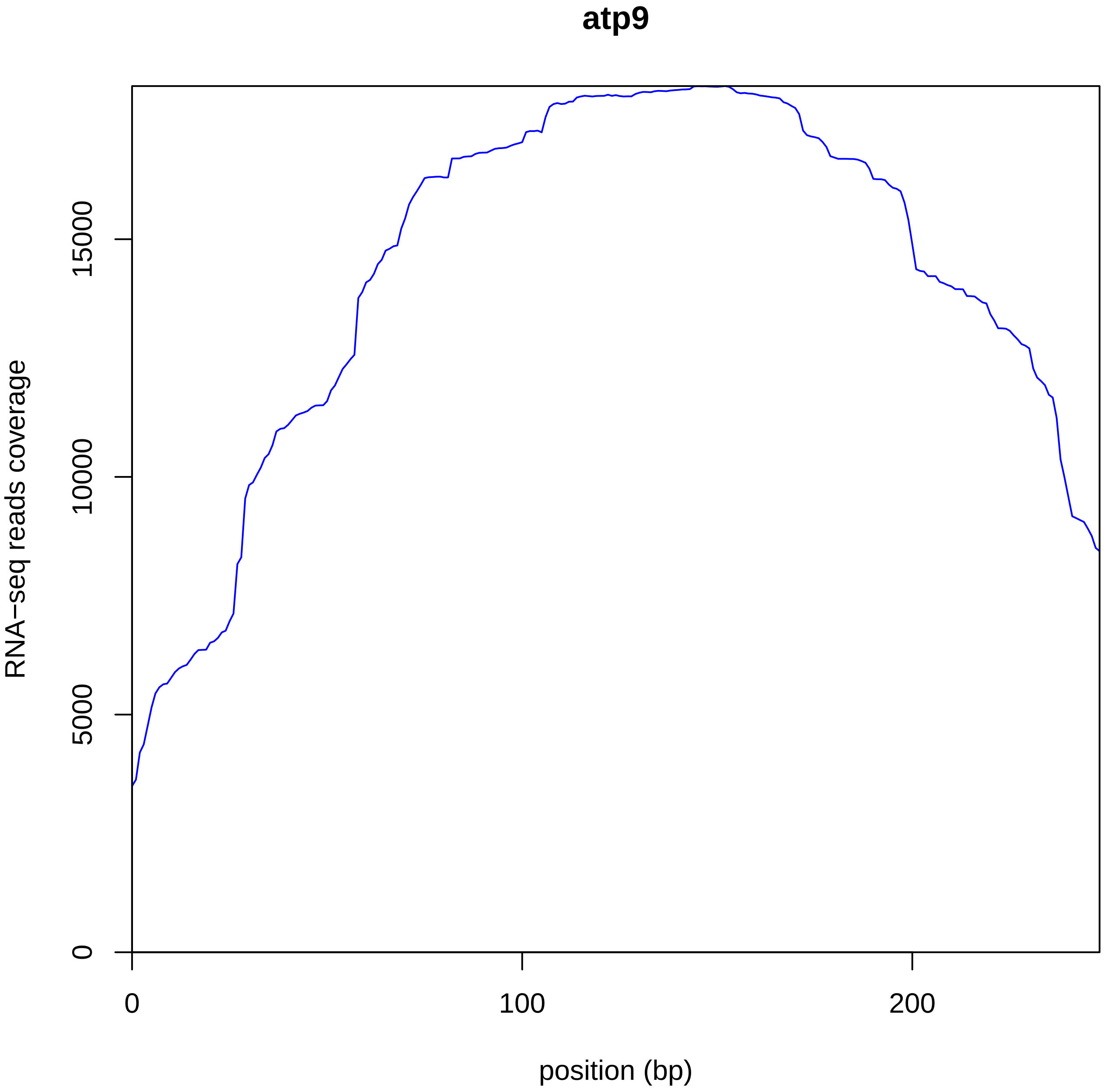

atp9
RNA−seq reads coverage
0
5000
10000
15000
0
100
200
position (bp)

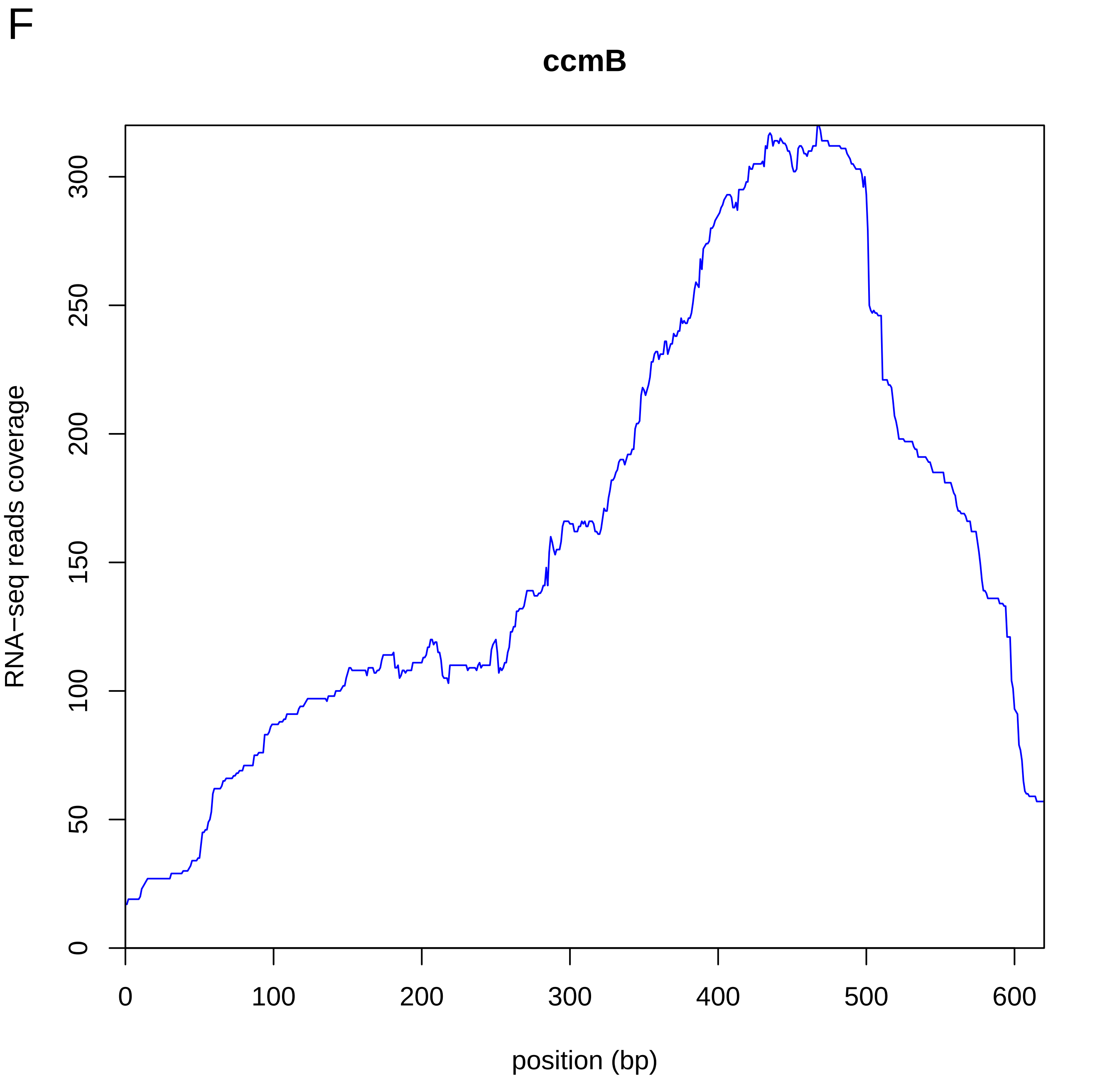
F
ccmB
RNA−seq reads coverage
0
50
100
150
200
250
300
0
100
200
300
400
500
600
position (bp)

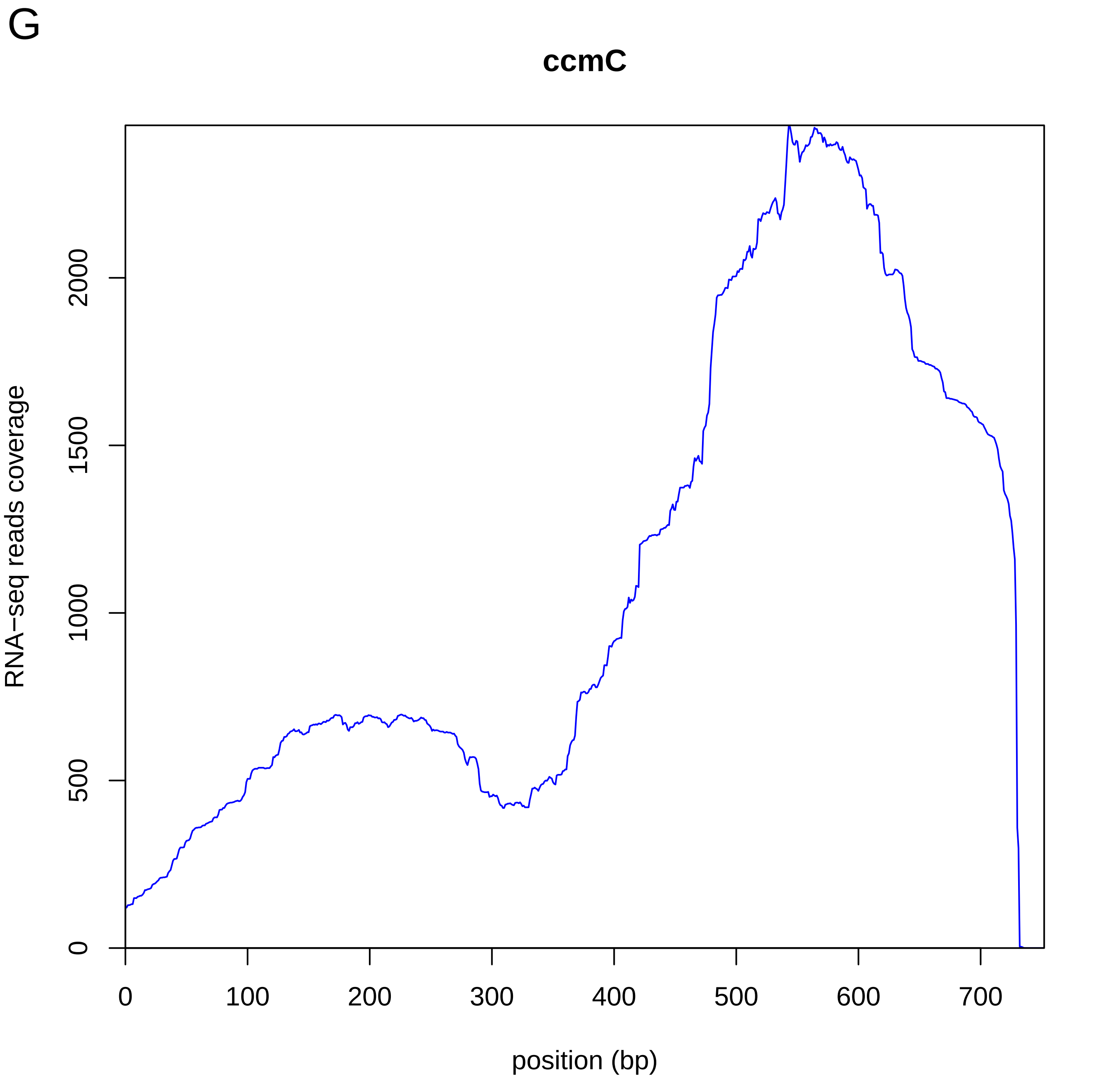
G
ccmC
RNA−seq reads coverage
0
500
1000
1500
2000
0
100
200
300
400
500
600
700
position (bp)

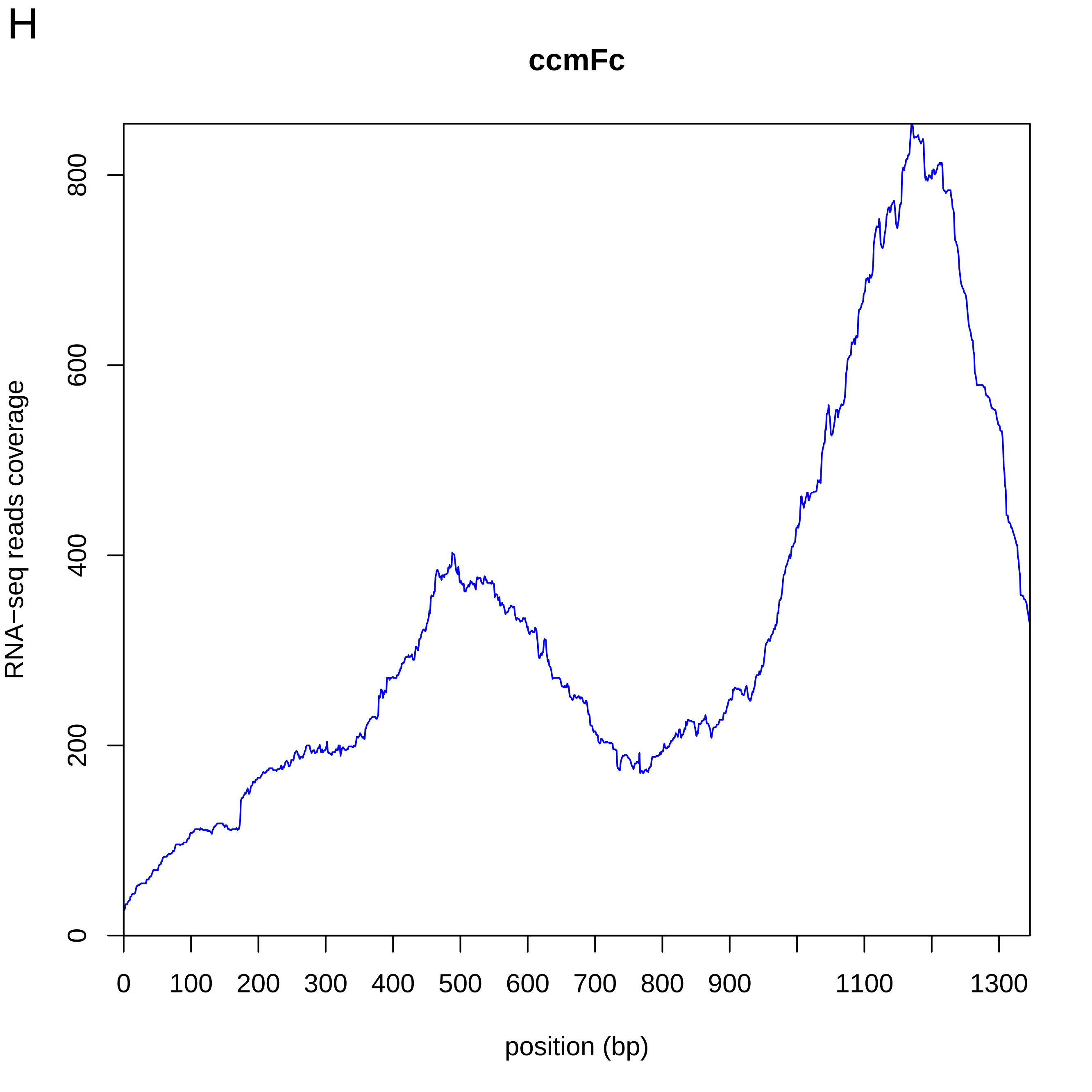
H
ccmFc
RNA−seq reads coverage
0
200
400
600
800
0
100
200
300
400
500
600
700
800
900
1100
1300
position (bp)

I

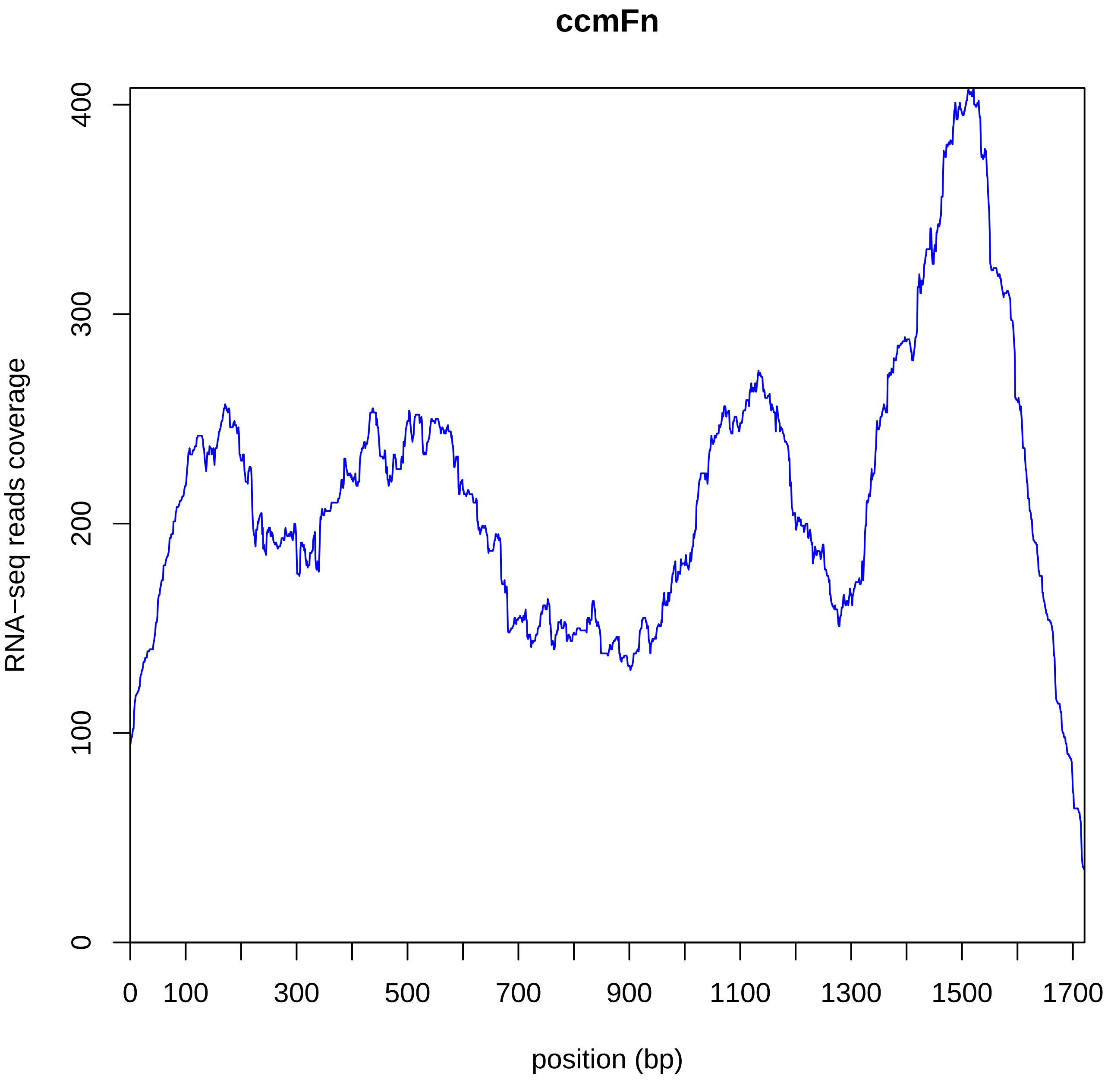
ccmFn
RNA−seq reads coverage
0
100
200
300
400
0
100
300
500
700
900
1100
1300
1500
1700
position (bp)

J

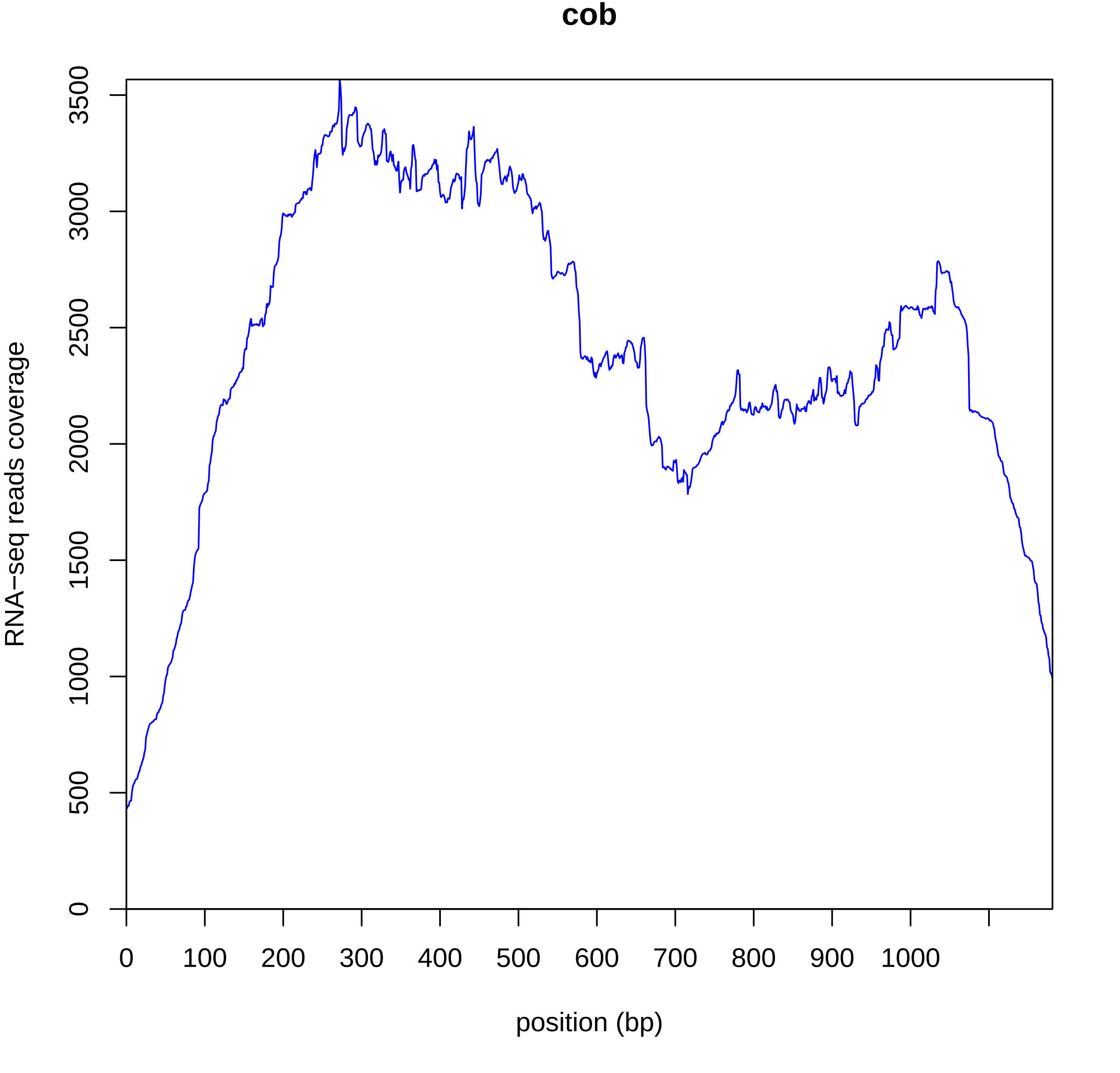
cob
RNA−seq reads coverage
0
500
1000
1500
2000
2500
3000
3500
0
100
200
300
400
500
600
700
800
900
1000
position (bp)

K

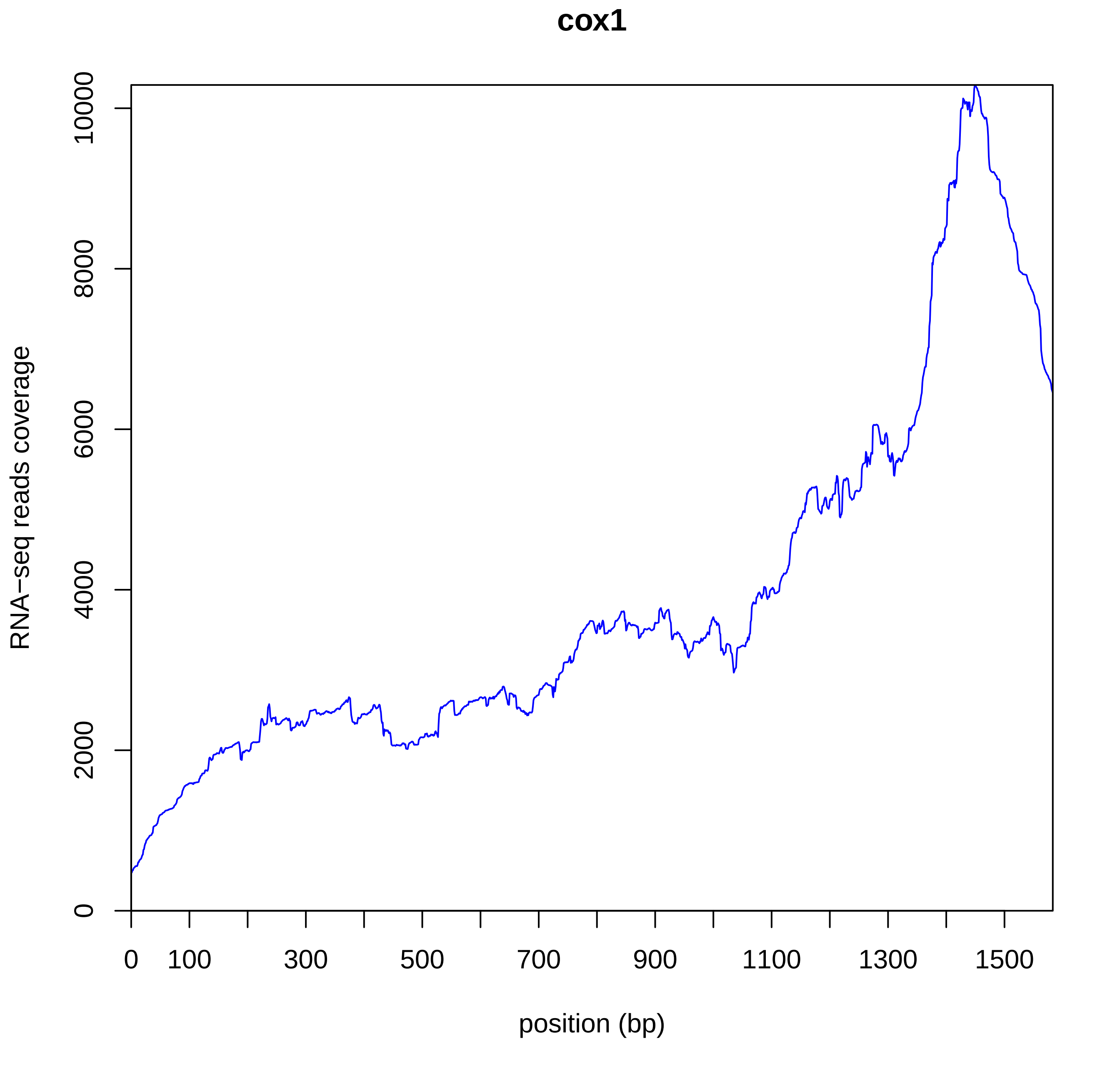
cox1
RNA−seq reads coverage
0
2000
4000
6000
8000
10000
0
100
300
500
700
900
1100
1300
1500
position (bp)

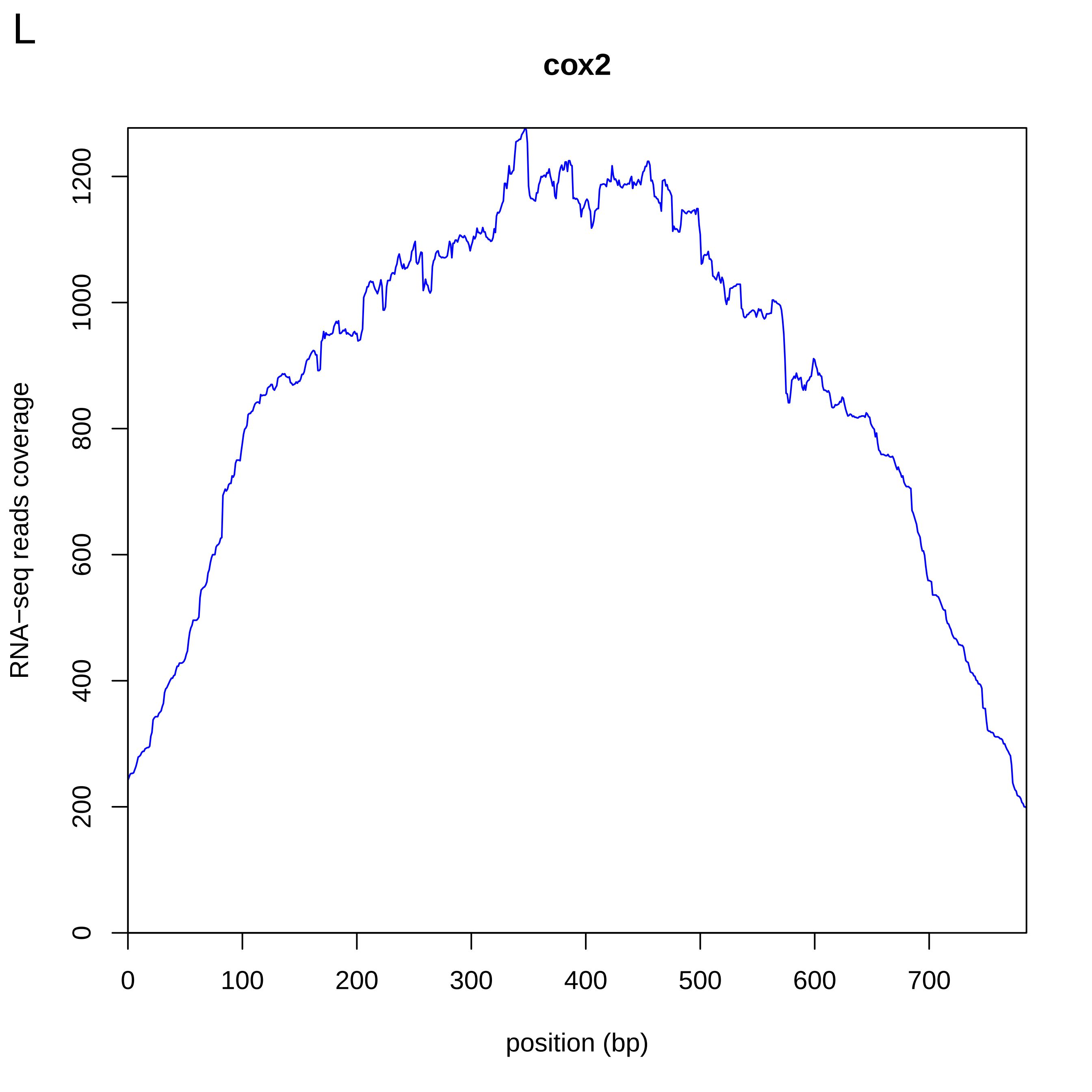

L
cox2
RNA−seq reads coverage
position (bp)
0
200
400
600
800
1000
1200
0
100
200
300
400
500
600
700

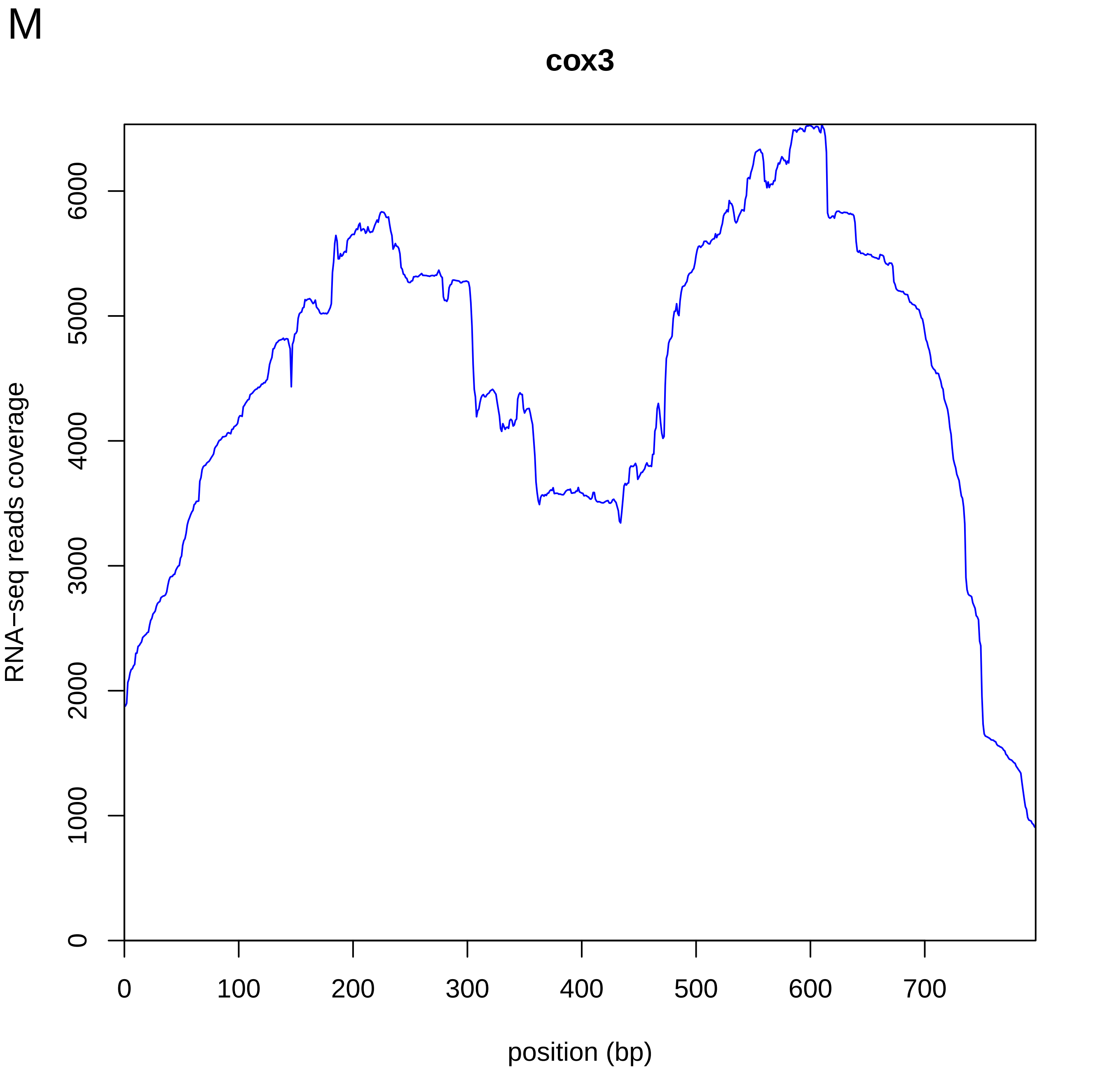
M
cox3
RNA−seq reads coverage
0
1000
2000
3000
4000
5000
6000
0
100
200
300
400
500
600
700
position (bp)

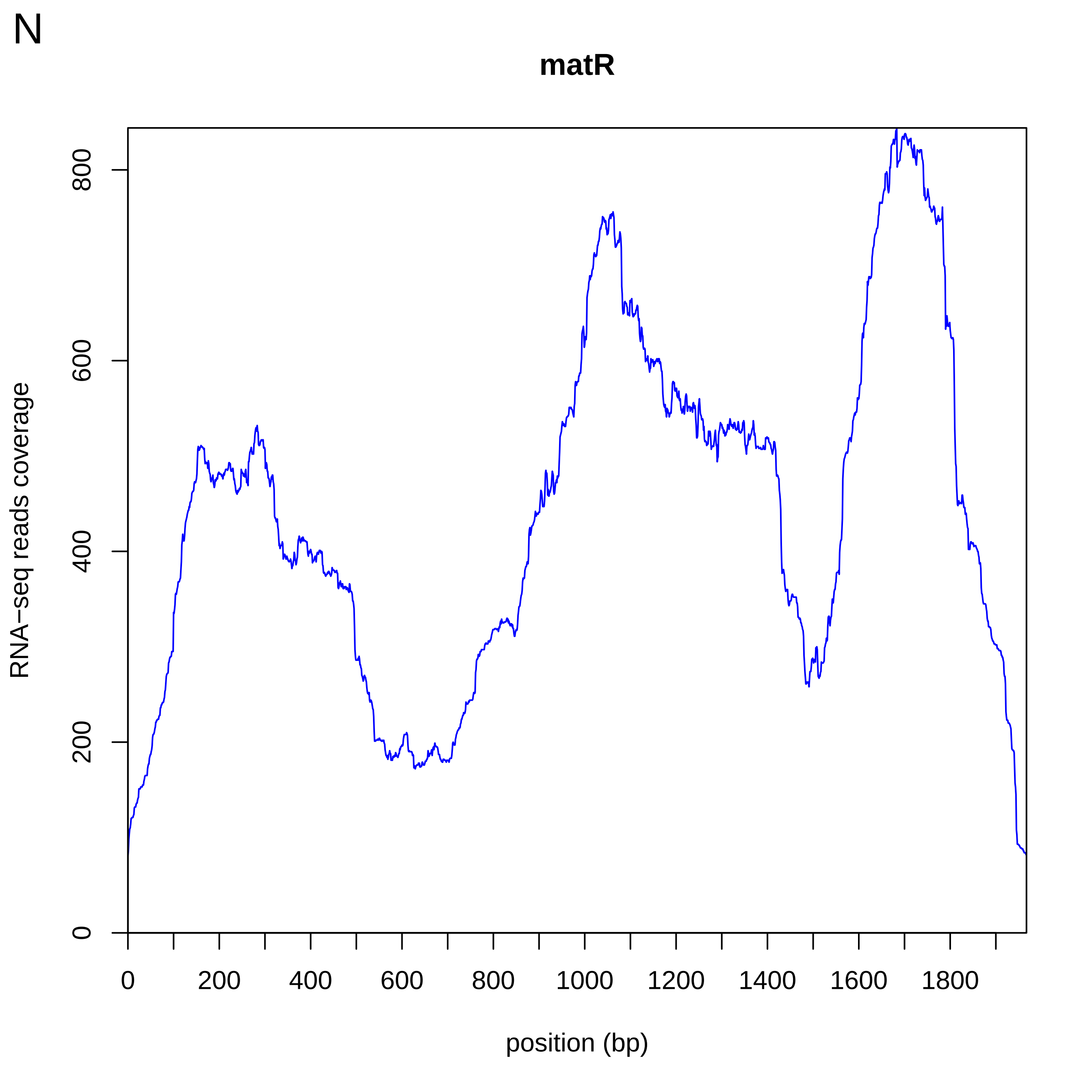
N
matR
RNA−seq reads coverage
0
200
400
600
800
0
200
400
600
800
1000
1200
1400
1600
1800
position (bp)

O

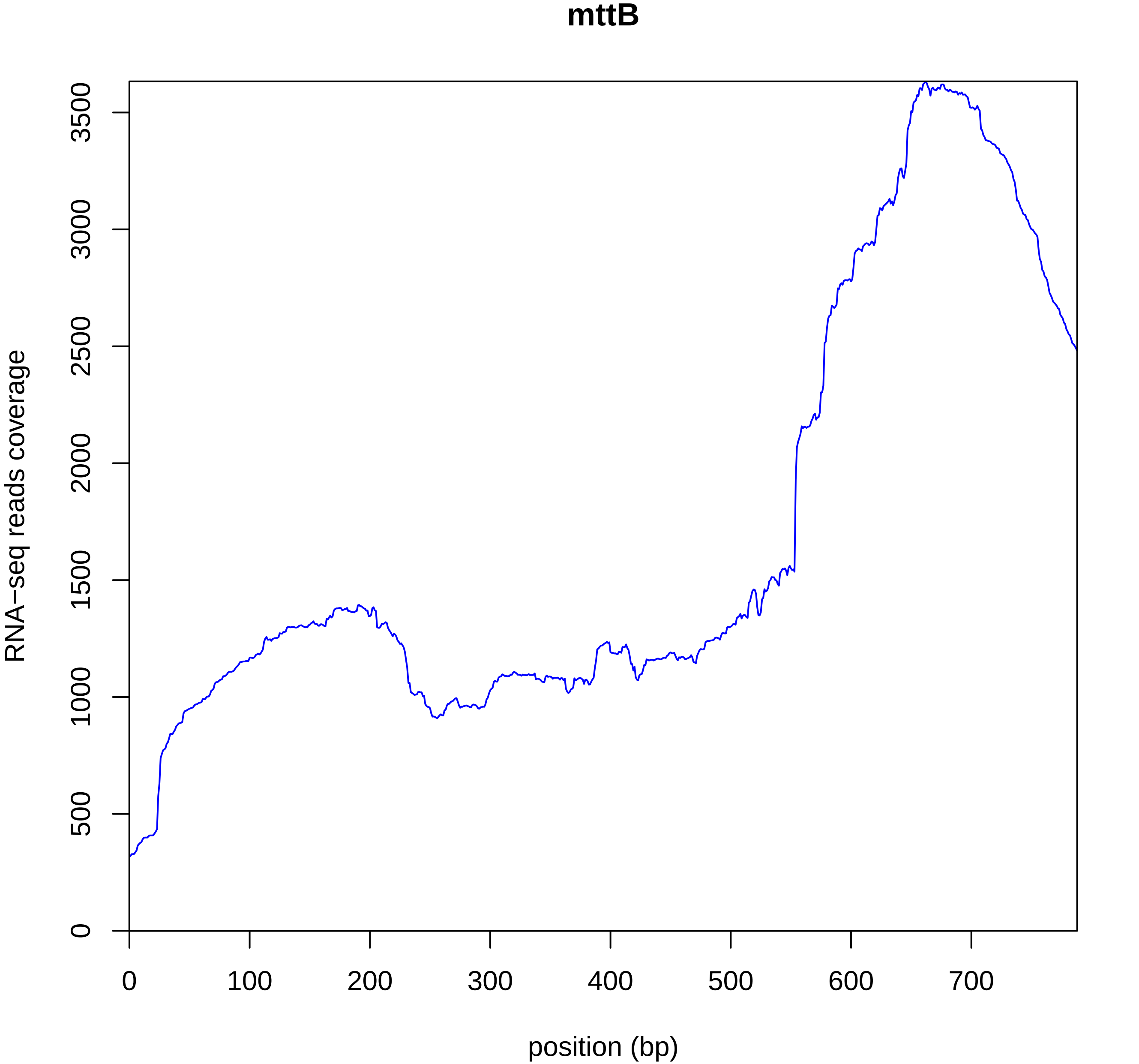

mttB
RNA−seq reads coverage
0
500
1000
1500
2000
2500
3000
3500
0
100
200
300
400
500
600
700
position (bp)

P

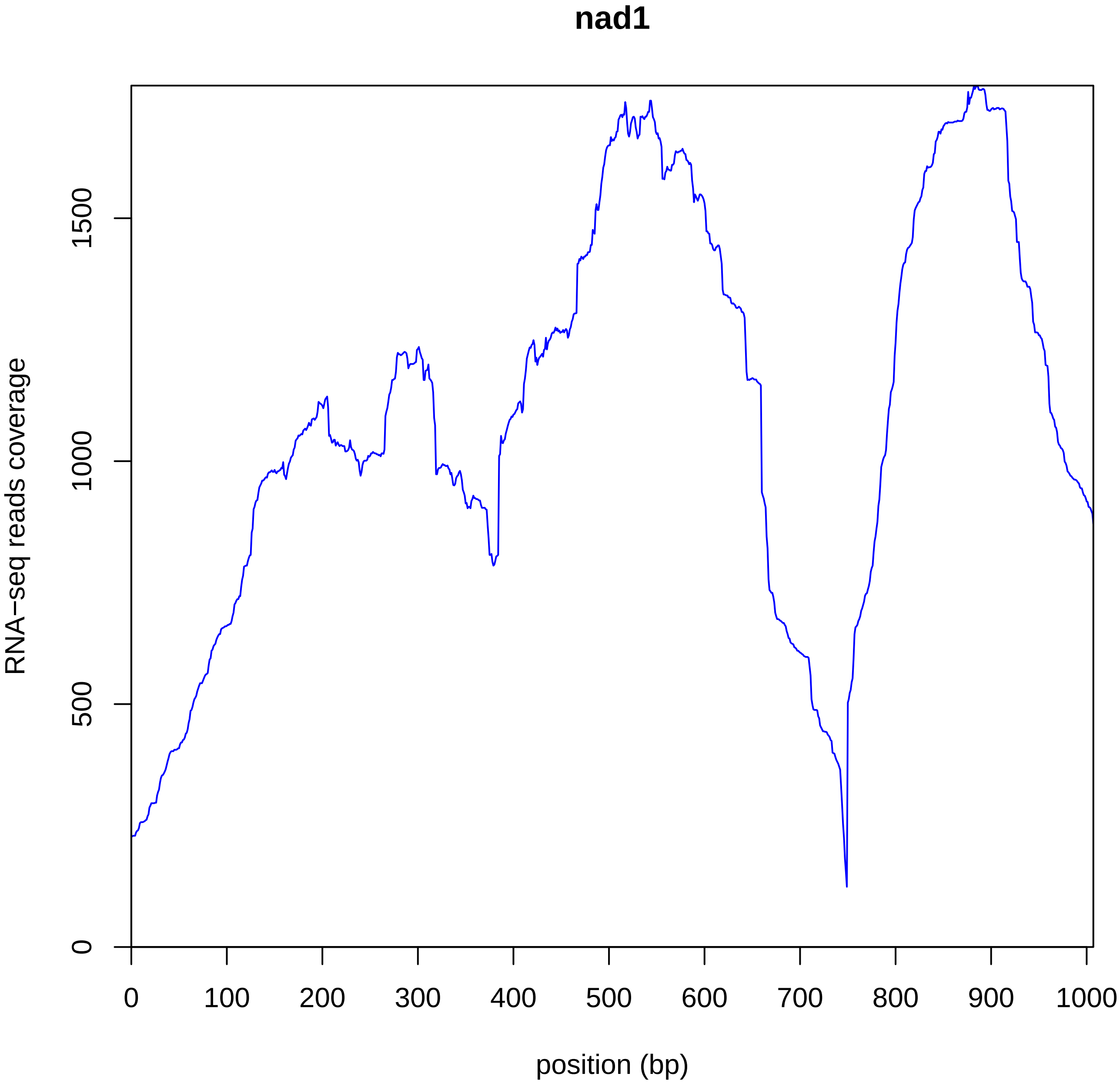
nad1
RNA−seq reads coverage
0
500
1000
1500
0
100
200
300
400
500
600
700
800
900
1000
position (bp)

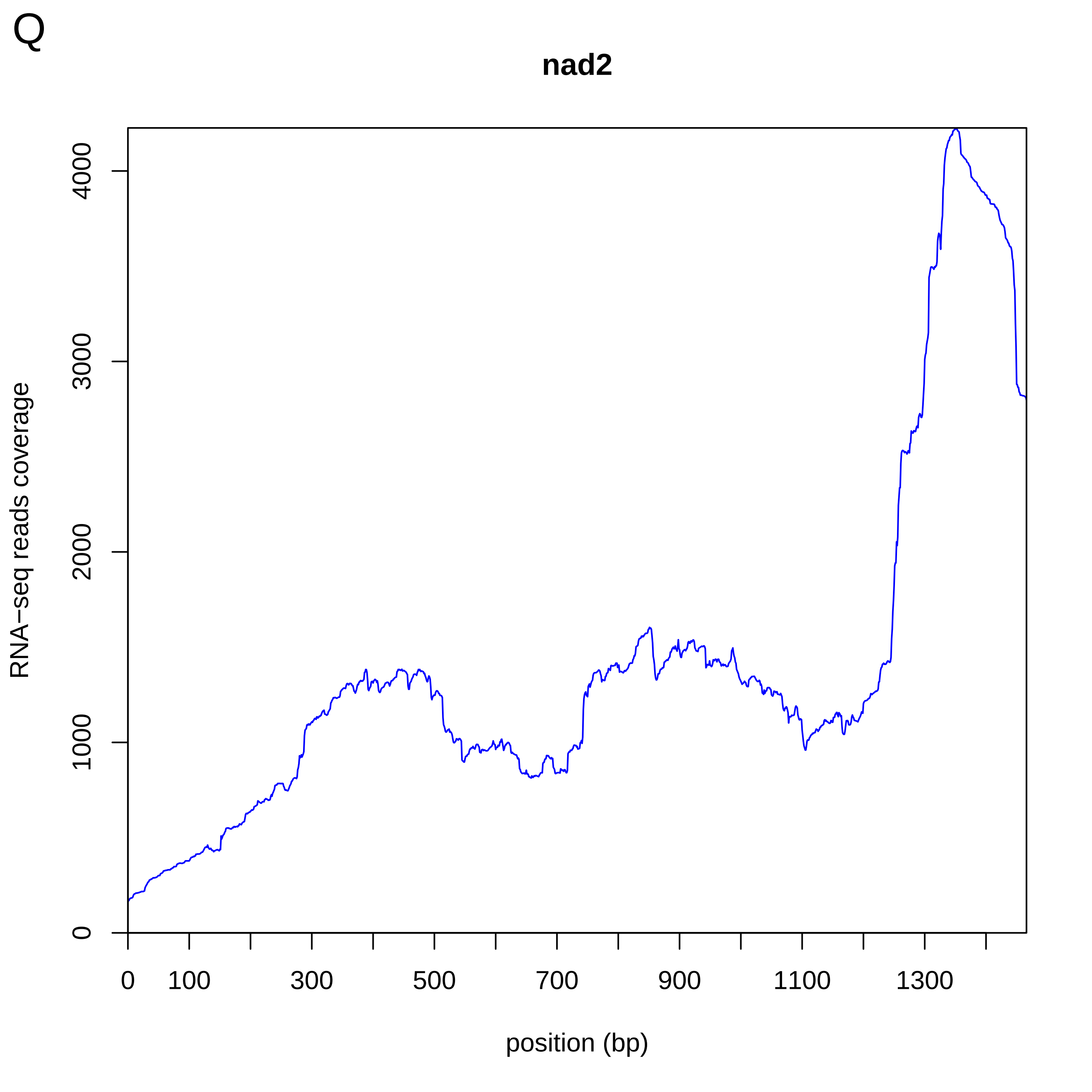
Q
nad2
RNA−seq reads coverage
0
1000
2000
3000
4000
0
100
300
500
700
900
1100
1300
position (bp)

R

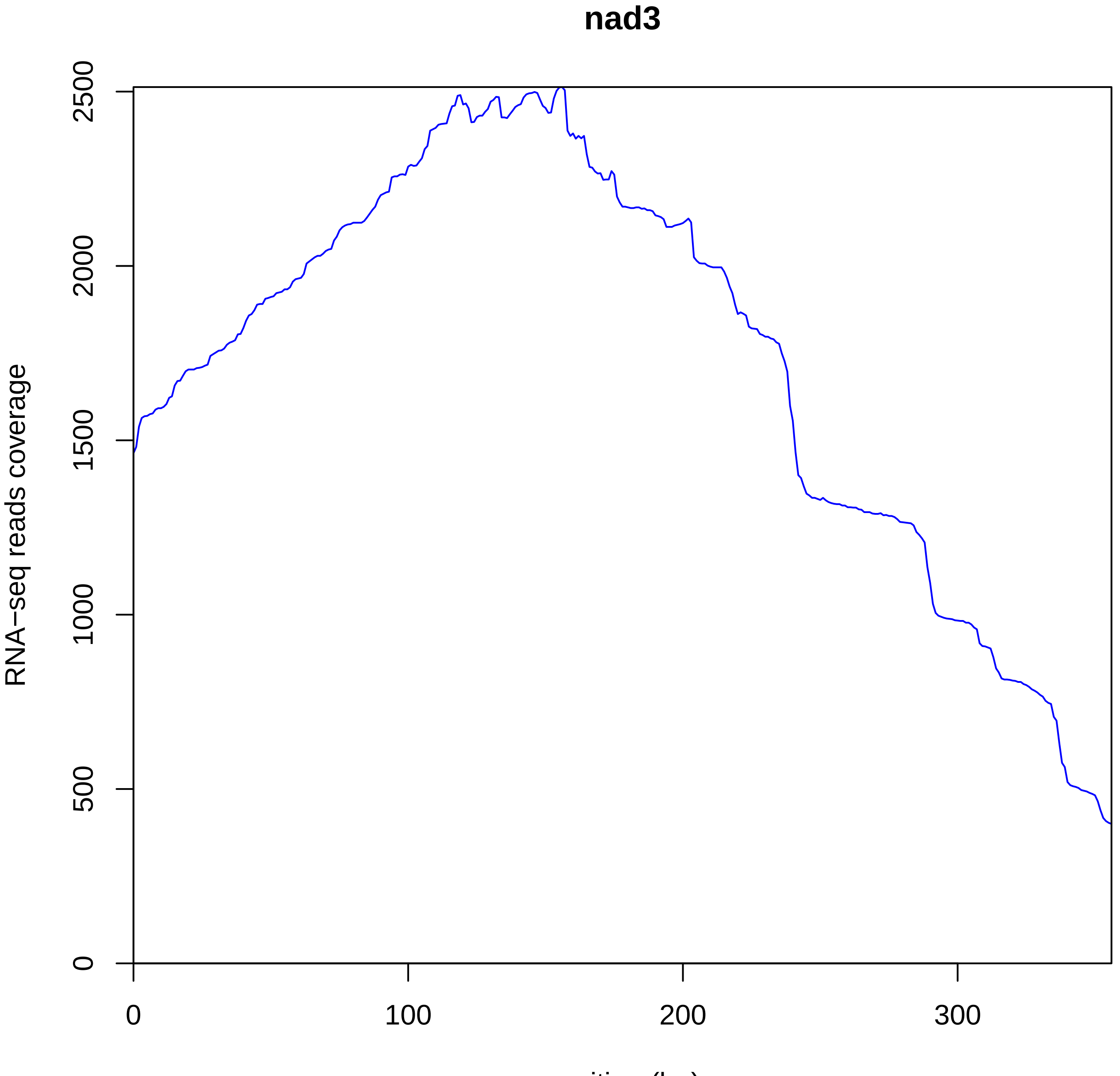
nad3
RNA−seq reads coverage
0
500
1000
1500
2000
2500
0
100
200
300
position (bp)

S

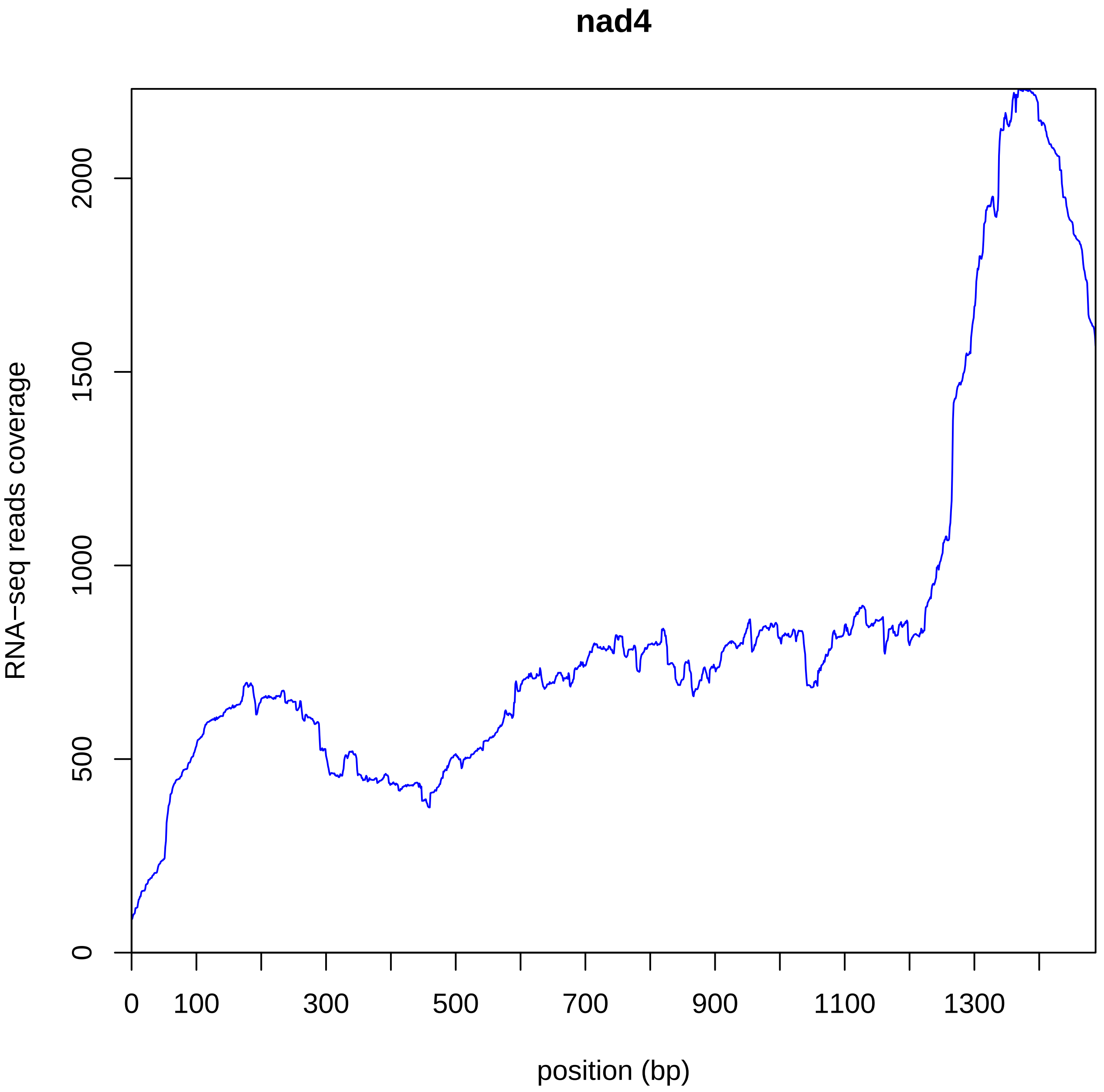
nad4
RNA−seq reads coverage
0
500
1000
1500
2000
0
100
300
500
700
900
1100
1300
position (bp)

T

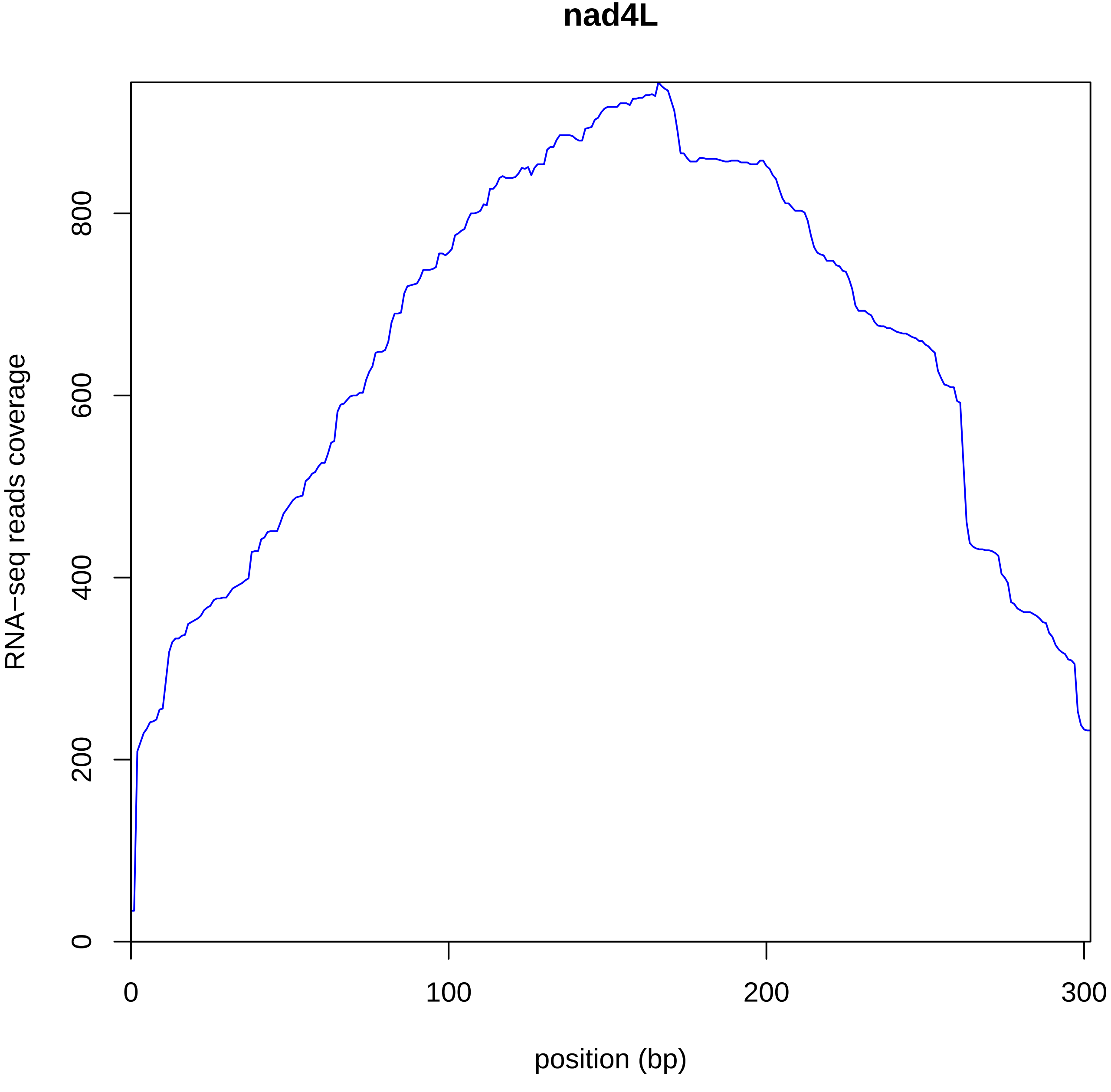
nad4L
RNA−seq reads coverage
0
200
400
600
800
0
100
200
300
position (bp)

U

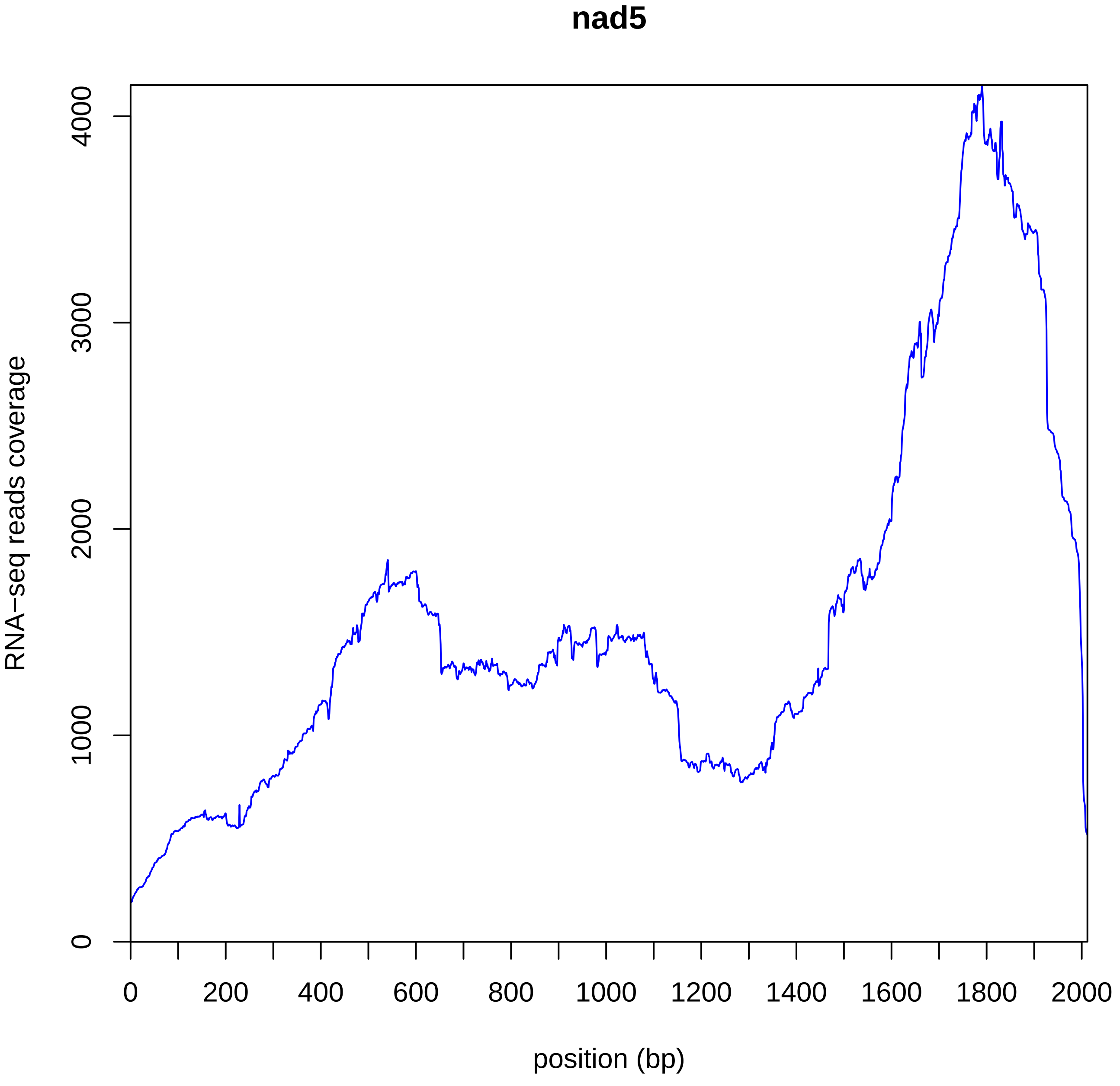

nad5
RNA−seq reads coverage
0
1000
2000
3000
4000
0
200
400
600
800
1000
1200
1400
1600
1800
2000
position (bp)

V

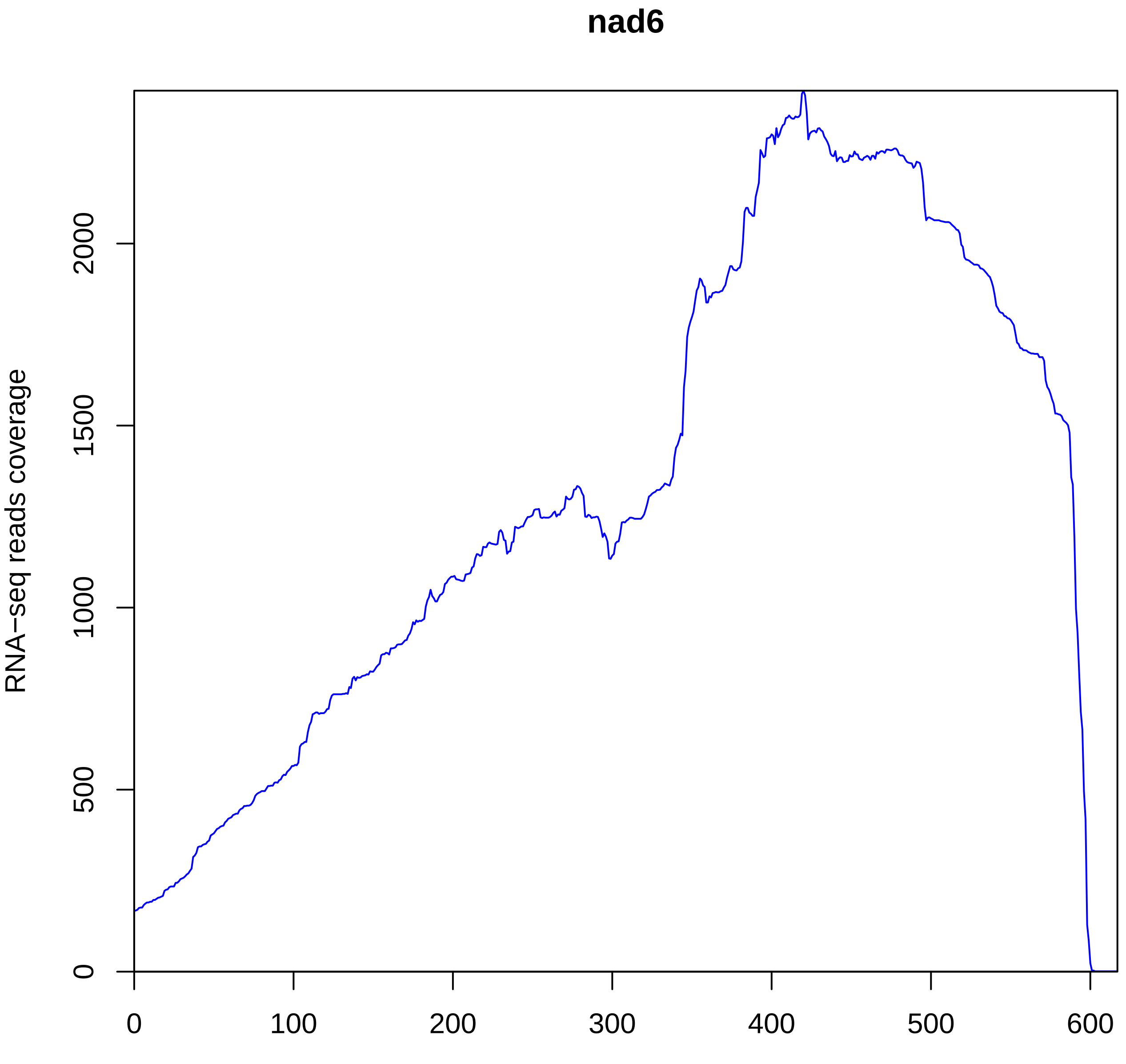
nad6
RNA−seq reads coverage
0
500
1000
1500
2000
0
100
200
300
400
500
600
position (bp)

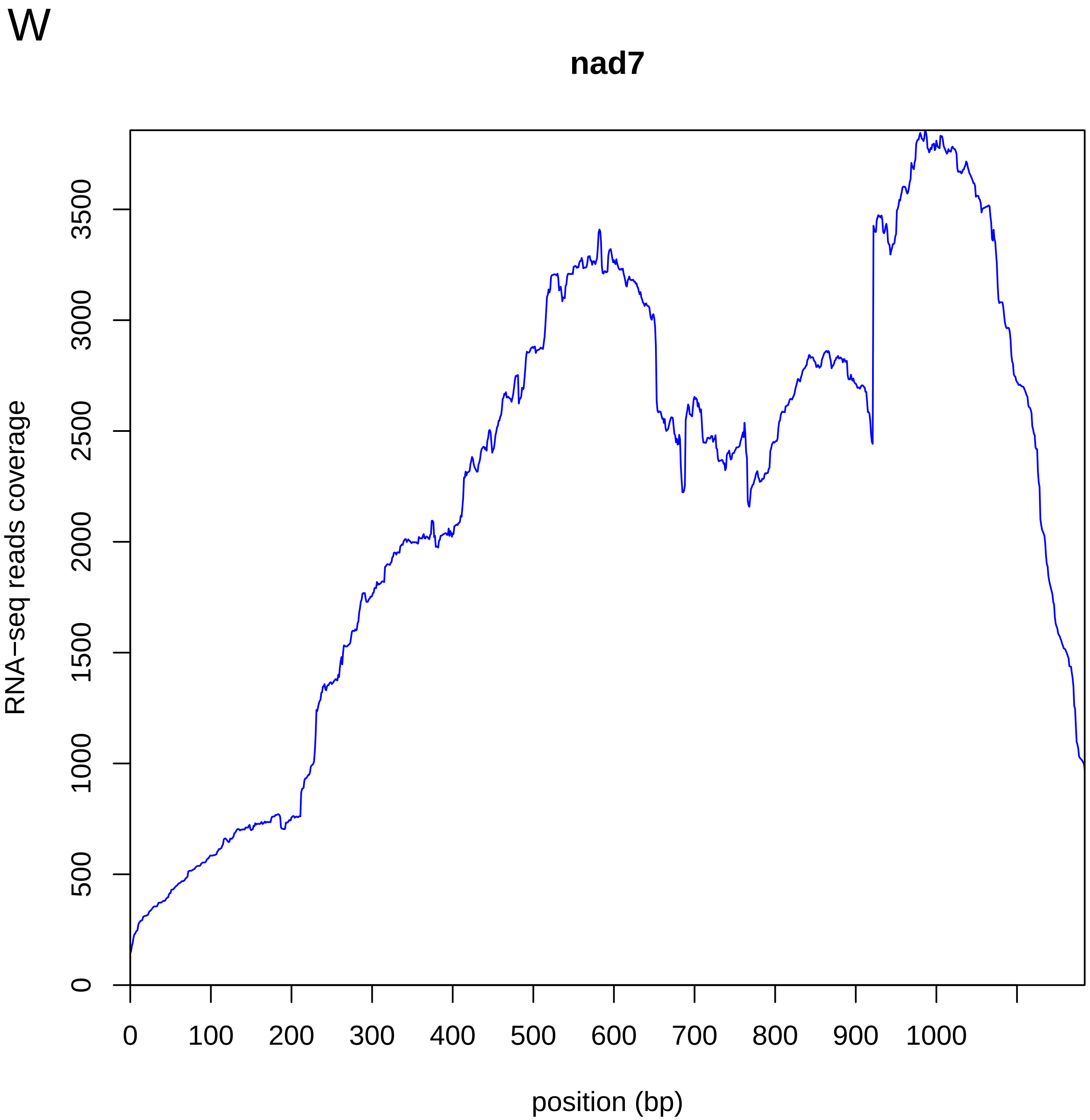

W
nad7
RNA−seq reads coverage
0
500
1000
1500
2000
2500
3000
3500
0
100
200
300
400
500
600
700
800
900
1000
position (bp)

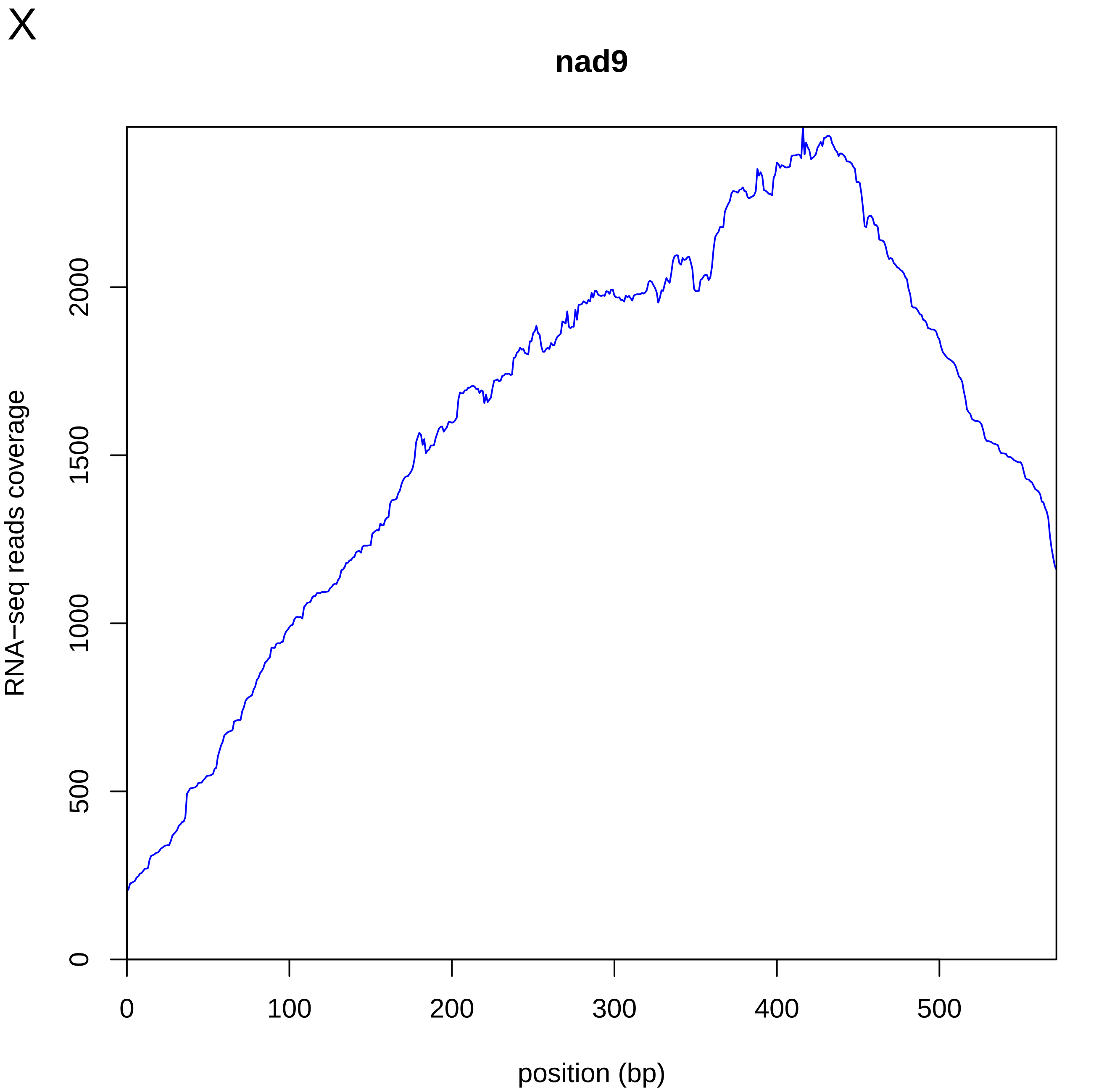
X
nad9
RNA−seq reads coverage
0
500
1000
1500
2000
0
100
200
300
400
500
position (bp)

Y

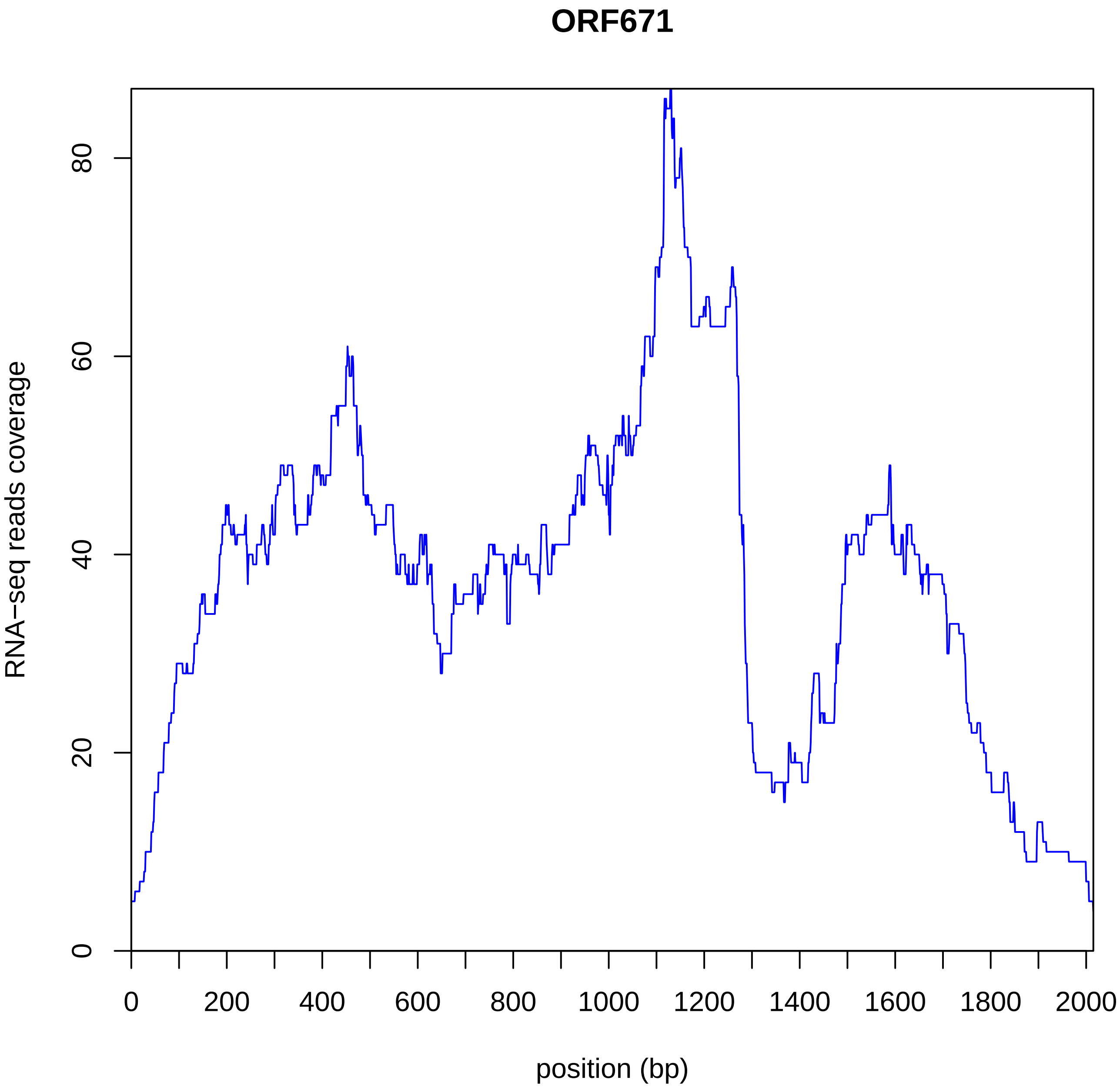
ORF671
RNA−seq reads coverage
0
20
40
60
80
0
200
400
600
800
1000
1200
1400
1600
1800
2000
position (bp)

Z

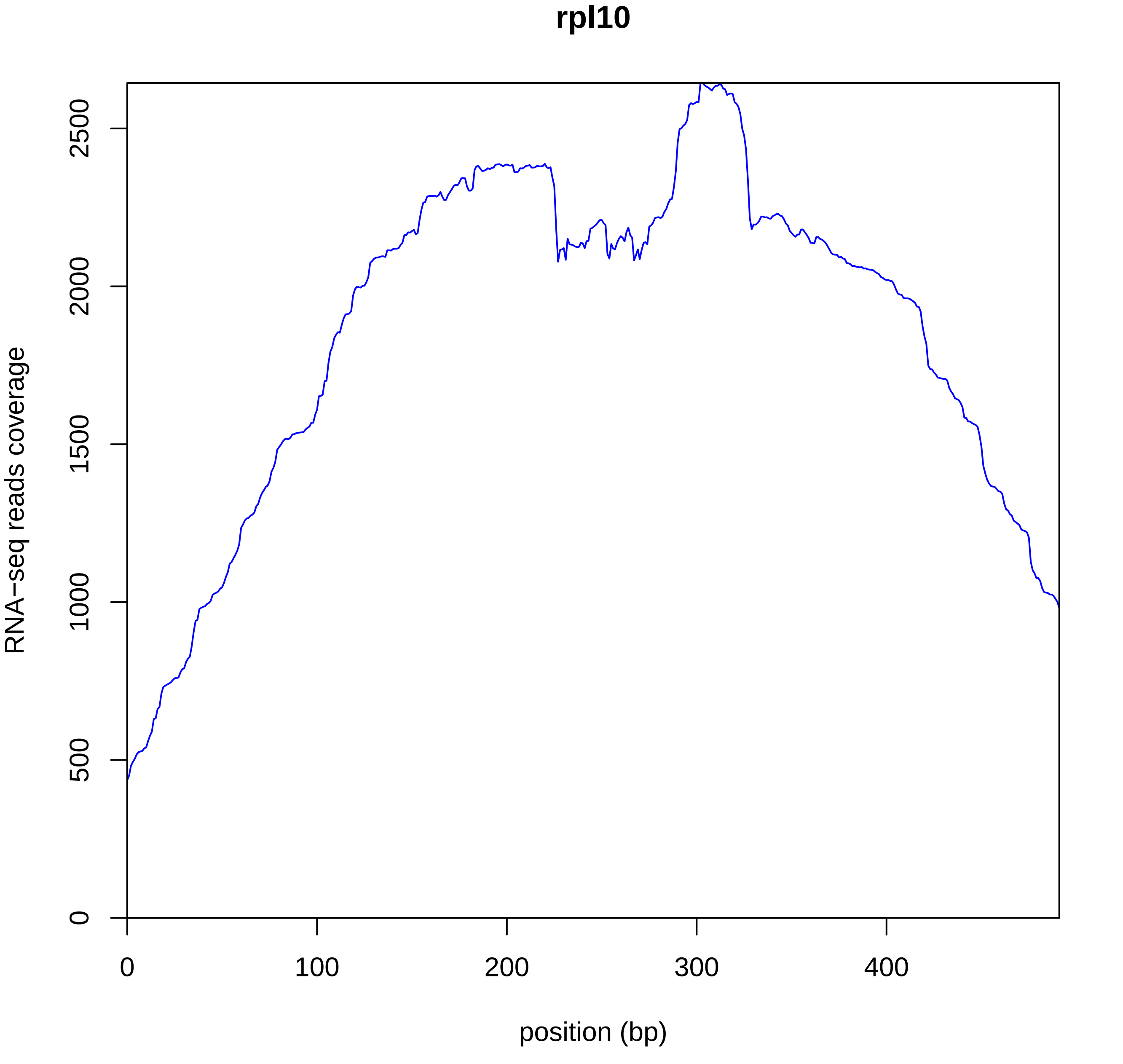
rpl10
RNA−seq reads coverage
0
500
1000
1500
2000
2500
position (bp)
0
100
200
300
400

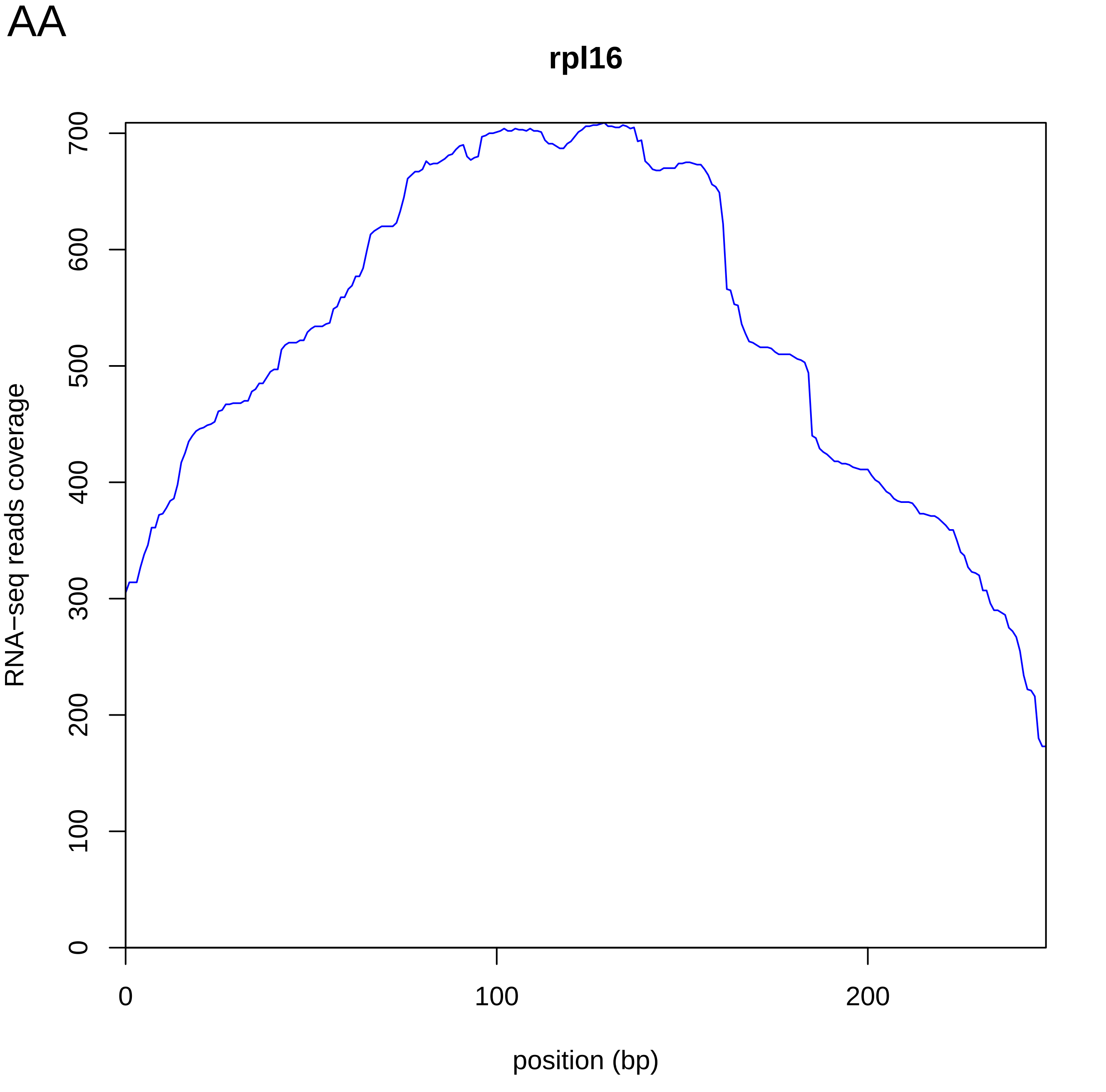
AA
rpl16
RNA−seq reads coverage
0
100
200
300
400
500
600
700
0
100
200
position (bp)

AB

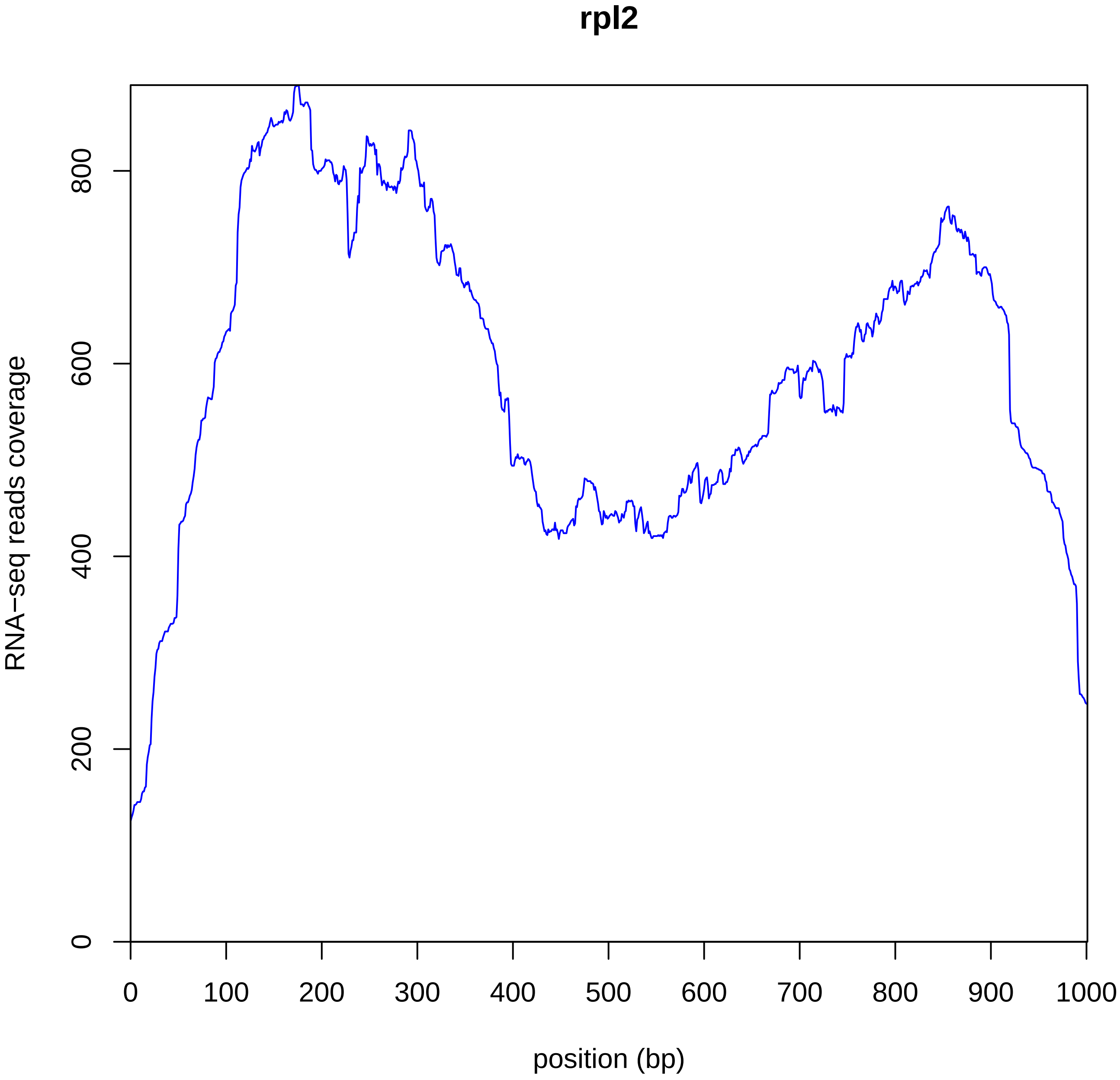
rpl2
RNA−seq reads coverage
0
200
400
600
800
0
100
200
300
400
500
600
700
800
900
1000
position (bp)

AC

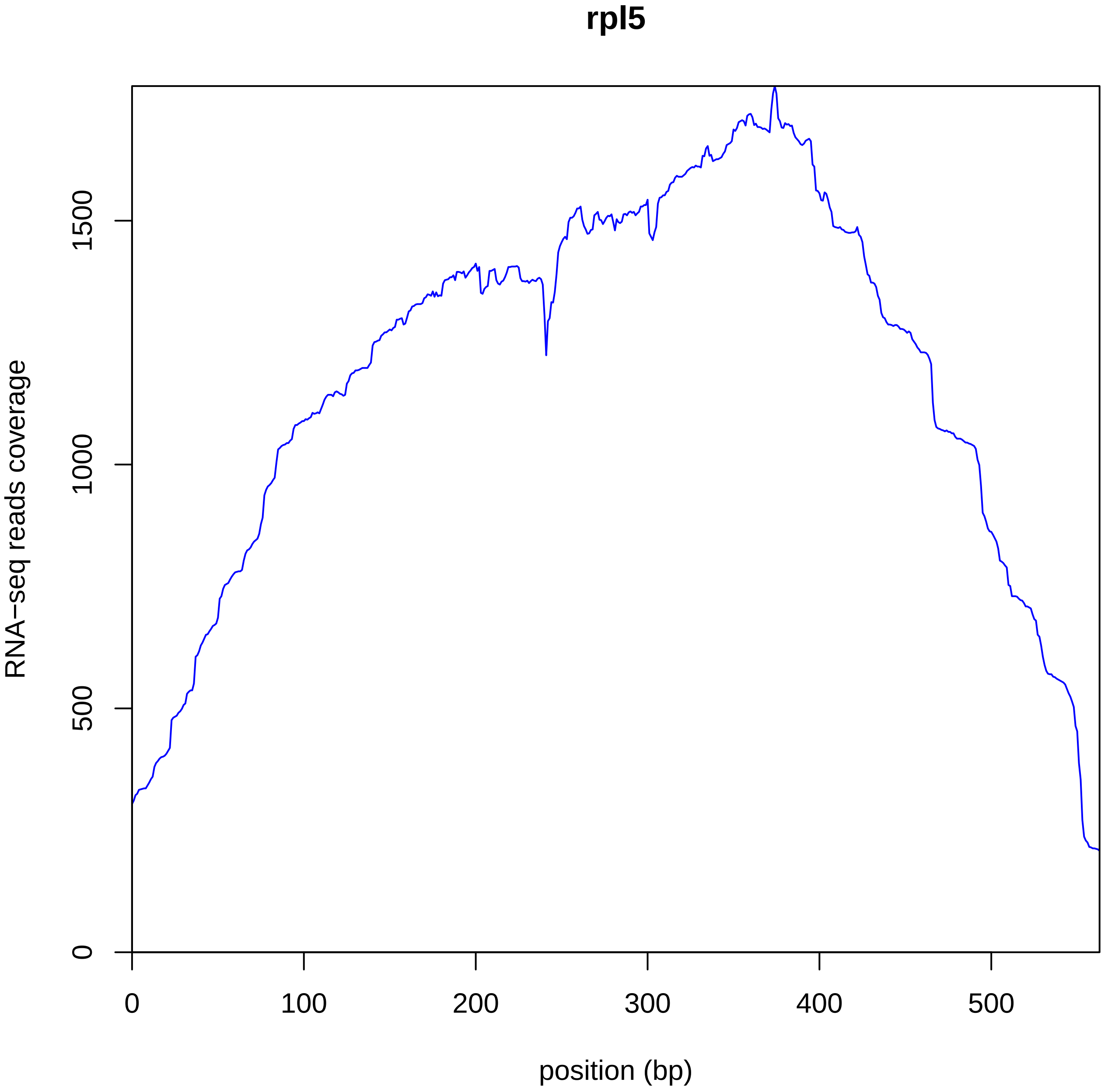
rpl5
RNA−seq reads coverage
0
500
1000
1500
0
100
200
300
400
500
position (bp)

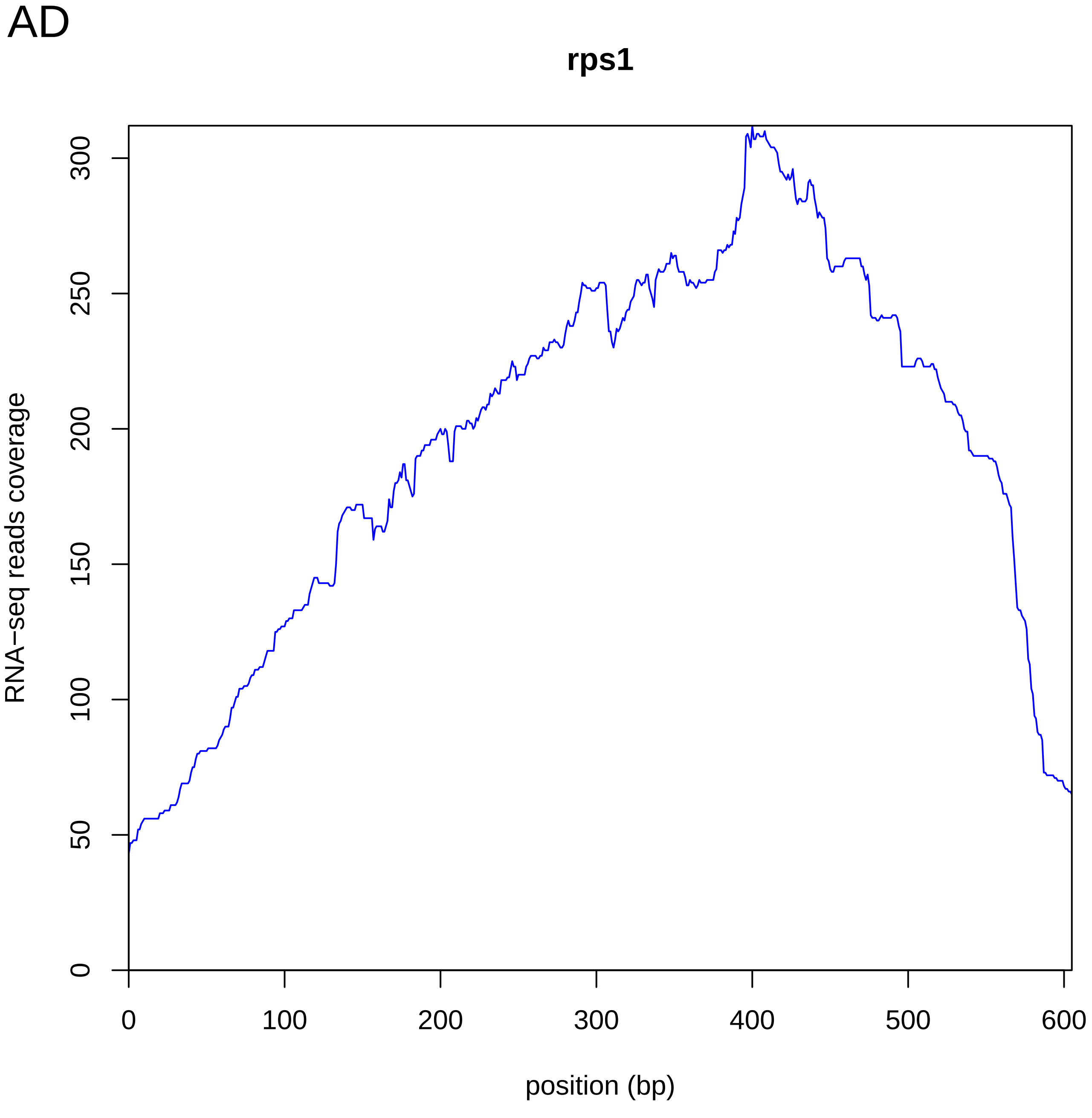

AD
rps1
RNA−seq reads coverage
0
50
100
150
200
250
300
0
100
200
300
400
500
600
position (bp)

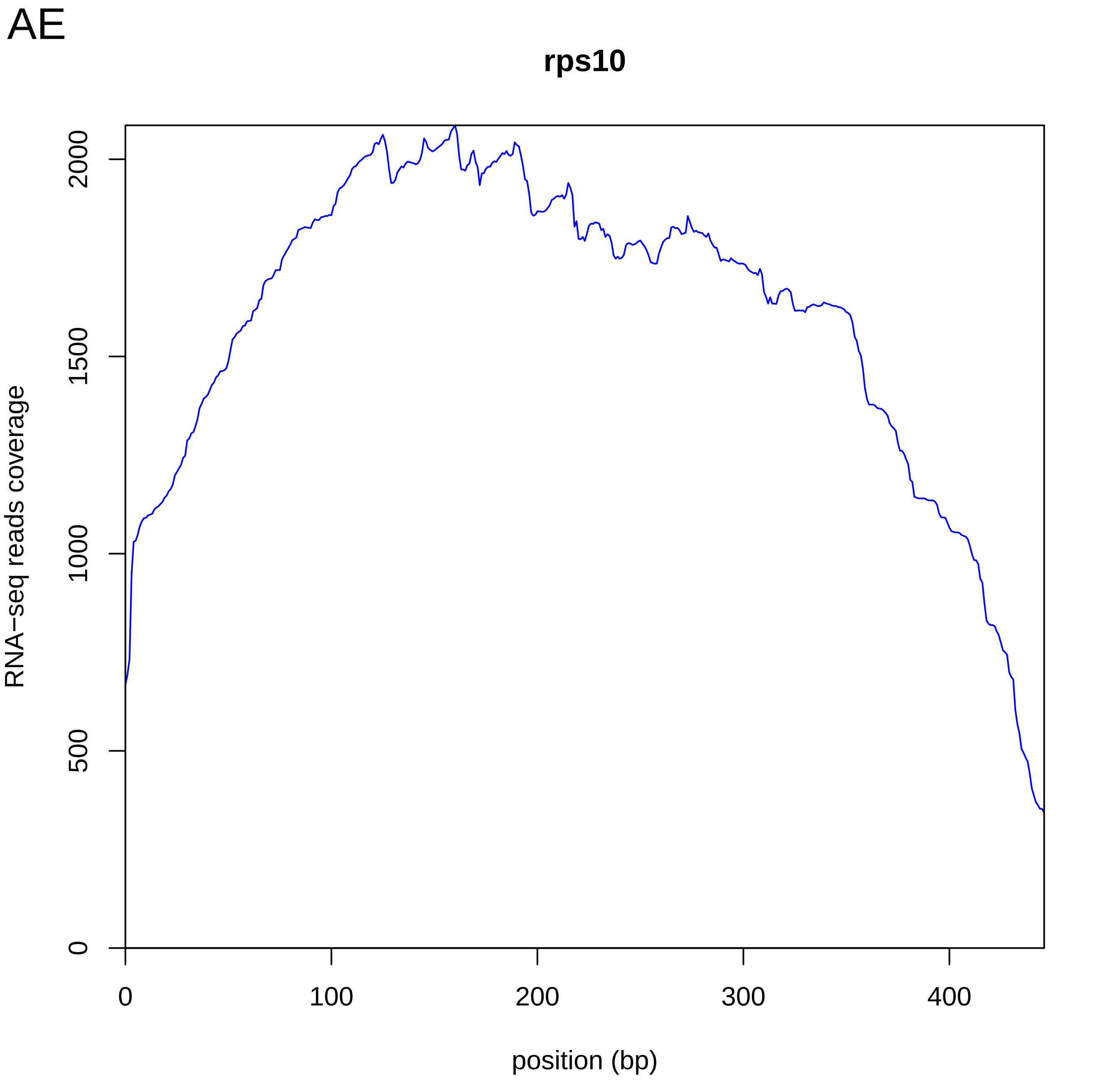
AE
rps10
RNA−seq reads coverage
0
500
1000
1500
2000
0
100
200
300
400
position (bp)

AF

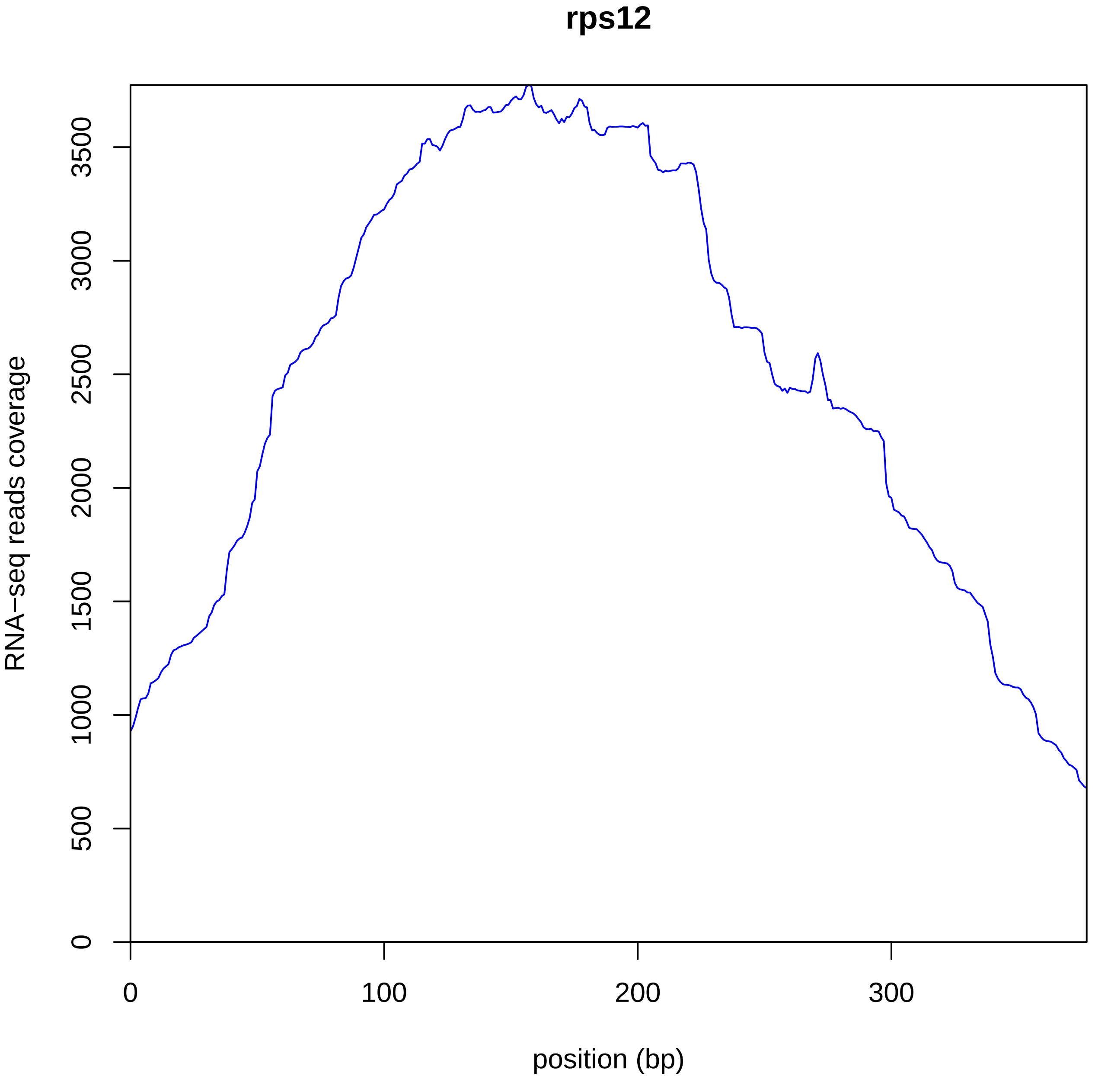
rps12
RNA−seq reads coverage
position (bp)
0
500
1000
1500
2000
2500
3000
3500
0
100
200
300

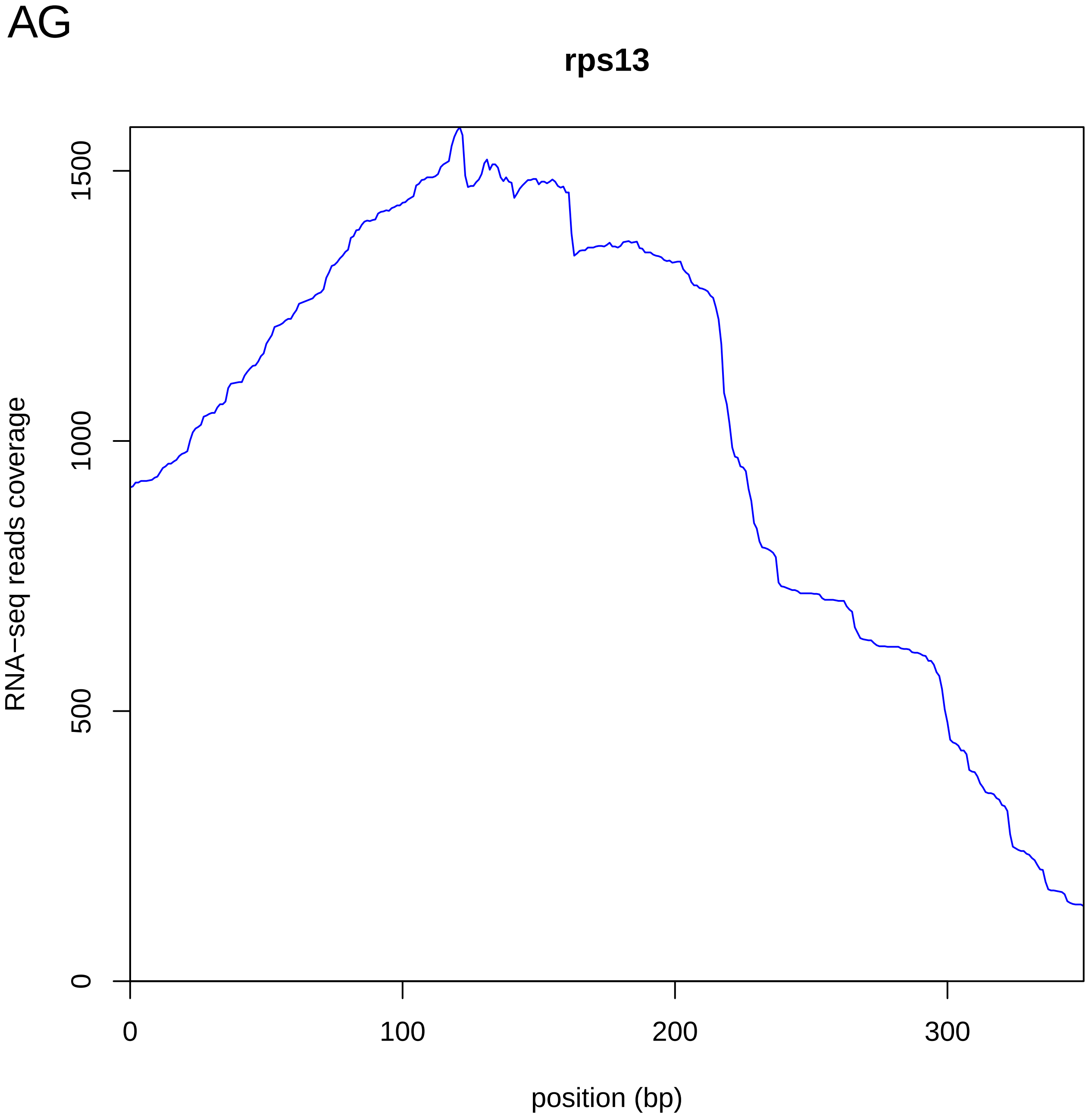
AG
rps13
RNA−seq reads coverage
0
500
1000
1500
0
100
200
300
position (bp)

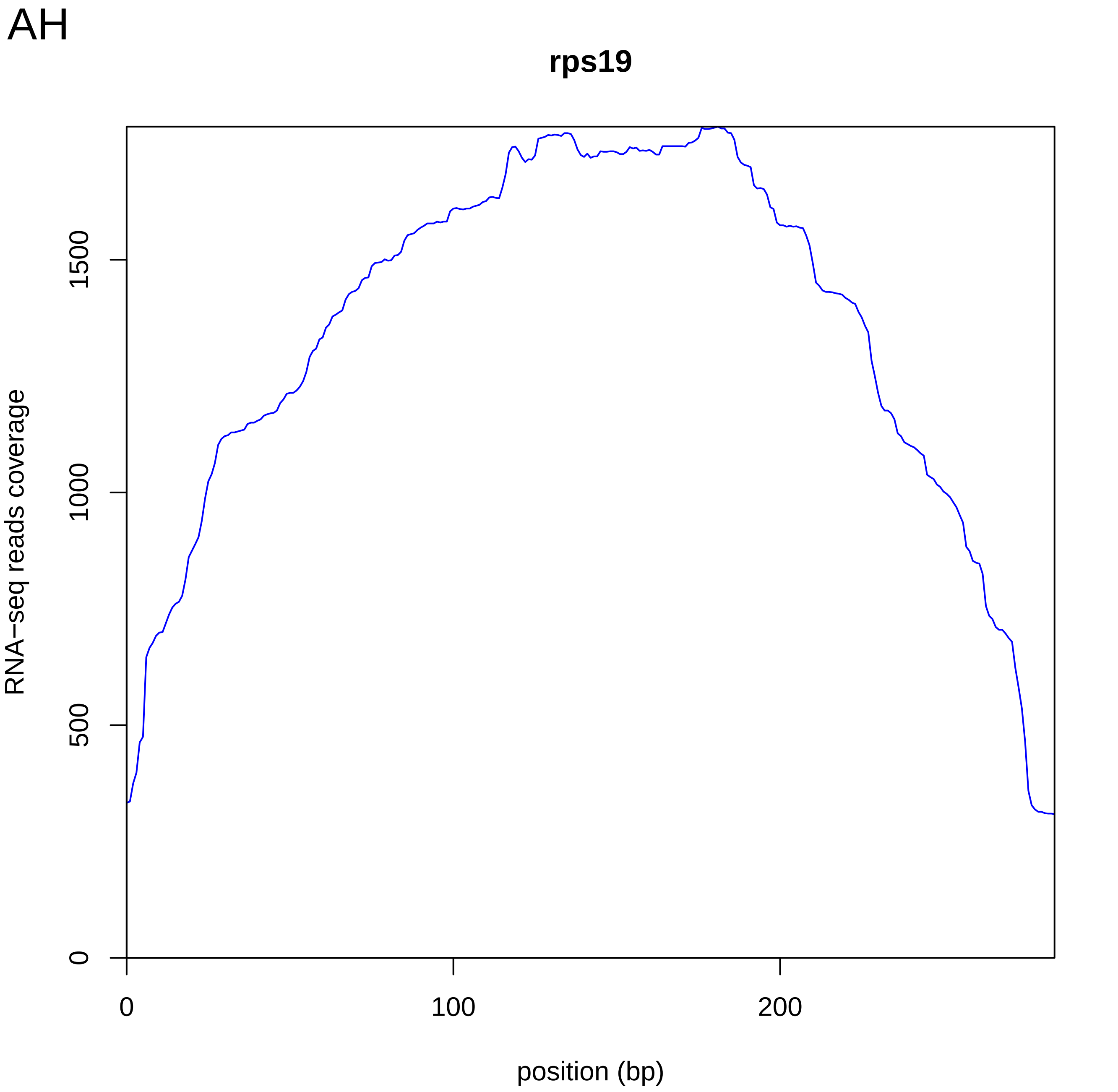
AH
rps19
RNA−seq reads coverage
0
500
1000
1500
0
100
200
position (bp)

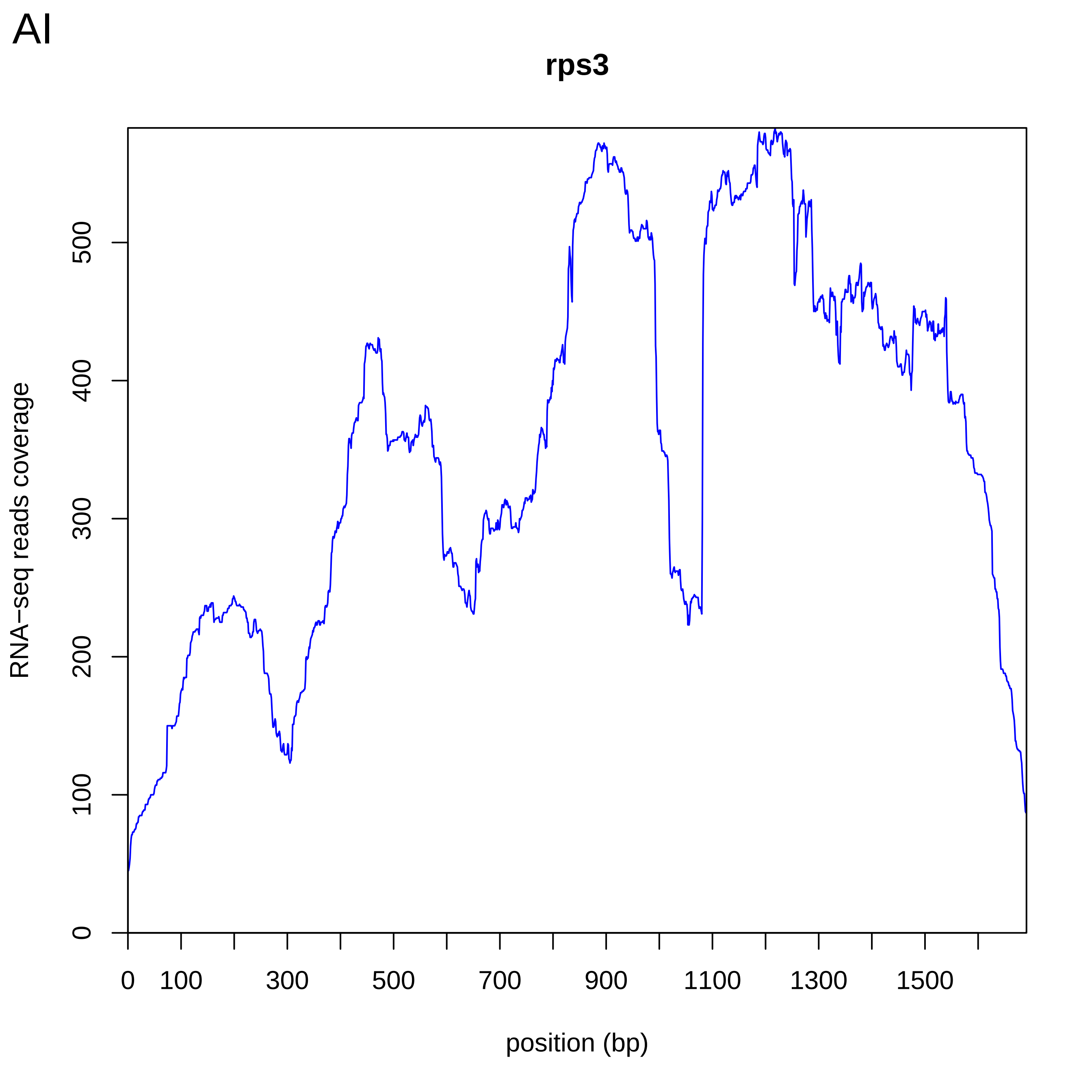
AI
rps3
RNA−seq reads coverage
0
100
200
300
400
500
0
100
300
500
700
900
1100
1300
1500
position (bp)

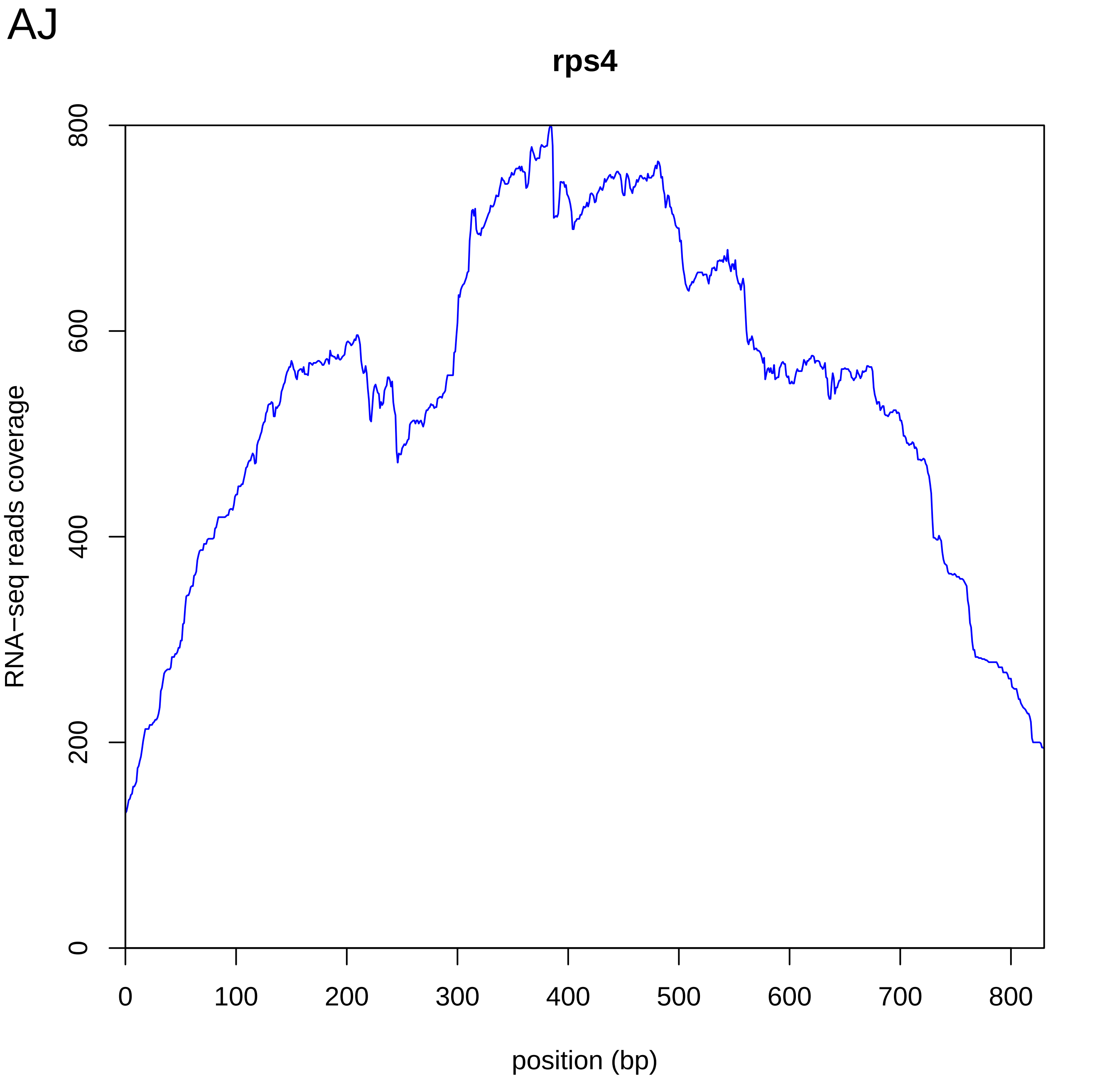
AJ
rps4
RNA−seq reads coverage
0
200
400
600
800
0
100
200
300
400
500
600
700
800
position (bp)

AK

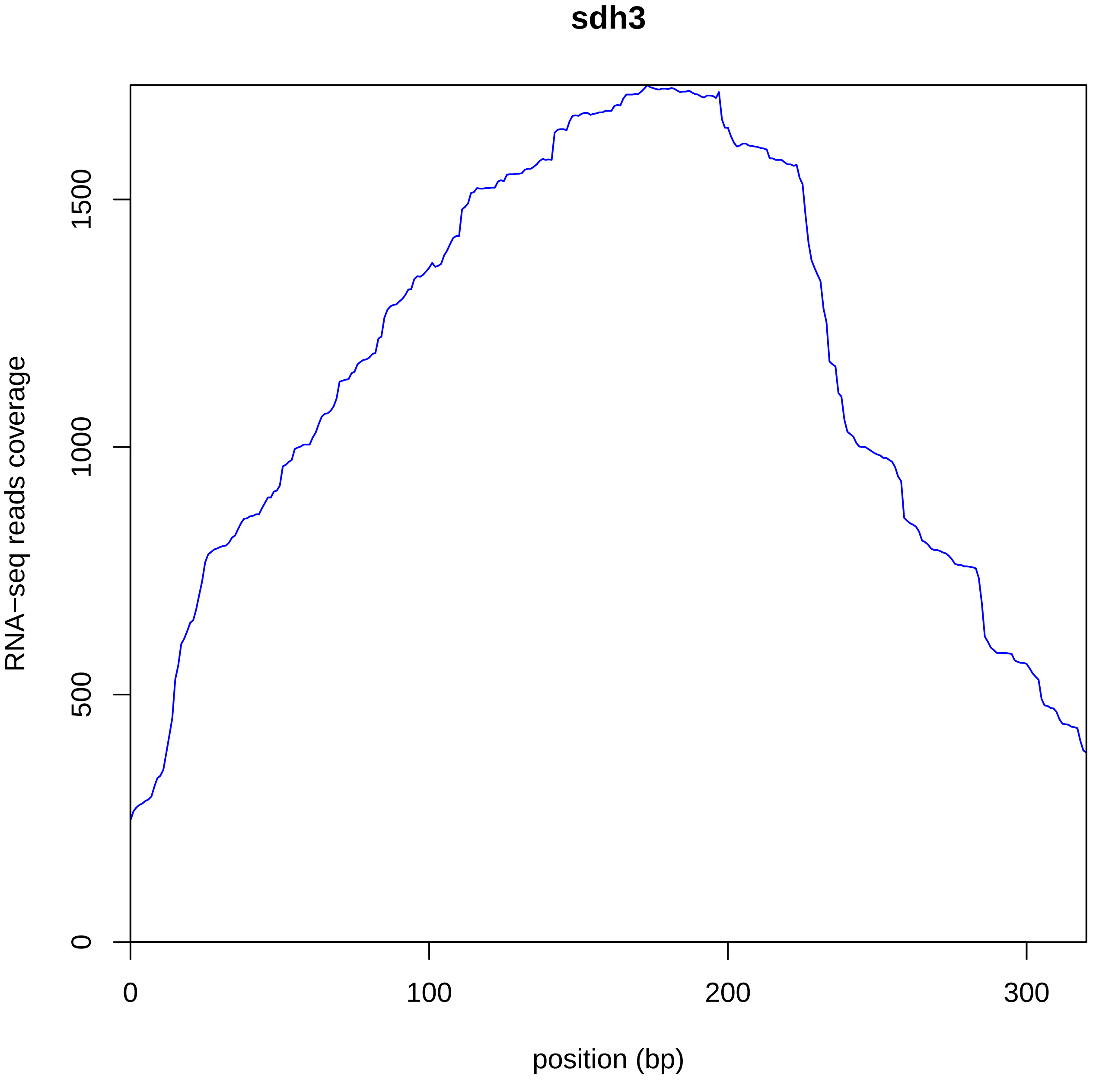

sdh3
RNA−seq reads coverage
position (bp)
0
500
1000
1500
0
100
200
300

AL

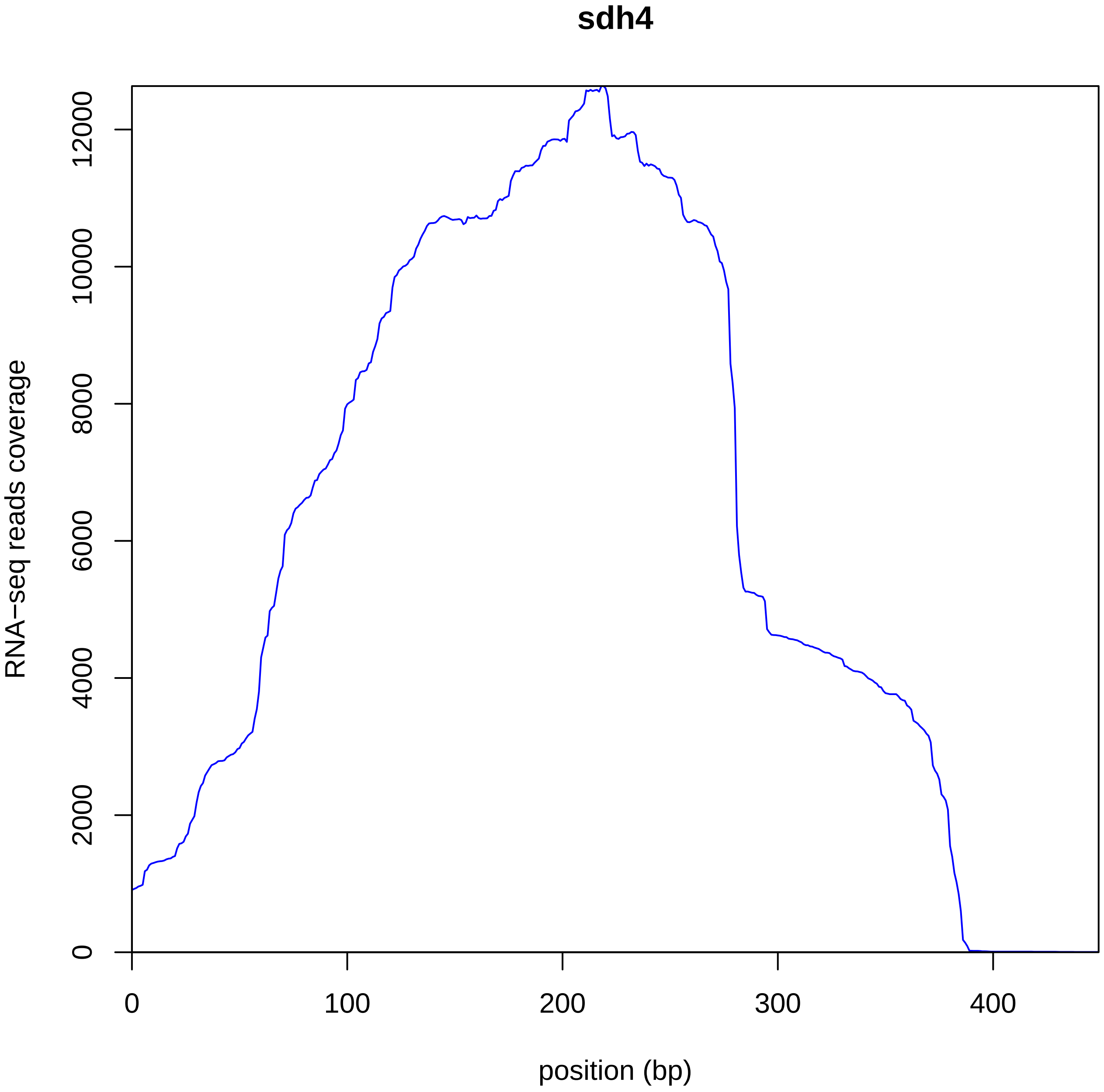

sdh4
RNA−seq reads coverage
0
2000
4000
6000
8000
10000
12000
0
100
200
300
400
position (bp)
