## Supplementary Figure S5 for "Mitochondrial genome of non-photosynthetic mycoheterotrophic plant *Hypopitys monotropa*,its structure, gene expression and RNA editing"

Supplementary figure 5.

Phylogenetic trees inferred from ML analysis of single mitochondrial genes. Protein coding genes that were annotated in mitochondrial genomes of *Hypopitys monotropa, Vaccinium macrocarpon* and all main evolutionary lineages of flowering plants are included in the analysis.

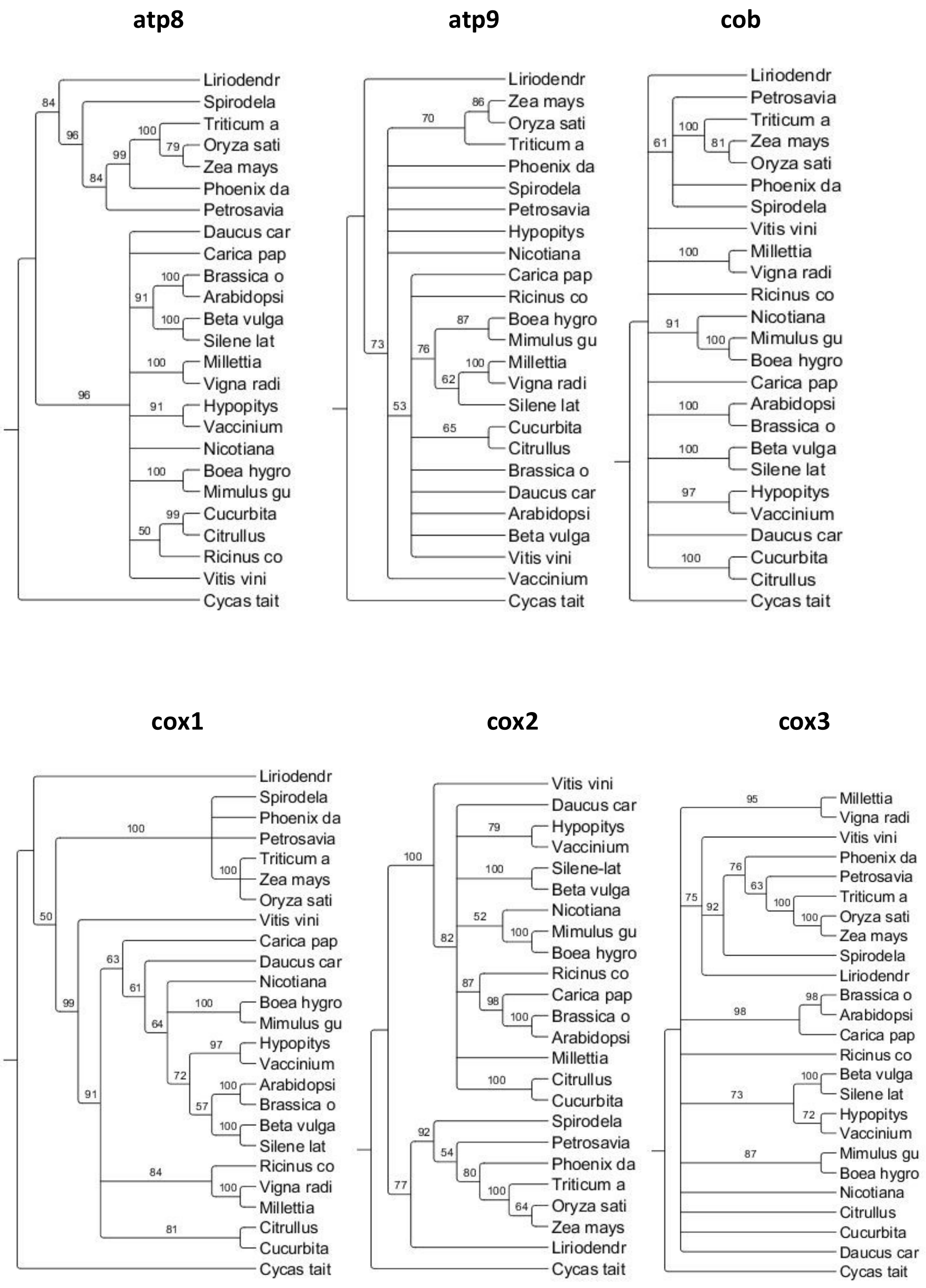

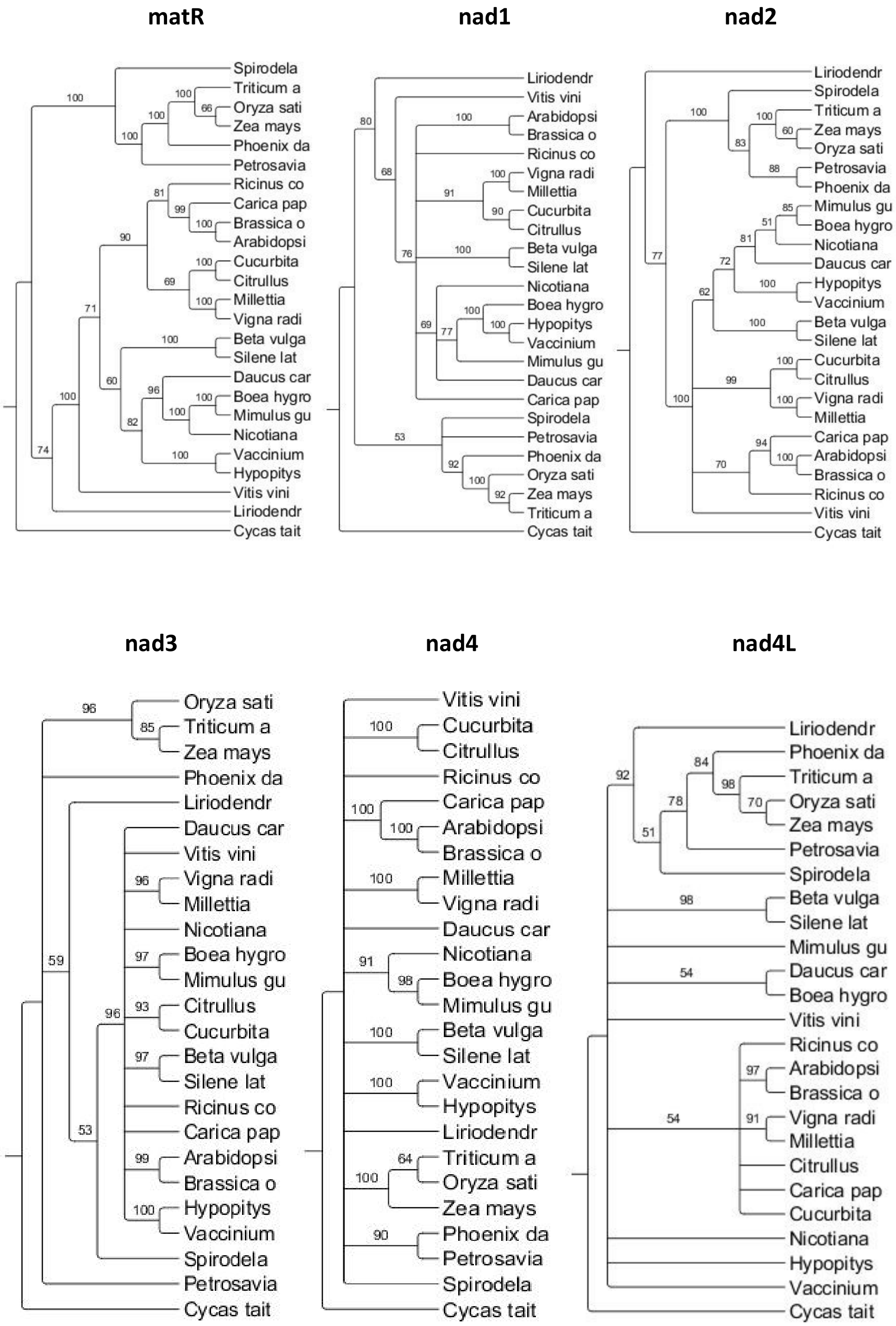
matR
Spirodela
Triticum a
Oryza sati
Zea mays
Phoenix da
Petrosavia
Ricinus co
Carica pap
Brassica o
Arabidopsi
Cucurbita
Citrullus
Millettia
Vigna radi
Beta vulga
Silene lat
Daucus car
Boea hygro
Mimulus gu
Nicotiana
Vaccinium
Hypopitys
Vitis vini
Liriodendr
Cycas tait
nad1
Liriodendr
Vitis vini
Arabidopsi
Brassica o
Ricinus co
Vigna radi
Millettia
Cucurbita
Citrullus
Beta vulga
Silene lat
Nicotiana
Boea hygro
Hypopitys
Vaccinium
Mimulus gu
Daucus car
Carica pap
Spirodela
Petrosavia
Phoenix da
Oryza sati
Zea mays
Triticum a
Cycas tait
nad2
Liriodendr
Spirodela
Triticum a
Zea mays
Oryza sati
Petrosavia
Phoenix da
Mimulus gu
Boea hygro
Nicotiana
Daucus car
Hypopitys
Vaccinium
Beta vulga
Silene lat
Cucurbita
Citrullus
Vigna radi
Millettia
Carica pap
Arabidopsi
Brassica o
Ricinus co
Vitis vini
Cycas tait
nad3
Oryza sati
Triticum a
Zea mays
Phoenix da
Liriodendr
Daucus car
Vitis vini
Vigna radi
Millettia
Nicotiana
Boea hygro
Mimulus gu
Citrullus
Cucurbita
Beta vulga
Silene lat
Ricinus co
Carica pap
Arabidopsi
Brassica o
Hypopitys
Vaccinium
Spirodela
Petrosavia
Cycas tait
nad4
Vitis vini
Cucurbita
Citrullus
Ricinus co
Carica pap
Arabidopsi
Brassica o
Millettia
Vigna radi
Daucus car
Nicotiana
Boea hygro
Mimulus gu
Beta vulga
Silene lat
Vaccinium
Hypopitys
Liriodendr
Triticum a
Oryza sati
Zea mays
Phoenix da
Petrosavia
Spirodela
Cycas tait
nad4L
Liriodendr
Phoenix da
Triticum a
Oryza sati
Zea mays
Petrosavia
Spirodela
Beta vulga
Silene lat
Mimulus gu
Daucus car
Boea hygro
Vitis vini
Ricinus co
Arabidopsi
Brassica o
Vigna radi
Millettia
Citrullus
Carica pap
Cucurbita
Nicotiana
Hypopitys
Vaccinium
Cycas tait

**nad5**

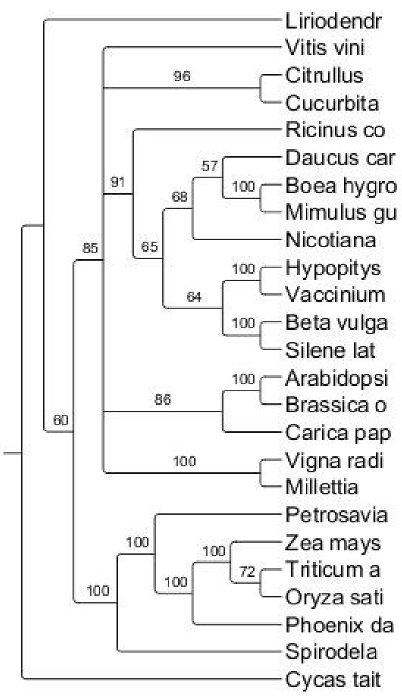


**nad6**

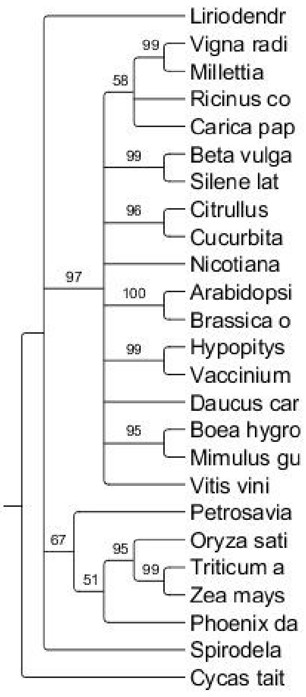


**nad7**

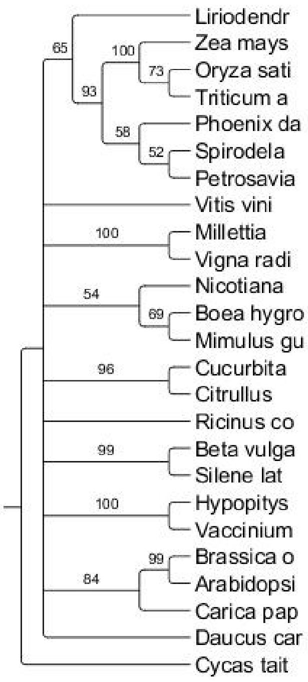


**nad9**

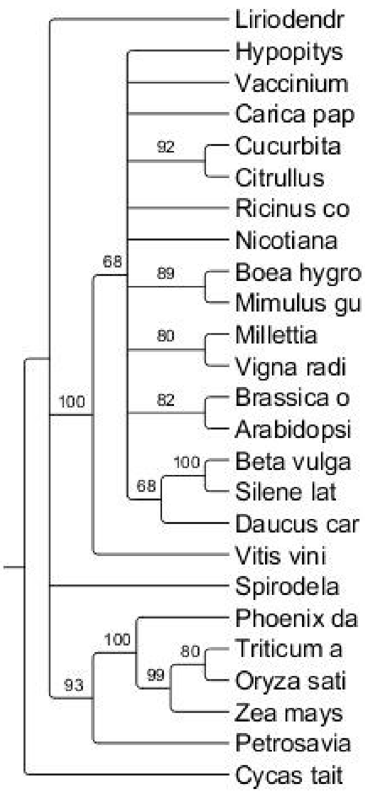


**rpl16**

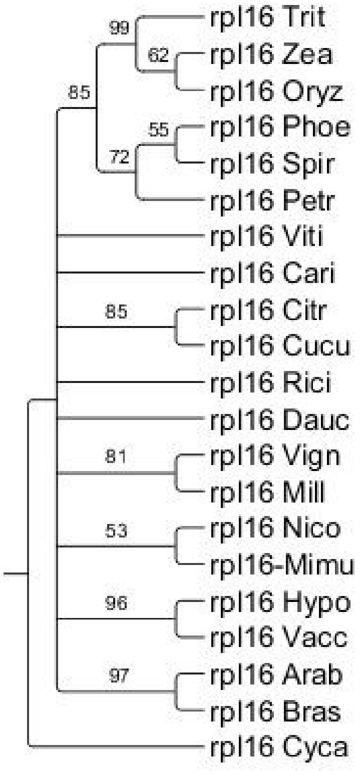


**rps1**

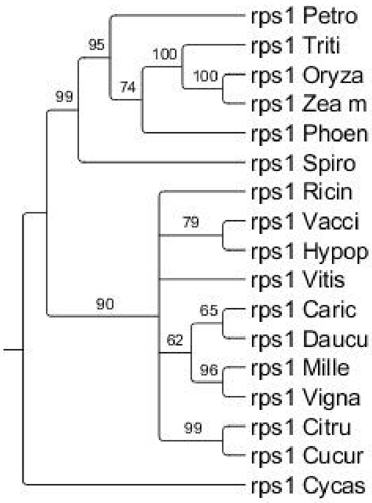

**rps3**

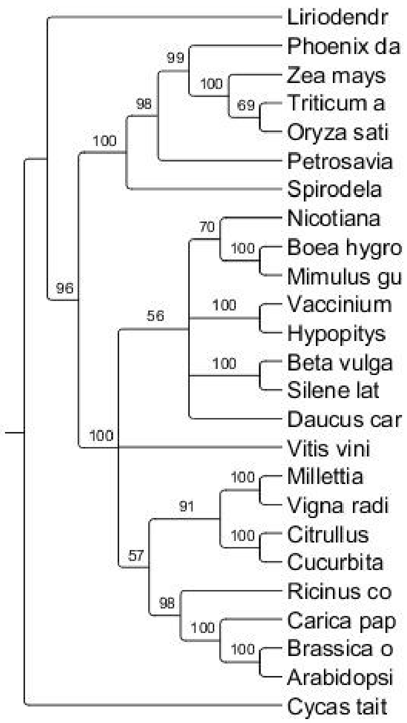


**rps12**

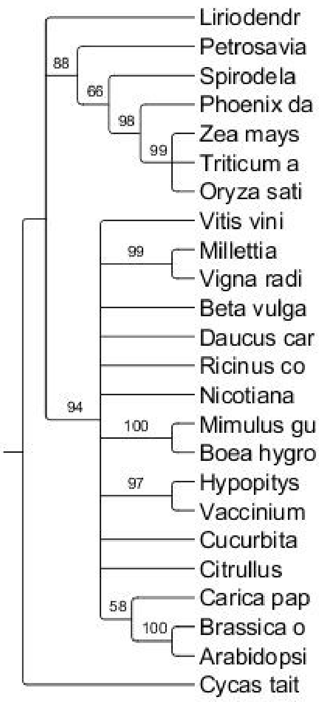


**rps14**

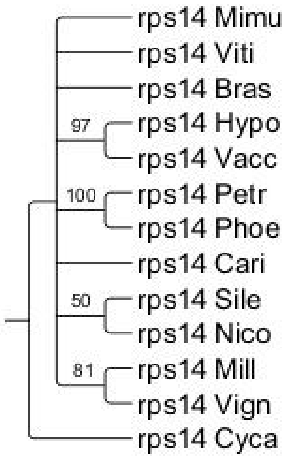


**rps19**

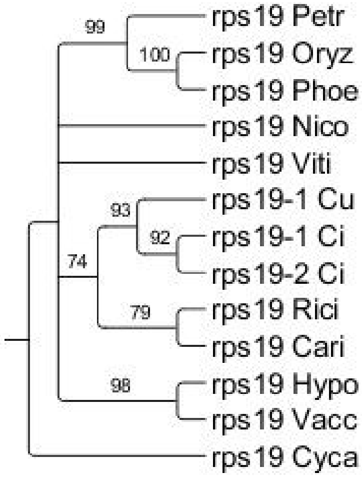


**sdh4**

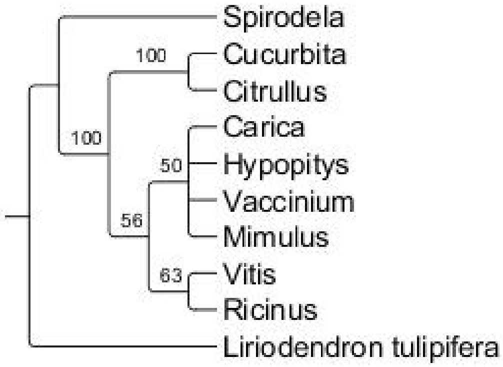


**rpl2**

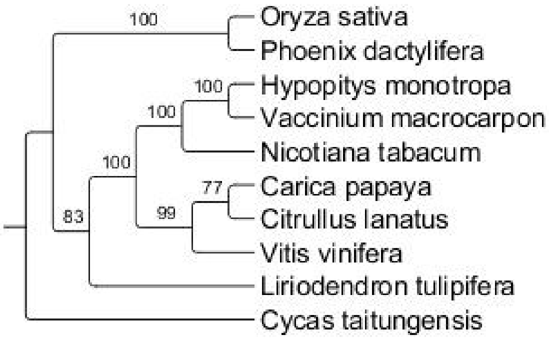

**atp1**

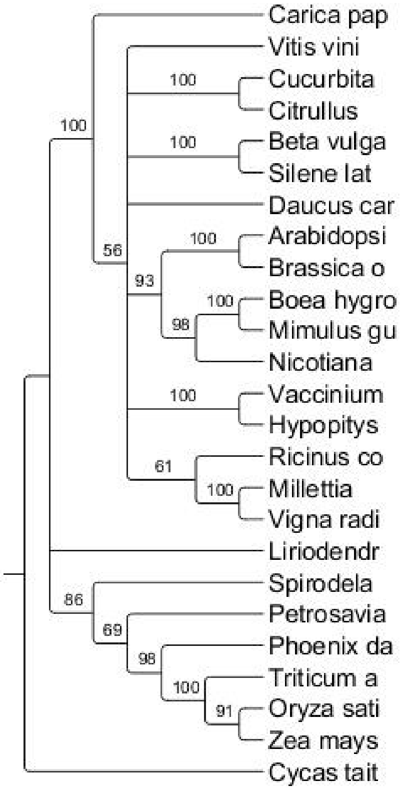


**atp4**

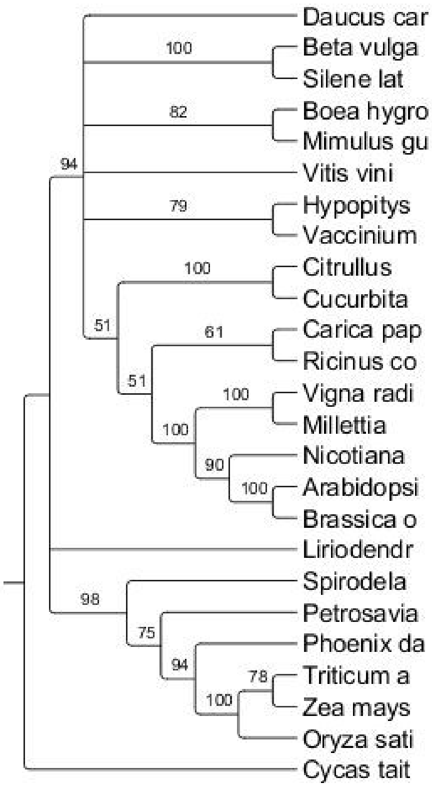


**mttB**

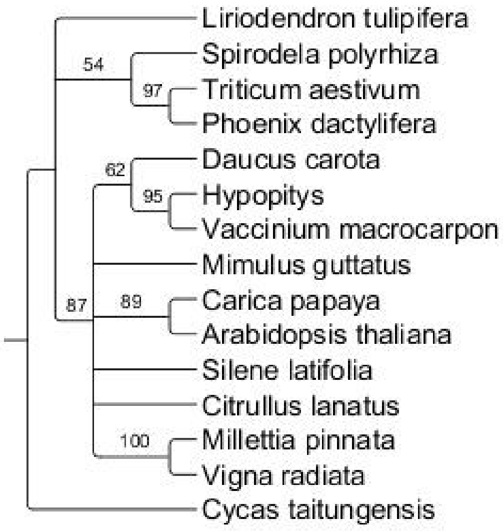


**ccmFc**

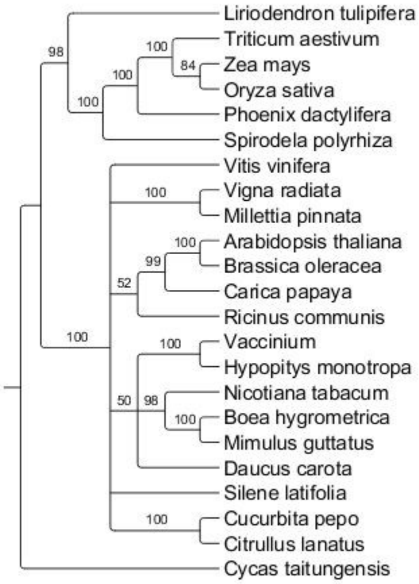

**ccmB**

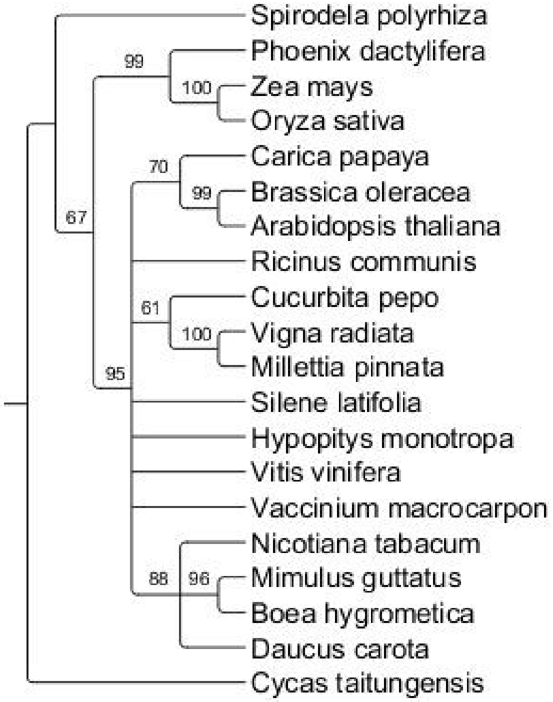


**ccmFn**

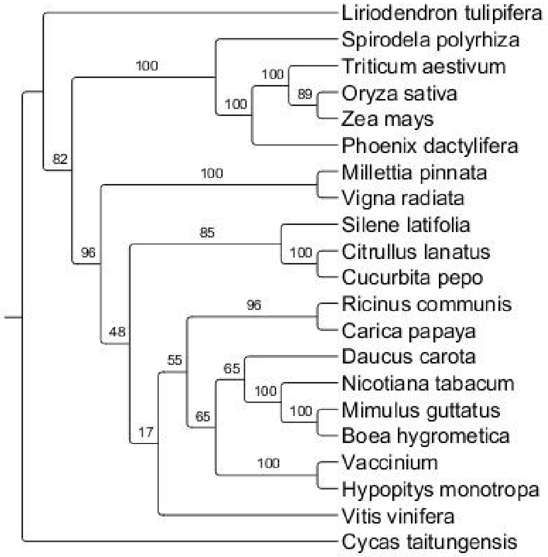


**rpl5**

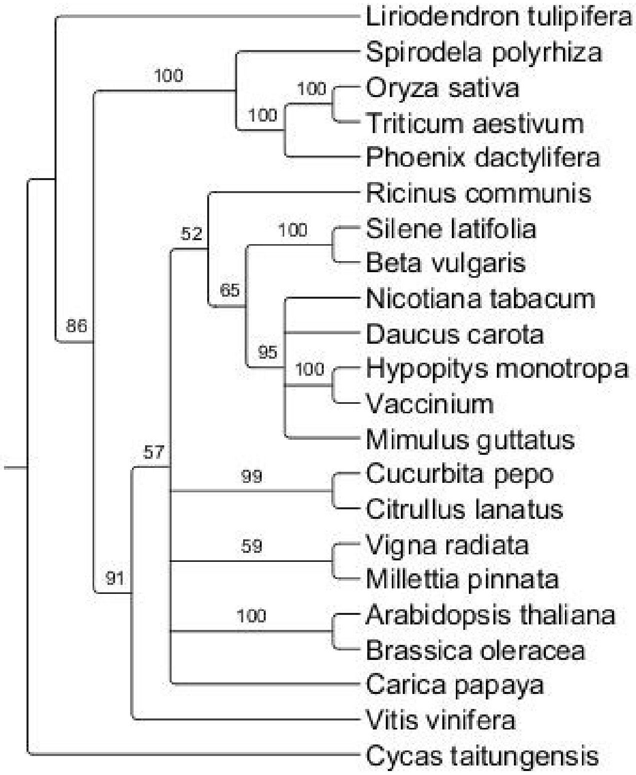


**rps10**

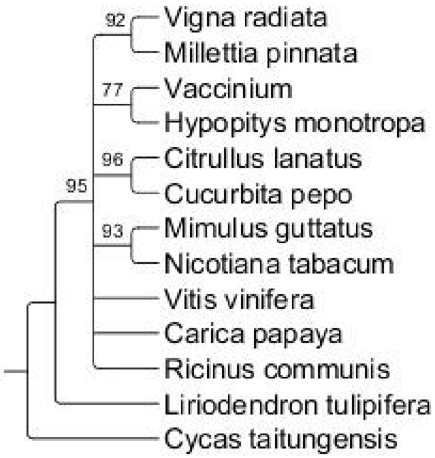

**rps4**

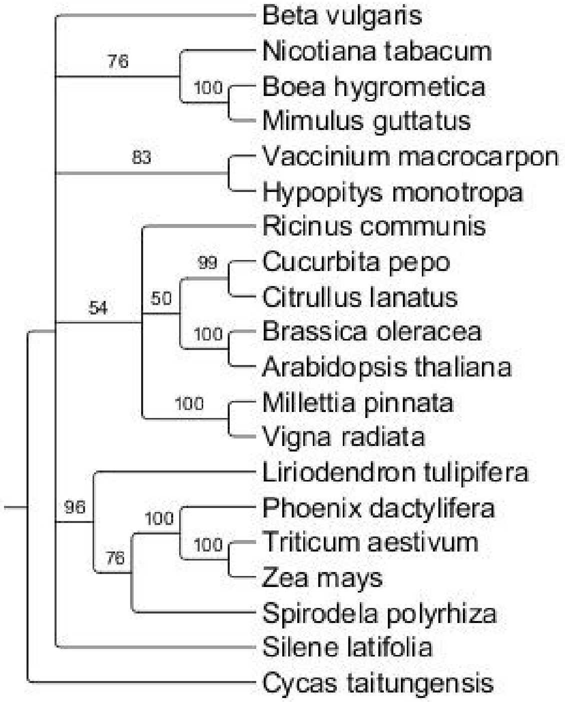
Beta vulgaris
76
Nicotiana tabacum
100
Boea hygrometica
Mimulus guttatus
83
Vaccinium macrocarpon
Hypopitys monotropa
Ricinus communis
99
Cucurbita pepo
Citrullus lanatus
54
50
100
Brassica oleracea
Arabidopsis thaliana
100
Millettia pinnata
Vigna radiata
Liriodendron tulipifera
96
100
Phoenix dactylifera
100
Triticum aestivum
76
Zea mays
Spirodela polyrhiza
Silene latifolia
Cycas taitungensis

**rps13**

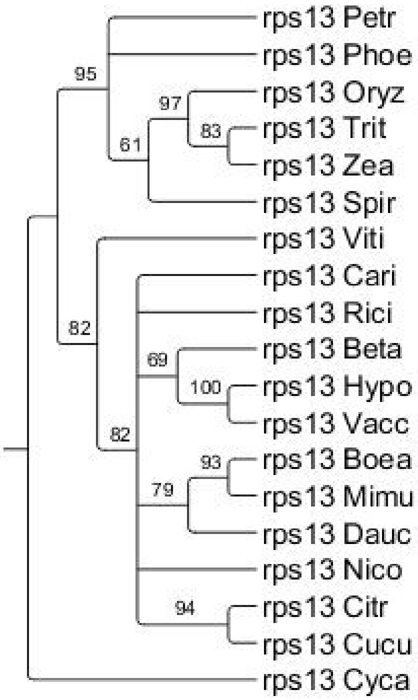
rps13 Petr
rps13 Phoe
95
97
rps13 Oryz
83
rps13 Trit
61
rps13 Zea
rps13 Spir
rps13 Viti
rps13 Cari
rps13 Rici
82
69
rps13 Beta
100
rps13 Hypo
rps13 Vacc
82
93
rps13 Boea
79
rps13 Mimu
rps13 Dauc
rps13 Nico
94
rps13 Citr
rps13 Cucu
rps13 Cyca
