## Supplementary Table S3 for "Mitochondrial genome of non-photosynthetic mycoheterotrophic plant *Hypopitys monotropa*,its structure, gene expression and RNA editing"

**Supplemental Table 3:**

**Plastid genes in the mitogenome of *H. monotropa.***

| **Gene** | **Gene length in the *Camellia sinensis* plastome (bp)*** | **Fragment length in the *H. monotropa* mitogenome (bp)** | **Fragment location (chromosome, bp)** | **Number of frameshifting indels** | **Number of nonsense mutations** |
| --- | --- | --- | --- | --- | --- |
|  | **protein coding genes** | | | | |
| *matK* | 1500 | 390 | linear chromosome, 409703–410092 | 0 | 0 |
| *ndhB*^1, 2^ | 777+756 | 484 | linear chromosome, 61755–62238 | 1 | 0 |
|  |  | 122 | linear chromosome, 368273–368394 | 1 | 0 |
| *ndhD* | 1515 | 177 | linear chromosome, 683982–684158 | 3 | 2 |
| *ndhJ* | 477 | 96 | linear chromosome, 383354–383449 | 0 | 2 |
| *psbA* | 1062 | 63 | linear chromosome, 301751–301813 | 0 | 0 |
| *psbC* | 1422 | 152 | linear chromosome, 178717–178868 | 2 | 0 |
| *rbcL* | 1428 | 117 | linear chromosome, 625016–625132 | 1 | 1 |
| *rpoB* | 3213 | 175 | linear chromosome, 547673–547847 | 1 | 0 |
| *rpoC1*^1^ | 453+1626 | 372 | linear chromosome, 297860–298231 | 0 | 2 |
| *rpoC2*^2^ | 4137 | 773 | circular chromosome, 2968–3740 | 0 | 0 |
|  |  | 925 | linear chromosome, 579856–580780 | 2 | 2 |
| *rps2* | 711 | 717 | circular chromosome, 3955–4671 | 2 | 1 |
| *rps4* | 606 | 309 | linear chromosome, 27980–28288 | 0 | 0 |
| *ycf2* | 6897 | 966 | circular, 43867–44832 | 3 | 2 |
| *ycf15* | 249 | 231 | circular, 43519–43749 | 0 | 1 |
|  | **RNA coding genes** | | | | |
| rrn23^2^ | 2809 | 50 | linear, 32069–32118 | - | - |
|  |  | 34 | linear, 222536–222569 |  |  |
| tRNA-Met (CAU) | 72 | 72 | linear, 621489–621560 | - | - |

* for intron-containing genes by "gene length" we mean the length of their CDSs

^1^ this gene has two exons in the plastid genome of *C. sinensis*. Their lengths are separated by the "+" sign.

^2^ there are two fragments of this gene in the mitochondrial genome of *H. monotropa*. Their features are described in separate lines.
