## Supplementary Table S5 for "Mitochondrial genome of non-photosynthetic mycoheterotrophic plant *Hypopitys monotropa*,its structure, gene expression and RNA editing"

**Supplemental Table 5:**

**Genes with stop codons that are introduced by RNA editing.**

| **Gene** | **Position** | **Length of original gene (CDS)** | **% of mapped transcriptomic reads** |
| --- | --- | --- | --- |
| *rpl2* | 127 | 1002 | 12.8% |
| *rps10* | 391 | 447 | 99.6% |
| *rps10* | 397 | 447 | 32.9% |
| *atp9* | 223 | 249 | 99.0% |
| *atp6* | 1162 | 1218 | 99.7% |
| *ccmFc* | 1315 | 1347 | 98.3% |
